# Modeling kidney development, disease, and plasticity with clonal expandable nephron progenitor cells and nephron organoids

**DOI:** 10.1101/2023.05.25.542343

**Authors:** Biao Huang, Zipeng Zeng, Hui Li, Zexu Li, Xi Chen, Jinjin Guo, Chennan C. Zhang, Megan E. Schreiber, Ariel C. Vonk, Tianyuan Xiang, Tadrushi Patel, Yidan Li, Riana K. Parvez, Balint Der, Jyun Hao Chen, Zhenqing Liu, Matthew E. Thornton, Brendan H. Grubbs, Yarui Diao, Yali Dou, Ksenia Gnedeva, Nils O. Lindström, Qilong Ying, Nuria M. Pastor-Soler, Teng Fei, Kenneth R. Hallows, Andrew P. McMahon, Zhongwei Li

**Affiliations:** USC/UKRO Kidney Research Center, Division of Nephrology and Hypertension, Department of Medicine, Keck School of Medicine, University of Southern California, Los Angeles, CA 90033, USA; Department of Stem Cell Biology and Regenerative Medicine, Keck School of Medicine, UXniversity of Southern California, Los Angeles, CA 90033, USA; College of Life and Health Sciences, Northeastern University, Shenyang 110819, P. R. China; Division of Maternal Fetal Medicine, Keck School of Medicine, University of Southern California, Los Angeles, CA 90033, USA; Department of Cell Biology, Duke University School of Medicine, Durham, NC 27710, USA; Department of Medicine, Department of Biochemistry and Molecular Medicine, University of Southern California, CA 90033, USA; Tina and Rick Caruso Department of Otolaryngology-Head and Neck Surgery, University of Southern California, Los Angeles, CA 90033, USA; Division of Stem Cell Biology Research, Department of Developmental and Stem Cell Biology, Beckman Research Institute of City of Hope, Duarte, CA 91010, USA; Lead contact; These authors contributed equally

## Abstract

Nephron progenitor cells (NPCs) self-renew and differentiate into nephrons, the functional units of the kidney. Here we report manipulation of p38 and YAP activity creates a synthetic niche that allows the long-term clonal expansion of primary mouse and human NPCs, and induced NPCs (iNPCs) from human pluripotent stem cells. Cultured iNPCs resemble closely primary human NPCs, generating nephron organoids with abundant distal convoluted tubule cells, which are not observed in published kidney organoids. The synthetic niche reprograms differentiated nephron cells into NPC state, recapitulating the plasticity of developing nephron *in vivo*. Scalability and ease of genome-editing in the cultured NPCs allow for genome-wide CRISPR screening, identifying novel genes associated with kidney development and disease. A rapid, efficient, and scalable organoid model for polycystic kidney disease was derived directly from genome-edited NPCs, and validated in drug screen. These technological platforms have broad applications to kidney development, disease, plasticity, and regeneration.

## INTRODUCTION

SIX2^+^ nephron progenitor cells (NPCs) play a central role in kidney organogenesis^1,2^. In the de-veloping kidney, niche signals coordinate two different NPC fates: some NPCs are induced to form nephrons, the functional units of the kidney, while others self-renew to repopulate the pro-genitor pool^3^. NPCs are then exhausted shortly after birth in the mice and before birth in humans, leaving limited regenerative potential in adult mammalian kidneys. Dysregulation of NPC fates underlies a number of congenital kidney diseases^4^ while uncontrolled proliferation of NPCs in Wilms tumor is the most prevalent pediatric kidney cancer^5^. Thus, a deeper insight into NPC bi-ology is central to improving an understanding of kidney development, congenital disease and cancer, and to applying developmental insight to regenerating kidney functions.

Over the past few years, we and others have developed systems to either generate NPCs *de novo* from pluripotent stem cells or expand NPCs from primary NPCs isolated from embry-onic/fetal kidneys such that: 1) mouse and human NPC-like cells can now be generated transi-ently using step-wise directed differentiation protocols from mouse and human pluripotent stem cells (hPSCs)^6–9^; 2) primary mouse and human NPCs can be isolated and expanded for a short period of time in two-dimensional (2D) culture format^10, 11^ or expanded long term over a few months in our previously reported three-dimensional (3D) culture format^12^. These systems have advanced our current understanding of NPC biology, and NPC-derived nephron organoids have been shown to be powerful tools in modeling kidney development and diseases^13^.

While acknowledging progress, significant limitations remain. First, current hPSC-derived nephron organoids fail to generate mature and functional kidney cell types and lack a diversity of critical cell types characteristic of distal nephron segments^13–16^, likely reflecting the quality of hPSC-derived NPC-like cells^16^. Second, compared to 2D culture, the currently available NPC 3D culture system is tedious and less compatible with functional genomics tools, such as CRISPR screens^17–19^, hindering genomic scale study of NPC biology. Third, despite attempts from us and others^10, 12, 20^, it has not been possible to expand NPCs derived from hPSCs, the desired cell source for kidney regeneration and disease modeling, over the long term.

Here, we report the development of chemically-defined 2D culture systems supporting the stable clonal expansion, over several months, of primary mouse and human NPCs, and hPSC-derived NPCs. In this system, NPCs maintain NPC properties and differentiate to generate distal cell types refractory to previous kidney organoid models. The novel NPC system facilitates ge-nome-scale genetic screening, disease modeling and provides a scalable platform for drug discovery.

## RESULTS

### p38 inhibition allows the derivation of clonal expandable NPC lines from any mouse strain

We previously developed a 3D culture system that can expand mouse NPCs (mNPCs) as clusters of cells in a chemically-defined culture medium, mNPSR^12^. However, mNPSR did not support mNPC expansion in a regular monolayer (2D) culture setting. To solve this problem, we have systematically screened the addition of a variety of small molecules and growth factors to mNPSR medium, resulting in mNPSR-v2: a chemically-defined medium that supports the derivation and long-term clonal expansion of mNPC lines from multiple mouse strains (**Fig. 1A**, **Fig. S1A** and **Tables S1 and S2**).

**Figure 1.**
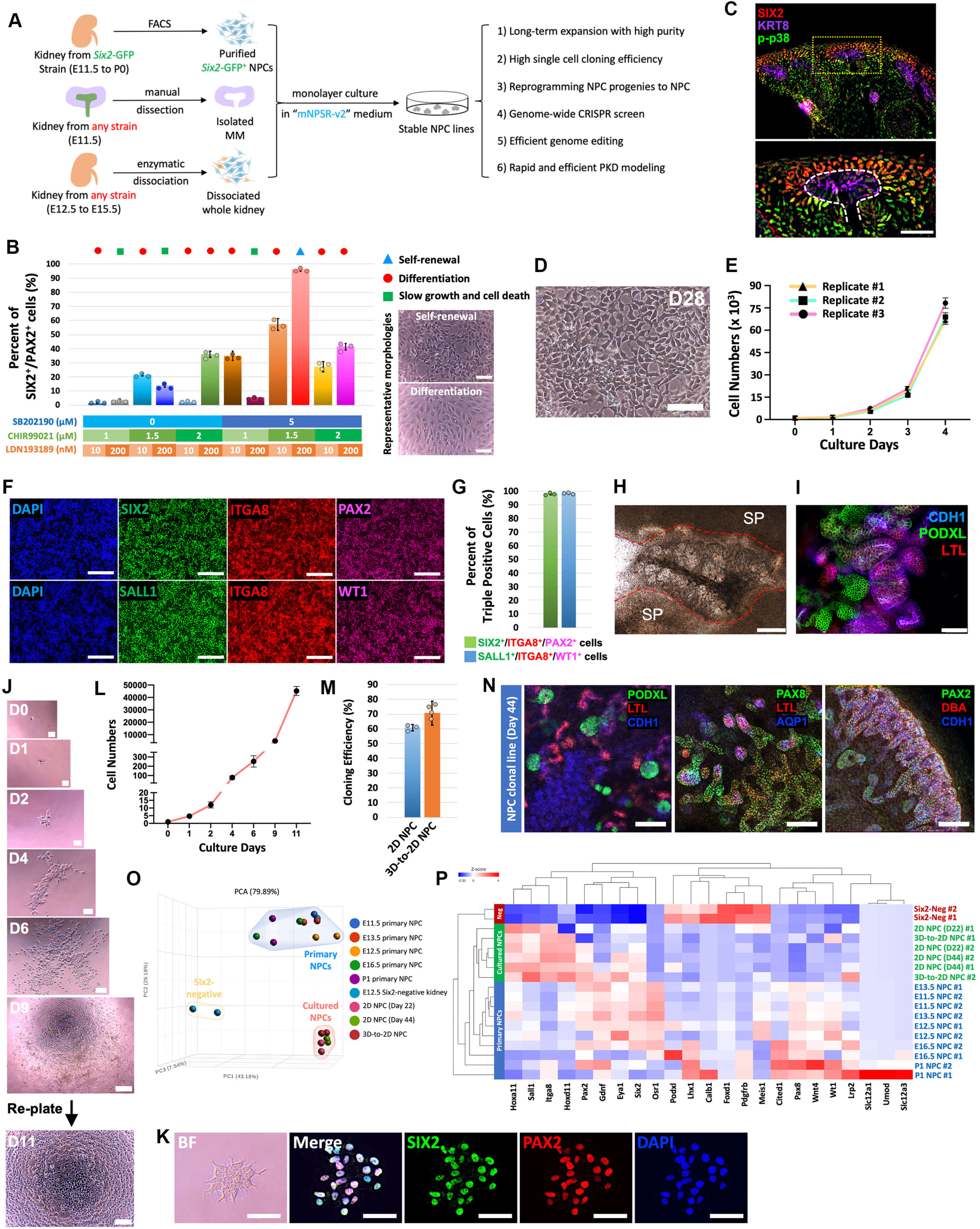
p38 inhibition allows the derivation of clonal expandable NPC lines from any mouse strain. (A) Schematic of mNPC line derivation and the applications. (B) Quantification of SIX2^+^/PAX2^+^ cell percentages in mNPCs after 4 days of culture in media containing different concentrations of CHIR99021, LDN193189, and SB202190 as indicated, on top of mNPSR medium (left panel). Typical bright field images of self-renewing or differentiating mNPCs are shown on the right panels. Scale bars, 50 μm. (C) Immunostaining of E13.5 mouse kidney section for phosphorylated p38 (p-p38), NPC marker SIX2, and ureteric epithelium marker KRT8. Scale bar, 50 μm. (D) Morphology of mNPCs cultured in mNPSR-v2 medium for 28 days (D28). Scale bar, 50 μm. (E) Growth curve of mNPCs cultured in mNPSR-v2 medium in a typical passage cycle starting from 5,000 cells. (F and G) Immunofluorescence staining (F) and quantification (G) of mNPCs cultured in mNPSR-v2 medium for 28 days for various NPC markers as indicated. Scale bars, 100 μm. (H) Bright-field image showing 7 days after co-culturing E12.5 spinal cord (SP) with aggregated mNPCs on Transwell filter. Red dashed lines indicate the boundaries between spinal cords and differentiated mNPCs. Scale bar, 200 μm. (I) Whole-mount immunofluorescence analysis of mNPC-derived nephron structures in (H) for marker genes representing different nephron segments. Scale bar, 100 μm. (J) Time-course bright-field images showing mNPC clonal expansion from one single cell. Scale bars, 50 μm. (K) Bright-field (BF) and immunofluorescence images of a representative single cell mNPC clone for NPC markers SIX2 and PAX2. Scale bars, 50 μm. (L) Growth curve of a single mNPC cultured in mNPSR-v2 medium over 11 days. (M) Single cell cloning efficiency of mNPCs derived in 2D culture format (2D NPC), or transitioned from existing 3D-cultured mNPCs to 2D culture (3D-to-2D NPC). (N) Whole-mount immunofluorescence analyses of nephron organoids generated from a clonal NPC line derived from parental mNPCs cultured in mNPSR-v2 for 44 days. Scale bars, 100 μm. (O) Three-dimensional presentation of principal component analysis (PCA) of bulk RNA-seq data of primary NPCs, NPCs cultured in mNPSR-v2 medium, and a negative control *Six2*-negative population from E12.5 kidney. (P) Heatmap showing gene expression of selected marker genes for undifferentiated NPCs and differentiated kidney cell types, in primary and cultured NPCs, based on bulk RNA-seq datasets. Data are presented as mean ± SD. Each column represents counts from three biological replicates (n=3). See also Figure S1, S2 and Tables S1-S4.

Compared to mNPSR, mNPSR-v2 has four additional small molecules: SB202190 (inhibitor of p38 MAPK), DAPT (inhibitor of Notch signaling), A83-01 (inhibitor of TGF-β signaling) and LDN193189 (inhibitor of BMP signaling), and required a different concentration (1.5 µM) of CHIR99021 (inhibitor of GSK3). Of these additional components, we have previously found that adding A83-01 and LDN193189 to mNPSR can enable the expansion of mNPCs at lower seeding mNPC numbers in a 3D culture setting^21^. LDN193189^10^ and DAPT^11^ have also been used in supporting short-term expansion of mNPCs. Consistently, we also noticed that adding DAPT prevented spontaneous differentiation from the 2D cultured mNPCs (**Fig. S1C and D**). The p38 MAPK inhibitor SB202190 has not been previously reported to support mNPC self-renewal. Interestingly, adding SB202190 appears to have the most significant effect in sustaining the percentage of SIX2^+^/PAX2^+^ mNPCs in culture (**Fig. 1B and S1B**). Importantly, phosphorylated p38 (p-p38), the active form of p38 MAPK, was not detected in primary mNPCs (**Fig. 1C**), indicating an intrinsic inhibition of p38 MAPK activity in the self-renewing mNPCs *in vivo*. mNPCs expanded in mNPSR-v2 stably proliferate with highly homogeneous morphology (**Fig. 1D and E**). After 4 weeks of culture and more than 10 passages, more than 95% of the cultured mNPCs are triple positive for SIX2/PAX2/ITGA8 and SALL1/WT1/ITGA8 (**Fig. 1F and G**). qRT-PCR showed the NPC marker genes, *Six2*, *Sall1*, *Pax2*, *Wt1* and *Gdnf,* were expressed at similar levels to those observed in primary NPCs (**Fig. S1E**). Upon withdrawing each individual medium component from mNPSR-v2, we confirmed that all components are essential to maintain the proper growth rate and undifferentiated state of the mNPCs (**Fig. S1F and G**).

We then utilized various assays to examine the nephrogenic potential of the cultured mNPCs *in vitro* and *in vivo*. When cultured mNPCs were aggregated and placed in close contact with the dorsal side of an isolated E12.5 spinal cord, a classic inducer of NPC development^6, 12^, we observed dramatic morphological changes, leading to the formation of numerous tubule-like structures after 7 days (**Fig. 1H**). Immunostaining further confirmed the formation of PODXL^+^ glomeruli, LTL^+^ proximal tubule, and CDH1^+^ distal tubule structures upon spinal cord induction (**Fig. 1I**). Nephron organoids were also formed from cultured mNPCs using our chemically-defined assay^12^, generating multiple segments of the nephron (**Fig. S1H**). When we reconstructed an engineered kidney from cultured mNPCs and cultured ureteric bud (UB)^22^, mNPCs induced dramatic branching morphogenesis from the UB (**Fig. S1I**), while the UB induced nephron formation from the mNPCs (**Fig. S1J**). Further, when mNPCs were transplanted onto the chicken chorioallantoic membrane (CAM) *in vivo*^23^, mNPCs differentiated into nephrons and chick vasculature infiltrated the transplant (**Fig. S1K-M, and Video S1**).

In strong contrast to the 3D-cultured mNPCs, which can be expanded only as clusters of cells^12, 21^, 2D-cultured mNPCs were efficiently expanded from single cells with a high cloning effi-ciency of 60–70% (**Fig. 1J-M**), with retention of NPC gene expression and nephrogenic potential for at least 2 months (**Fig. 1N, S2A and B**). To compare the global gene expression of cultured and primary mNPCs^12, 24^, we performed bulk RNA-seq (**Table S3**). Based on principal component analysis (PCA), cultured NPCs were clustered tightly together, with a clear separation of cultured and primary NPCs from the SIX2-negative non-NPCs along the PC1 axis (**Fig. 1O and S2C**). Importantly, cultured mNPCs largely overlapped with primary E11.5, E12.5, and E13.5 mNPCs in PC1 and PC3 axes (**Fig. S2D**). Consistently, cultured mNPCs were clustered with these early-stage mNPCs in the heatmap, using a selective group of typical NPC and nephron marker genes (**Fig. 1P**). These results suggest that cultured mNPCs likely represent early-stage (E11.5 to E13.5) mNPCs. Furthermore, the robustness of mNPSR-v2 allowed us to derive mNPC lines from iso-lated E11.5 metanephric mesenchyme (**Fig. S2E and F**) or from whole kidney cells of an early embryonic kidney (**>Fig. S2G-K**), thus enabling the derivation of mNPC lines from any mouse strain (**Fig. 1A and Table S4**).

### Capture the plasticity of developing nephron cells with mNPSR-v2 medium

While developing the mNPSR-v2 medium, we made an unexpected but interesting finding—a significant portion (10–20%) of *Six2*-GFP^-^ cells isolated from dissociated kidneys at all stages (E12.5, E14.5, E16.5 and P0 kidneys) adopted a SIX2^+^/SALL1^+^ phenotype within 4 days of culture (**Fig. 2A and B**). This finding suggests the possibility of phenotypic plasticity on the part of non-NPC type(s) in mNPSR-v2 medium. To verify this observation, we excluded first the possibility of potential contamination of a small number of *Six2*-GFP^+^ mNPCs in the *Six2*-GFP^-^ population dur-ing FACS. For this purpose, we confirmed the reported loss of SIX2^+^ NPCs in the postnatal day 3 (P3) kidney^1, 2^ (**Fig. S3A**) and cultured dissociated single P3, P4, P5 and P7 kidney cells in mNPSR-v2 for 4 days, and examined SIX2/SALL1 by immunostaining. Approximately 17% of the P3 kidney cells and 7% of the P4 kidney cells were scored as SIX2^+^/SALL1^+^, rare SIX2^+^/SALL1^+^ cells were observed also within P5 and P7 cultures (**Fig. 2C and D, and S3B and C**). These results support the contention that a non-NPC-like cell-type generates SIX2^+^/SALL1^+^ NPC-like cells under mNPSR-v2 condition. The reprogramming is most efficient in mNPSR-v2 (18%), less efficient in mNPSR (6%), and does not happen in basal medium with 10% FBS (0%) (**Fig. S3D and E**), suggesting that, in addition to the intrinsic plasticity of the cells, proper extrinsic signals are also needed for successful reprogramming. To further characterize the induced *Six2*^+^ cells, we developed an assay to purify these cells using an NPC lineage tracing reporter system^3^ (**Fig. 2E**). P3 kidneys were isolated from *Six2*-GFP-CreERT2; *Rosa26*-tdTomato (*Six2*-tdT for short) mice, dissociated into single cells, and cultured in mNPSR-v2. Three days after culture, tamoxifen was added, which allows the permanent labeling of tdTomato (tdT) in cells that express *Six2*. After another 24 hours (on day 4 of culture), similar to immunostaining results, around 20% of the cultured P3 kidney cells became *Six2*-tdT^+^ as shown by FACS (**Fig. S3F and G**). Upon additional culture to day 8, these *Six2*-tdT^+^ cells showed a more complete NPC profile expressing *Six2*, *Pax2*, *Sall1*, *Wt1*, *Gdnf*, *Hoxd11*, and *Eya1*, coinciding with loss of expression of nephron marker genes *Slc12a1*, *Slc12a3*, and *Aqp1* (**Fig. 2F**). Taken together, these data suggest that some kidney cell types retain a developmental plasticity to convert to NPCs for a limited period following the cessation of active nephrogenesis, and these cells were reprogrammed to NPC-like cells in mNPSR-v2 culture.

**Figure 2.**
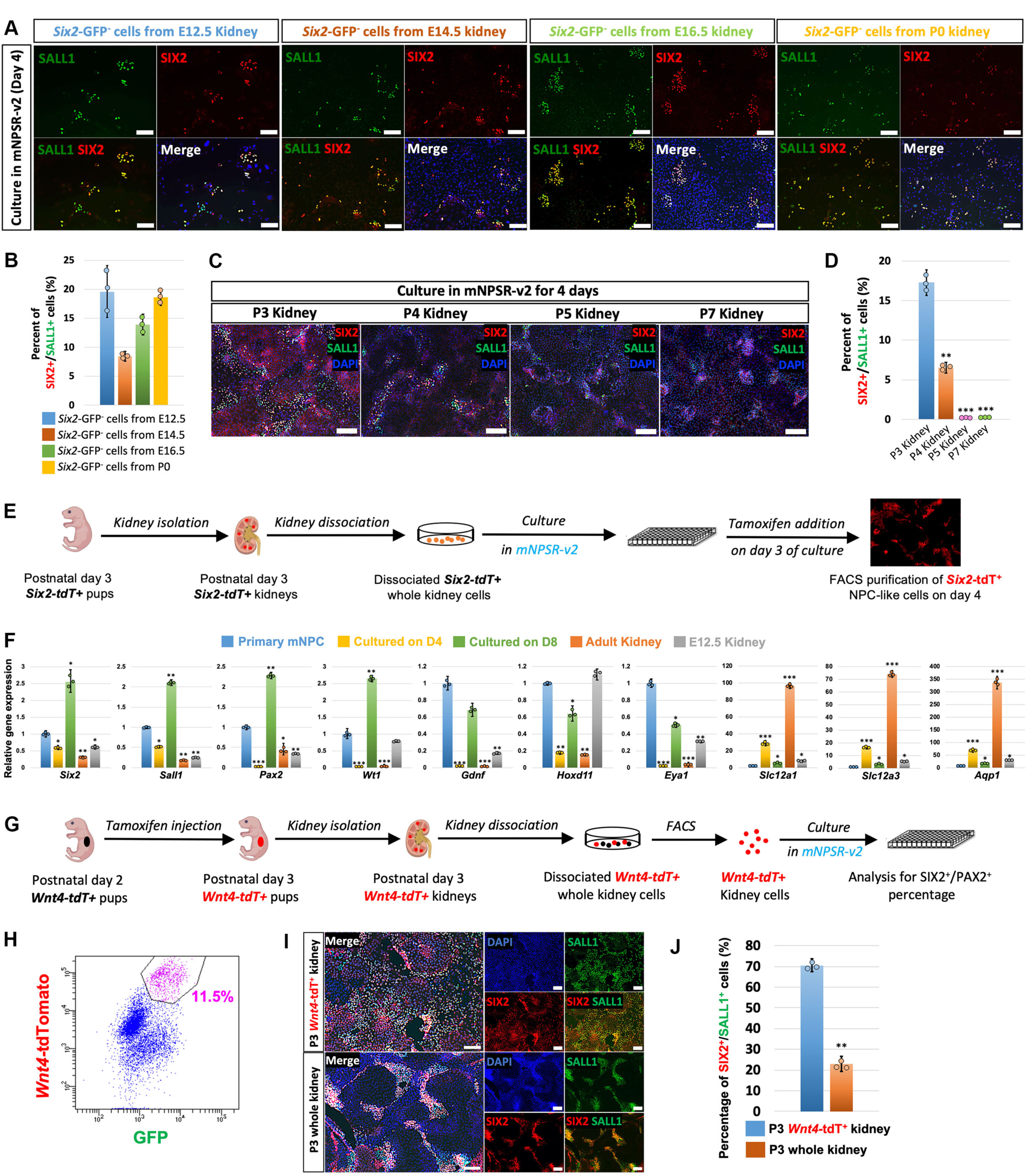
Capture the plasticity of developing nephron cells with mNPSR-v2 medium. (A and B) Immunofluorescence images (A) and quantification (B) of the expression of NPC marker genes SIX2 and SALL1, in *Six2-*GFP^-^ cells isolated from E12.5, E14.5, E16.5, and P0 kidneys and cultured in mNPSR-v2 medium for 4 days. Scale bars, 100 μm. (C and D) Immunofluorescence images (C) and quantification (D) of the expression of SIX2 and SALL1 in P3, P4, P5, and P7 whole kidney cells cultured in mNPSR-v2 medium for 4 days. Note that only fluorescence signals in the nucleus were true SIX2 signals. Membrane-bound signals were from the non-specific binding of the SIX2 primary antibody. Scale bars, 100 μm. (E) Schematic showing the genetic labeling and FACS isolation of the induced *Six2-*tdT^+^ cells from P3 kidney cells cultured in mNPSR-v2 for 4 days. (F) qRT-PCR analysis of primary mNPC, *Six2-tdT*^+^ cells cultured for 4 days and 8 days, adult kidney, and E12.5 kidney, for various marker genes of NPC and differentiated nephron cell types. (G) Schematic showing the genetic labeling, FACS isolation, and culturing of *Wnt4-*tdT^+^ cells from P3 kidneys with mNPSR-v2 medium. (H) Flow cytometry gating plot showing the purification of *Wnt4*-tdT^+^ cells from P3 kidneys. (I and J) Immunofluorescence images (I) and quantification (J) of the expression of SIX2 and SALL1 in *Wnt4*-tdT^+^ kidney cells isolated from P3 kidney, and in whole P3 kidney cells, cultured in mNPSR-v2 medium for 4 days. Scale bars, 100 μm. Data are presented as mean ± SD. Each column represents counts from three biological replicates (n=3). The significance was determined by two-tailed unpaired Student’s t tests; ns, not significant; *, p<0.05; **, p<0.01; ***, p<0.001. See also Figure S3.

*Wnt4*+ cells, the immediate differentiated progenies of the NPCs, can migrate back to the cap mesenchyme niche, where NPCs reside, and reverse back to the NPC state *in vivo*^25^. To investigate whether *Wnt4*+ cells might also contribute to the plasticity we observed *in vitro*, we injected tamoxifen into P2 *Wnt4*-GFP-Cre-ERT2; *Rosa26*-tdTomato (*Wnt4*-tdT for short) mice to permanently label *Wnt4*^+^ cells and their progeny. Kidneys were isolated 24hrs later on P3, and tdTomato^+^ cells FACS purified and cultured in mNPSR-v2 (**Fig. 2G and H**). After 4 days of culture, more than 70% of the cultured *Wnt4*-tdT+ cells stained positive for SIX2/SALL1, as compared to 20% starting from P3 whole kidneys (**Fig. 2I and J**). These results suggest that, like *in vivo*, the *Wnt4*+ cells represent a highly plastic cell population that can be efficiently reprogrammed to an NPC state *in vitro*. It also further strengthened the notion that the mNPSR-v2 medium represents a *bona fide* synthetic niche that allows NPC self-renewal and *Wnt4*+ cells lineage reprogramming as *in vivo*.

### Genome-wide CRISPR screen in the NPC lines identify functional genes for kidney devel-opment and disease

The ease of generating CRISPR/Cas9 sgRNAs has afforded the development of CRISPR technologies for pooled genome-wide screens^17–19^, enabling the interrogation of functional genes in the whole genome for the discovery of novel insights into biological processes and the identification of novel drug targets and therapies for diseases^26^.

Considering the central roles NPCs play in kidney organogenesis, Wilms tumor, and congenital kidney diseases such as congenital anomalies of the kidney and urinary tract (CAKUT), the use of CRISPR screen tools to understand NPC biology from the genome-wide perspective could provide valuable molecular and genetic insight into these areas. Thus, as a proof-of-concept, we introduced a genome-wide CRISPR knockout library^27^ into cultured NPCs to screen for genes essential for NPC fitness (**Fig. 3A**). Abundance of sgRNAs in the cultured NPCs were determined by next-generation sequencing 3 weeks after CRISPR library was introduced (**Tables S5**). Compared to sgRNA abundance in the original CRISPR screen library, decrease of sgRNA abundance indicates that expressing the sgRNA results in gene knockout that reduces cell fitness (e.g., enhanced NPC apoptosis or reduced NPC proliferation), while conversely, increase of sgRNA abundance indicates increased fitness upon gene removal (e.g. blocked apoptosis or differentiation, or enhanced NPC proliferation). With the MAGeCK-VISPR tool^28^, sgRNA abundance changes from 4 different sgRNAs targeting the same gene in the library are statistically integrated as beta scores (**Table S6**). Positively-selected (i.e. increased sgRNA abundance) genes with more dra-matic sgRNA abundance increase will have higher positive beta scores; while negatively-selected genes with more dramatic sgRNA abundance decrease appear as lower negative beta scores.

**Figure 3.**
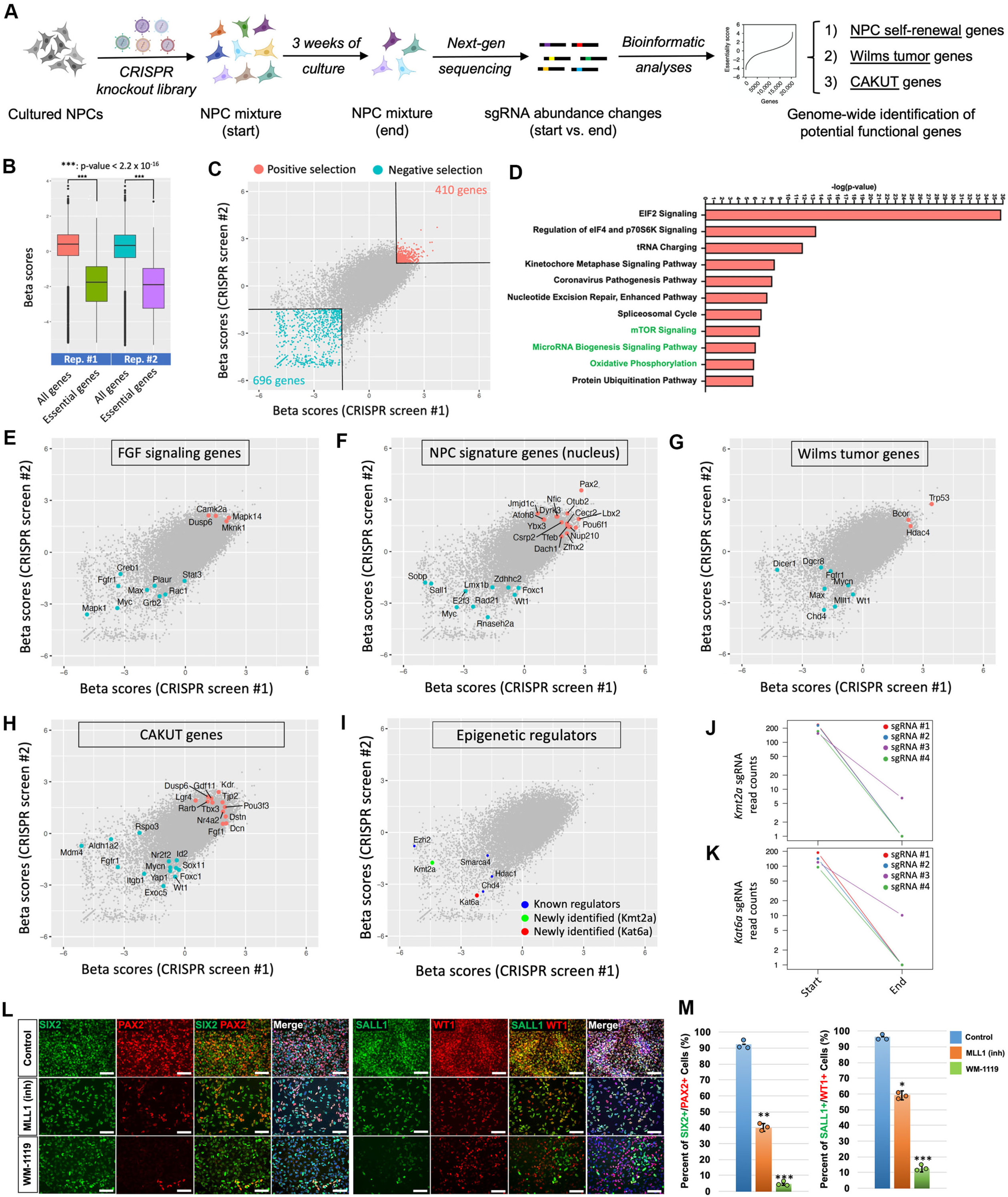
Genome-wide CRISPR screen in the NPC lines identify functional genes for kidney development and disease. (A) Schematic diagram illustrates the workflow of genome-wide CRISPR knockout screen in mNPC lines. (B) Box plot showing the distribution of beta scores of all genes or essential genes in two CRISPR screen replicates. Boxes, 25th to 75th percentiles; whiskers, 1st to 99th percentiles. (C) MAGeCKFlute scatterplots of beta scores showing common positively and negatively selected genes from two CRISPR screen replicates. (D) Top 11 enriched ingenuity pathway analysis (IPA) Canonical Pathways from CRISPR screen replicate #1. (E–I) MAGeCKFlute scatterplots of beta scores from two CRISPR screen replicates showing FGF signaling related genes (E), NPC signature genes that encode nucleus-localizing proteins (F), Wilms tumor-related genes (G), CAKUT-related genes (H), and genes that regulate epigenetic mechanisms (I). (J and K) Normalized read counts of 4 individual sgRNAs targeting *Kmt2a* (J) and *Kat6a* (K) at the start and the end of the CIRPSR screen. (L and M) Immunofluorescence analyses (L) and quantification (M) of the expression of SIX2, PAX2, SALL1, and WT1 in mNPCs after treated with KMT2A inhibitor MLL1 (inh), or KAT6A inhibitor WM-1119, for 8 days. Scale bars, 50 μm. Data are presented as mean ± SD. Each column represents counts from three biological replicates (n=3). The significance was determined by two-tailed unpaired Student’s t tests; ns, not significant; *, p<0.05; **, p<0.01; ***, p<0.001. See also Figure S4 and Tables S5-11.

Two biological replicates of this CRISPR screen showed high similarity overall (**Fig. S4A**). As expected, significant global changes of sgRNA abundance were observed after 3 weeks of the screen, with similar abundance for all sgRNAs in the library before the screen, and dispersed distribution of the abundance of different sgRNAs after the screen (**Fig. S4B and S4C**). As ex-pected, “pan-essential genes” in CRISPR knockout screens which include ribosomal and cell cy-cle regulatory genes^29^, underwent significant negative selection (**Fig. 3B**). Statistical analysis (beta scores >1.5 or <-1.5) identified 410 positively-selected genes and 696 negatively-selected genes shared between replicate assays (**Fig. 3C**).

Next, we used Ingenuity Pathway Analysis (IPA) to analyze identified gene sets (**Table S7**). Consistent with the large number of essential genes identified in this type of screen, funda-mental cellular pathways necessary for cell survival, such as basic transcription and translation machineries, were found to be at the top of the enriched pathways from IPA’s Canonical Pathways analysis. Importantly, mTOR signaling, microRNA biogenesis, and oxidative phosphorylation pathways, which play critical roles in NPC self-renewal in *in vivo* studies^30–34^, were all identified among the top enriched pathways in both replicates (**Fig. 3D, S4D and E, and Table S8**). Con-sistent with the essential role of FGF signaling in NPC self-renewal *in vitro* (**Fig. S1F and G**) and *in vivo*^1, 2^, *Fgfr1*, *Grb2*, *Mapk1*, and *Myc*, genes encoding critical FGF receptor, signaling mediator, and effector proteins, were among the top negatively-selected genes (**Fig. 3E, and Table S9**). On the contrary, *Dusp6*, which encodes a phosphorylase that inhibits FGF signaling in a feedback loop, was a top positively-selected gene (**Fig. 3E**). *Mapk14*, which encodes p38 MAPK, was among the top positively-selected genes, consistent with our observation that p38 inhibition is essential for NPC self-renewal (**Fig. 1B and C, S1B, F and G**). Similar patterns were evident for Wnt and LIF signaling pathway components (**Fig. S4F and G, and Table S9**). These results fur-ther validated the high quality of our CRISPR screen results.

We next set out to systematically identify potential NPC self-renewal genes in the whole genome from our CRISPR screen datasets. To further narrow down functional genes, we cross-referenced our previously published bulk RNA-seq datasets for primary NPCs from various de-velopmental stages^12, 24^. First, we removed essential genes from the CRISPR screen hits. Second, we removed the genes that showed low gene expression levels in the primary NPCs. Third, we overlapped our CRISPR screen identified gene list with two NPC-enriched gene lists—E12.5 *Six2*^+^ NPC vs. *Six2*^-^ non-NPC^12^ and E16.5 NPC vs. IPC (interstitial progenitor cell)^24^. With this stringent data filtration method, a short-list of 64 genes were obtained that showed consistent results from both screen replicates (**Table S10**). Interestingly, 25 out of the 64 genes encode proteins that localize to the nucleus (**Fig. 3F**). Among these genes, we identified *Sall1*^35–37^, *Wt1*^38^, *Foxc1*^39^, and *Myc*^40, 41^ as top negatively-selected genes, which are well-established transcription factors for maintaining NPC identity *in vivo*. Genes with previously unknown functions in NPC self-renewal were also identified. As an example, *Sobp* is a transcription factor known to regulate cochlear development^42^. It is also expressed in the NPCs^12, 24^, with its expression dramatically decreased in P0 NPCs, as compared to NPCs at earlier developmental stages. A recent study reported that, *Sobp*, as a co-factor for *Six1*, interferes with the transcriptional activation of *Six1*/*Eya1* target genes during craniofacial development, likely leading to Branchio-oto-renal syn-drome (BOR)^43^. These observations, together with our CRISPR screen result, support a role for *Sobp* in the regulation of NPC fates.

Wilms tumor, the most prevalent pediatric kidney cancer, is associated with the retention and expansion of cells that have overlapping signatures between NPCs and their early nephron-committed descendants. Currently, around 30 Wilms tumor-related genes have been well estab-lished^5^. Of note, 11 out of the 30 genes (36.7%) were identified by our CRISPR screen. The tumor suppressor genes *Trp53* and *Bcor* were among the top positively selected genes, while onco-genes such *Mycn* and *Max* were amongst the negatively selected gene set (**Fig. 3G, and Table S11**). Dysregulation of normal kidney development process can also lead to congenital kidney diseases, such as CAKUT, for which around 330 genes might be related^4^. Out of the 330 genes, 25 (7.6%), were identified in our CRISPR screen, consistent with the notion that dysregulation of NPC fates represents a significant source for kidney malformation (**Fig. 3H and Table S11**). Con-firmation of the large numbers of known Wilms tumor and CAKUT related genes suggests that our CRISPR screen datasets might identify other genes with previously unknown functions in these diseases (**Table S7**).

Epigenetic mechanisms have been found to play critical roles in stem/progenitor cell fate decisions^44, 45^, including NPC self-renewal *in vivo*^46^. Our CRISPR screen has identified the majority of these reported epigenetic factors, including *Hdac1*^47^, *Chd4*^48^, *Ezh2*^49^, and *Smarca4*^50^. In addi-tion, two other epigenetic regulators, *Kmt2a* (*Mll1*) and *Kat6a*, were found to be top negatively-selected genes, whose function has not been examined in NPCs (**Fig. 3I-K**). Interestingly, recent whole-exome sequencing in families with CAKUT identified dominant monogenic point mutations in human *KMT2D* and *KAT6B* genes to cause syndromic CAKUT^51^. Thus, we were intrigued to validate experimentally whether perturbations of *Kmt2a* and *Kat6a* in mouse NPCs can lead to dysregulation of NPC fates. mNPCs treated with two small molecule inhibitors to KMT2A, MLL1 (inh), and WDR5 degrader^52^, and two small molecule inhibitors to KAT6A, WM-1119, and MOZ-IN-3, showed significantly reduced proliferation, and decreased expression of NPC marker genes SIX2, PAX2, SALL1, and WT1, linking KMT2A and KAT6A activities to the maintenance of NPC identity (**Fig. 3L and M, and S4H and I**). These findings suggest mutations in *KMT2D* and *KAT6B* might dysregulate NPC programs in association with human CAKUT .

Taken together, our CRISPR screen datasets (**Tables S5-S11**) provide valuable genome-scale resources for future studies of kidney development, Wilms tumor, and CAKUT.

### Rapid, efficient, and scalable PKD modeling directly from genome-edited mouse NPCs

The long-term clonal expansion ability of the cultured NPCs, coupled with the ease of genome-editing in 2D culture, encouraged us to explore genetic modeling of kidney disease. Autosomal-dominant polycystic kidney disease (ADPKD) is the most prevalent inherited kidney disease oc-curring at a frequency of approximately 1 in 500 live births^53^. ADPKD patients develop large cysts in their kidneys, which can frequently lead to end-stage kidney disease. There are limited effective therapeutic options for patients with ADPKD. Studies from several groups^54–58^ have successfully modeled ADPKD through the genetic removal of *PKD1* or *PKD2* in human pluripotent stem cells (hPSCs), and PSC differentiation to cyst-forming kidney organoids, under a variety of conditions.

We designed a one-vector multiplexed CRISPR-Cas9 system, comprising three sgRNA expression cassettes cloned in tandem into a basic lentiviral vector expressing a puromycin re-sistance gene, for highly efficient one-step gene knockout directly in NPCs (**Fig. 4A**). The three sgRNAs target the same gene with sgRNA targeting sequences 50-100 bp apart from each other for increased gene knockout efficiency^59^. Using a mouse NPC line constitutively expressing Cas9 and GFP (Cas9-GFP)^60^ (**Table S4**), targeting with lentivirus expressing multiplexed sgRNAs against *EGFP* led to near complete loss of EGFP fluorescence in NPCs (**Fig. 4B and C**).

**Figure 4.**
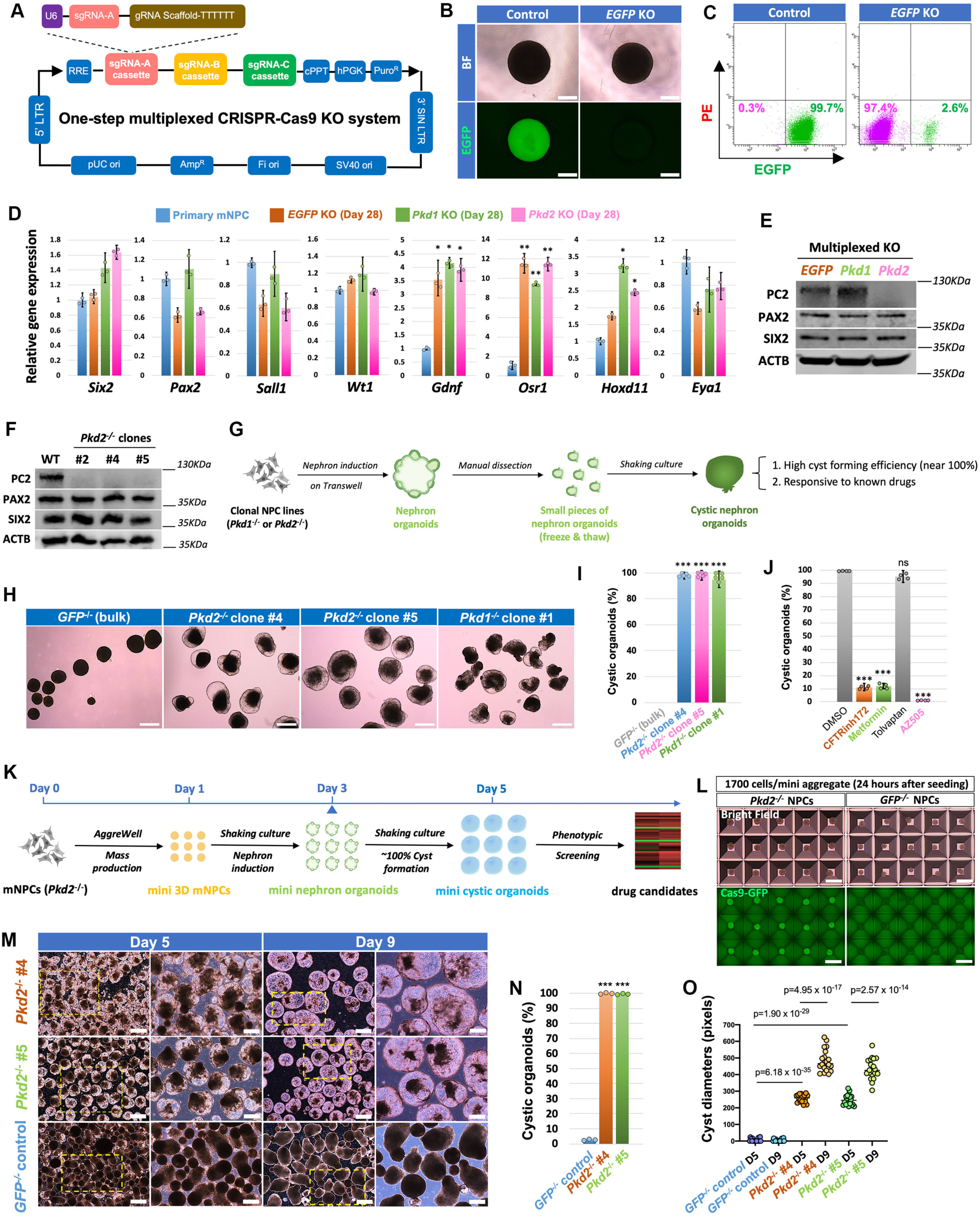
Rapid, efficient, and scalable PKD modeling from genome-edited mouse NPCs. (A) Schematic showing the lentiviral vector design for one-step multiplexed CRISPR-Cas9 knockout (KO). (B) Bright-field and EGFP images of aggregated *Cas9*-GFP mNPCs 7 days upon infection of lentivirus targeting *EGFP* gene and puromycin selection. Empty lentiviral vector without sgRNA was used as control. Scale bars, 200 μm. (C) Flow cytometry analysis of EGFP expression in control (left) or *EGFP* KO (right) *Cas9-*GFP mNPCs. (D) qRT-PCR analysis for various NPC marker genes as indicated in bulk *EGFP^-/-^*, *Pkd1^-/-^*, and *Pkd2^-/-^* NPC lines 28 days after lentiviral infection and puromycin selection. Primary NPC sample was used as control. (E) Western blot analysis of bulk *EGFP^-/-^*, *Pkd1^-/-^*, and *Pkd2^-/-^* mNPC lines (day 28 of culture after lentiviral infection) for PC2, PAX2, and SIX2, and beta-actin (ACTB) expression. (F) Western blot analysis of *Pkd2^-/-^* clonal mNPC lines for PC2, PAX2, and SIX2, and beta-actin (ACTB) expression. Wild-type (WT) mNPC line without genome-editing, cultured for a similar time period, was used as control. (G) Schematic of the experimental protocol for deriving cystic nephron organoid model from *Pkd1^-/-^* or *Pkd2^-/-^* clonal mNPC lines. (H) Bright-field images showing cyst formation in nephron organoids derived from *Pkd1^-/-^* or *Pkd2^-/-^* clonal mNPC lines. mNPC line with *EGFP* KO in bulk was used as control. Scale bars, 200 μm. (I) Quantification of the percentages of cystic organoids in samples shown in (H). (J) Percentages of cystic organoids in *Pkd2^-/-^* clonal mNPC line-derived nephron organoids treated with DMSO (control) or various drugs as indicated. (K) Schematic of the experimental protocol for generating scalable mini cystic nephron organoids from *Pkd2^-/-^*clonal mNPC lines for high-throughput drug screening. (L) Bright-field and GFP images showing mini 3D mNPC aggregates upon seeding onto Aggrewell plates overnight, from *EGFP^-/-^*or *PKD2^-/-^ Cas9-*EGFP genetic backgrounds. Scale bars, 500 μm. (M) Bright-field images showing cyst growth at 5 days and 9 days after shaking culture using protocol described in (K). Scale bars under each timepoint, 500 μm (left panels) and 200 μm (right panels). (N and O) Quantification of cystic organoid percentages (N) and cyst diameters (O) for samples from (M). Data are presented as mean ± SD. Each column represents counts from three biological replicates (n=3). The significance was determined by two-tailed unpaired Student’s t tests; ns, not significant; *, p<0.05; **, p<0.01; ***, p<0.001. See also Figures S5 and S6.

We then generated *Pkd1* or *Pkd2* knockout NPCs using this system. All the genome-ed-ited mNPC lines showed stable expression of NPC marker genes at similar levels to that of pri-mary NPCs (**Fig. 4D**). Immunoblot analysis of the bulk *Pkd2*^-/-^ mNPCs confirmed that polycystin 2 (PC2, the protein product of *Pkd2*) was completely depleted (**Fig. 4E**). We next isolated single cell clones from *Pkd1* or *Pkd2* bulk knockout mNPCs (**Fig. S5A**) and randomly selected *Pkd2^-/-^* clones #2, #4, and #5 and *Pkd1^-/-^* clone #1 for further studies. All these clonal NPC lines showed stable expression of NPC marker genes similar to primary NPCs (**Fig. S5B**). Targeted PCR flank-ing the sgRNA targeting areas of the genome indicated deletions around the genomic loci (**Fig. S5D**). Sanger sequencing confirmed genome editing resulted in premature termination of gene transcription (**Fig. S5C**), and immunoblotting confirming the complete depletion of PC2 in all the clonal *Pkd2*^-/-^ mNPC lines (**Fig. 4F**).

To examine cyst formation, we used our published protocol to make nephron organoids at the air-liquid-interface in Transwells^12^, then manually dissected organoids into smaller pieces which were transferred to shaking culture^54^ (**Fig. 4G).** Almost all *Pkd1*^-/-^ and *Pkd2*^-/-^ NPC-derived nephron organoids formed cysts of different sizes, whereas no cysts were formed in *EGFP*^-/-^ con-trol organoids (**Fig. 4H and I**). Cysts formed at similar efficiency following a freeze-and-thaw cycle, opening up opportunities for easy transportation and distribution of ready-to-use PKD organoids (**Fig. S5E-G**).

Treatment with CFTRinh172^61^, metformin^62, 63^, and AZ505^64, 65^, previously reported to have cyst growth suppressing activities, decreased the percentage of organoids undergoing cyst formation and cyst diameter within the organoids (**Fig. 4J, S5H and I**). As expected, these PKD organoids did not respond to tolvaptan, which is specific for cystic cells of ureteric progenitor cell-derived collecting duct epithelium^66, 67^. Next, to establish a scalable PKD organoid model, we used commercially available AggreWell plates to mass produce mini 3D NPC aggregates (1700 NPCs per mini aggregate, making ∼1500 mini aggregates in each well of a 6-well plate). After aggregation, the mini NPC aggregates were transferred into shaking culture, the setting for nephron induction and cyst formation (**Fig. 4K and L**).

Under the optimized culture condition (**Fig. S6A-D, and Methods**), *Pkd1*^-/-^ or *Pkd2^-^*^/-^ NPC aggregates differentiated into cystic nephron organoids under shaking culture in a synchronized manner with the budding of cysts as early as 4 days after shaking culture (**Fig. S5J**). This method allows a more rapid cystic organoid generation than current PKD organoids starting from hPSCs, which require at least 3 weeks to form cysts^54–58^. In our system, the cysts continued to grow in size over the next few days, at which point 100% of the observed mini nephron organoids formed cysts with similar diameters (**Fig. 4M-O**). Thus, a rapid, efficient, and scalable organoid model for PKD is established directly from genetically modified NPCs.

### A drug screen identifies PTC-209 as a potential drug candidate for PKD treatment

We further determined whether the NPC-derived PKD organoids recapitulates key molecular, cellular, and metabolic features of PKD. Whole-mount immunostaining of the *Pkd2*^-/-^ cystic organoids clearly showed the co-expression of LTL and CDH1 in the majority of the cyst-lining cells (**Fig. 5A**). Phosphorylated histone H3 (pHH3) staining indicated a dramatic elevation of proliferative cells in the cyst-lining epithelial cells of the *Pkd2*^-/-^ organoids, as compared to the epithelial cells in the control organoids (**Fig. 5A and B**). Consistently, starting from the same number of NPCs, *Pkd2* knockout cystic organoids have significantly higher total genomic DNA (**Fig. S7A**), suggesting increased cellular proliferation. At the molecular level, gene expression analysis clearly showed enhanced expression of a group of cell cycle genes that promote cell proliferation in the *Pkd2*^-/-^ organoid as compared to *GFP*^-/-^ organoids (**Fig. 5C**), and mTOR pathway^68, 69^ and MYC^70, 71^ activity were significantly activated in these cystic organoids (**Fig. 5D and E**). Comparison of gene expression in the cystic and non-cystic portions from the *Pkd2*^-/-^ organoids further confirmed that it is the cystic portions contributed to the dramatic gene expression changes (**Fig. S7B-E**).

**Figure 5.**
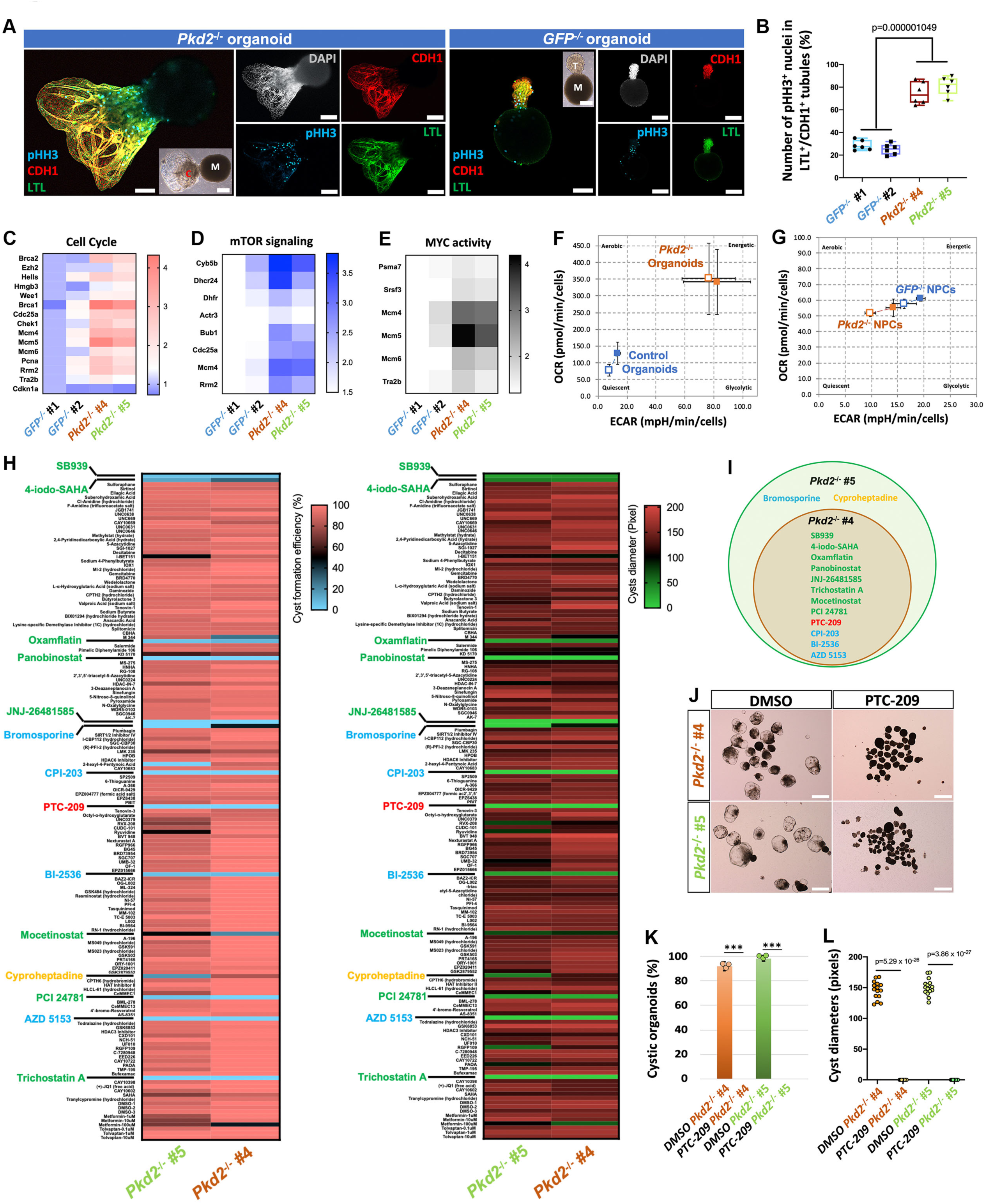
Drug screening identifies PTC-209 as a potential candidate for PKD treatment. (A) Immunofluorescence analysis of *Pkd2^-/-^* or *GFP^-/-^* mini nephron organoids for pHH3, LTL and CDH1. C, cystic portion; M, mesenchymal portion; T, tubule portion. Scale bars, 100 μm. (B) Quantification of the numbers of pHH3^+^ nuclei in LTL^+^/CDH^+^ cells from *Pkd2^-/-^* or *GFP^-/-^* mini nephron organoids. (C–E) Heat map presentation of cell cycle (C), mTOR signaling (D), and MYC activity (E) associated gene expression levels as determined by qRT-PCR, in *Pkd2^-/-^*or *GFP^-/-^* mini nephron organoids. (F and G) Metabolic analyses of OCR and ECAR in *Pkd2^-/-^* or *EGFP^-/-^* mini nephron organoids (F) and *Pkd2^-/-^*or *EGFP^-/-^* NPCs (G) using Seahorse assays. Blank boxes indicate baseline levels and filled boxes indicate stressed levels upon oligo/FCCP treatment. (H) Heat maps showing the quantification of drug screening results in terms of cyst formation efficiency (left) and cyst diameter (right) using mini PKD organoid models derived from two clonal *Pkd2^-/-^* mNPC lines #4 and #5. Identified HDAC inhibitor hits are shown in green, and BRD4 inhibitors are shown in blue. (I) Venn diagram showing the common drug candidates identified through drug screening from *Pkd2^-/-^* #4 and #5 mNPC line-derived mini PKD organoids. (J) Bright-field images showing mini PKD organoids derived from *Pkd2^-/-^* #4 and #5 NPC lines, treated with DMSO (control) and PTC-209, during the drug screening. Scale bars, 500 μm. (K and L) Quantification of cyst organoid percentages (K) and cyst diameter (L) in samples shown in (J). Data are presented as mean ± SD. Each column represents counts from three biological replicates (n=3). The significance was determined by two-tailed unpaired Student’s t tests; ns, not significant; *, p<0.05; **, p<0.01; ***, p<0.001. See also Figures S7 and S8.

We then interrogated the metabolic status of the *Pkd2*^-/-^ organoids and the *GFP*^-/-^ organ-oids using SeahorseXFp^TM^ analysis. Both the oxygen consumption rate (OCR, indicator of oxidative phosphorylation) and the extracellular acidification rate (ECAR, indicator of glycolysis) were dramatically higher in the *Pkd2*^-/-^ organoids as compared to control organoids (**Fig. 5F, S7F-I, N, and P**), consistent with the observed metabolic reprogramming in PKD in which the cyst-lining cells use significantly more energy from glycolysis to support the high metabolic needs for cyst growth^72, 73^. No further OCR and ECAR increases were observed in the *Pkd2*^-/-^ organoids upon Oligomycin/FCCP treatment to metabolically stress the organoids, suggesting that cystic cells in PKD organoid culture engaged maximal energy consumption in support of rapid growth (**Fig. 5F, S7F-I, N, and P**). This is in contrast to the control *GFP^-/-^* organoids, which showed a capacity to increase energy consumption through both OCR and ECAR by 50-100% from baseline in re-sponse to stress (**Fig. S7N**). Importantly, comparing undifferentiated NPCs with organoids indi-cated the metabolic phenotypes following *Pkd2* removal were specific to differentiated epithelial cells but not progenitor cells (**Fig. 5G, S7J-M, O, and P**). Moreover, although the oxidative (OCR) to glycolytic (ECAR) metabolism ratio was unchanged in *Pkd2^-/-^* NPCs relative to *GFP^-/-^*NPCs, the OCR/ECAR ratio was significantly elevated in control *GFP^-/-^* organoids relative to that in *Pkd2^-/-^* organoids (**Fig. S7P**). Thus, *Pkd2^-/-^* organoids had reduced oxidative relative to glycolytic capac-ity, which is consistent with prior reports demonstrating impaired mitochondrial oxidative function in ADPKD cells and tissues lacking polycystin function^74, 75^. Taken together, the cultured NPC-derived PKD organoid models recapitulate important molecular, cellular, and metabolic features of ADPKD pathogenesis. Lastly, as proof-of-concept, we established a similar PKD organoid model from a wild-type NPC line without endogenous CAS9 (**Fig. S8A-E**), opening new avenues for modeling PKD from existing transgenic mouse strains that are of interest to PKD researchers. We next set out to perform a proof-of-concept drug screen using our scalable PKD organ-oid model described above (**Fig. 4K**). For our candidate pool, we selected a commercially avail-able 148 small molecule library, which targets major pathways of the epigenome (**Fig. S8F and Methods**). *Pkd2*^-/-^ NPC clones #4 and #5 were used as two biological replicates for the screen. To allow robust statistical analyses, 30–50 mini organoids were seeded into each well of a 12-well plate and cultured in suspension with shaking. Three days after nephron induction, small molecules were added individually into the culture medium. Two days after drug treatment, sig-nificant differences in cyst formation were observed, so we quantified the cyst forming efficiencies and the cyst diameters (**Fig. 5H**). Out of the 148 small molecules, 14 showed significant inhibititory effects on cyst formation in at least one biological replicate (**Fig. 5H and I**). As a positive control, metfomin dose-dependently inhibited cyst formation while tolvaptan, a negative control, had no effect (**Fig. S8G**). 12 of the 14 hits identified were shared between the two biological replicates (**Fig. 5I**), among which 8 were HDAC inhibitors, and 3 were BRD4 inhibitors, two previously described treatments for ADPKD^76, 77^. The other shared hit identified was PTC-209, a specific inhibitor of BMI-1, which has not been previously reported in ADPKD. PTC-209 emerged as a top hit for cyst inhibiting properties in the screen (**Fig. 5J-L**). Secondary validation further confirmed that it can decrease cyst forming efficiency and cyst diameters in a dose-dependent manner (**Fig. S8H-J**). Taken together, a proof-of-concept drug screen was performed, leading to the discovery of a prospective drug candidate, PTC-209, for ADPKD treatment.

### YAP activation captures human NPCs with restored distal nephron differentiation potential

The broad applications of mouse NPC lines in 2D culture format inspired us to determine whether we can also derive long-term expandable human NPC lines in 2D. For that, we first performed immunostaining in human fetal kidney sections and verified that p38 MAPK activity (phosphorylated p38, p-p38, **Fig. 6B**), TGF-β signaling (phosphorylated Smad2/3, p-Smad2/3, **Fig. 6C**) and BMP signaling (phosphorylated Smad1/5/8, p-Smad1/5/8, **Fig. 6D**), are intrinsically inhibited in the primary hNPCs *in vivo*. Thus, we included p38, TGF-β, and BMP inhibitors in the culture medium to expand human NPCs as we did in mNPSR-v2. By replacing mouse LIF with human LIF in the mNPSR-v2 medium (mouse LIF does not activate human LIF receptors), we prepared a culture medium, designated as hNPSR-v1, to determine whether it was suitable for 2D expansion of human NPCs. For that, we first engineered H1 human embryonic stem cells (hESCs) with CRISPR-Cas9 to knock in a dual reporter system for *SIX2*-GFP and *PAX2*-mCherry. Following a well-established 10-day hPSC-to-NPC differentiation protocol^8^ (**Fig. S9A**), the SIX2^+^/PAX2^+^ induced NPC (iNPC) population was collected by FACS (**Fig. S9B**) and cultured in hNPSR-v1 medium. However, iNPCs were only able to stably expand for the first 2 weeks before cell proliferation slowed down with decreased NPC marker gene expression (data not shown), suggesting hNPSR-v1 medium was not optimal for long-term expansion of iNPCs.

**Figure 6.**
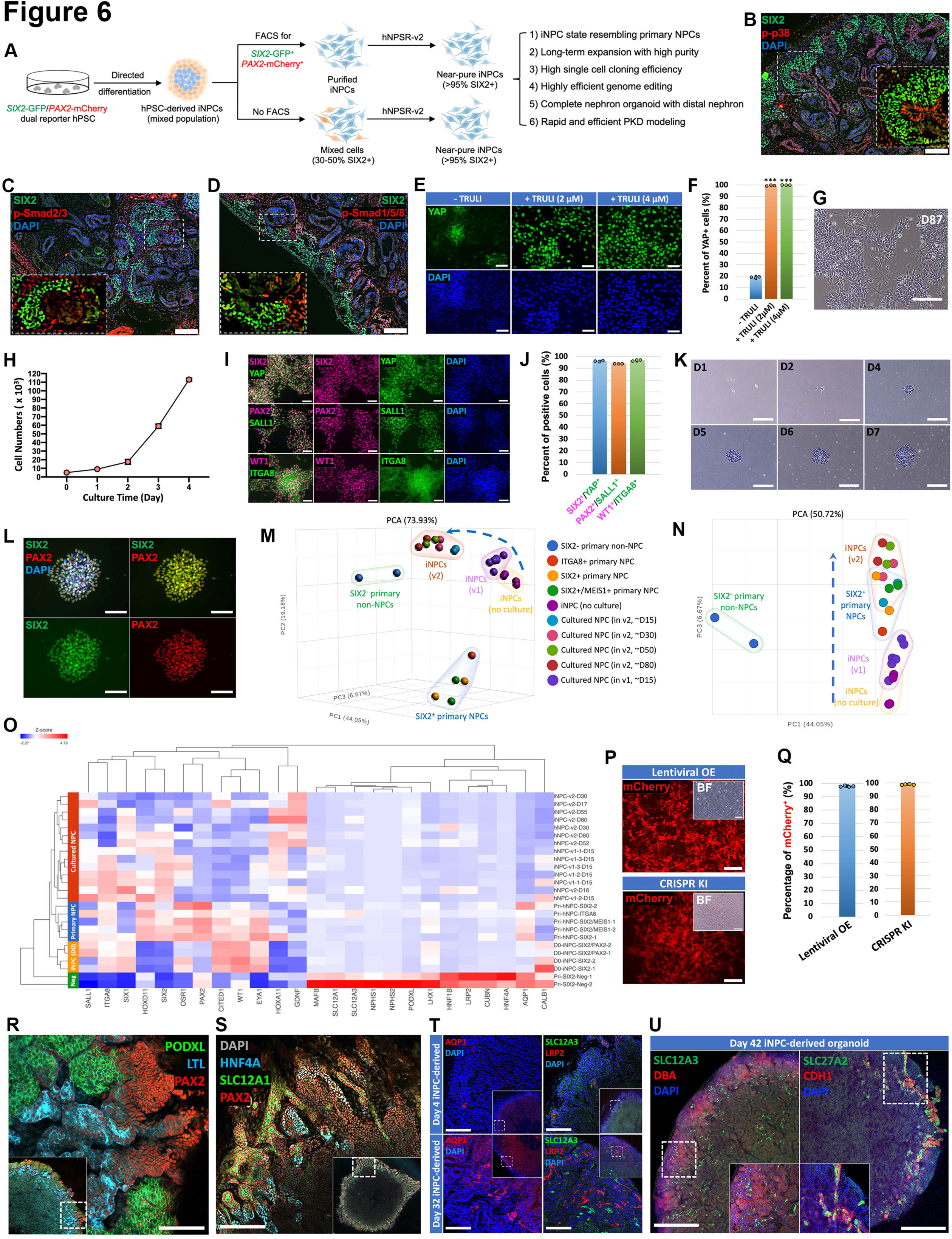
YAP activation captures human NPCs with restored distal nephron differentiation potential. (A) Schematic showing the derivation of long-term expandable iNPC lines from hPSCs and its versatile applications. (B–D) Immunofluorescence analyses of 11.3-week human fetal kidney sections for SXI2, p-p38 (B), p-SMAD2/3 (C), and p-SMAD1/5/8 (D). Scale bars, 50 μm. (E and F) Immunofluorescence analyses (E) and quantification (F) of YAP expression in iNPCs cultured in hNPSR-v1 medium supplemented with 0, 2, or 4 µM TRULI for 6 days. Scale bars, 50 μm. (G) Bright-field image of iNPCs cultured in hNPSR-v2 medium for 87 days. Scale bar, 100 μm. (H) Growth curve of iNPCs cultured in hNPSR-v2 in a typical 4-day passage cycle starting from 5,000 cells. (I and J) Immunofluorescence analyses (I) and quantification (J) of iNPCs cultured in hNPSR-v2 medium for 21 days for various NPC marker genes as indicated. Scale bars, 100 μm. (K) Time-course bright-field images showing clonal expansion of iNPCs from one single cell in hNPSR-v2 medium. Scale bars, 100 μm. (L) Immunofluorescence analysis of a single cell iNPC clone for SIX2 and PAX2. Scale bars, 50 μm. (M and N) 3D (M) and 2D (N) PCA plots of bulk RNA-seq data. (O) Heatmap showing gene expression of selected marker genes for undifferentiated NPCs and differentiated kidney cell types, in primary and cultured NPCs, as well as FACS-purified SIX2^+^ or SIX2^+^/PAX2^+^ iNPCs without further culture (D0-iNPC-SIX2 and D0-iNPC-SIX2/PAX2). Primary SIX2-negative non-NPCs (Pri-SIX2-Neg) isolated from human fetal kidneys were used as negative controls. (P and Q) Bright field (BF) and fluorescence images (P) and quantification (Q) of mCherry expression in iNPCs upon lentiviral overexpression of mCherry (lentiviral OE), or targeted CRISPR/Cas9 knock-in of mCherry-expressing cassette into *AAVS1* allele (CRISPR KI). Scale bars, 50 μm. (R and S) Whole-mount immunofluorescence analyses of human nephron organoids generated from iNPCs cultured in hNPSR-v2 medium for 42 days for various nephron marker genes as indicated. Scale bars, 200 μm. (T) Whole-mount immunofluorescence analyses of human nephron organoids generated from iNPCs cultured in hNPSR-v2 for 4 days (upper panels) or 32 days (lower panels) for various nephron marker genes as indicated. Scale bars, 200 μm. (U) Whole-mount immunofluorescence analyses of human nephron organoid generated from iNPCs cultured in hNPSR-v2 medium for 42 days for various nephron marker genes as indicated. Scale bar, 200 μm. Data are presented as mean ± SD. Each column represents counts from three biological replicates (n=3). The significance was determined by two-tailed unpaired Student’s t tests; ns, not significant; *, p<0.05; **, p<0.01; ***, p<0.001. See also Figures S9, S10, and Tables S12 and 13.

iNPCs cultured in hNPSR-v1 medium frequently lost YAP expression in the nucleus, suggesting spontaneous loss of YAP activity (**Fig. 6E**). Considering the established role of YAP activity in the self-renewal of various types of stem cells and progenitor cells^78, 79^, including NPCs^80^, we supplemented hNPSR-v1 medium with our recently reported, highly specific small molecule YAP agonist TRULI^81^. Addition of 2 µM to 4 µM TRULI to an hNPSR-v2 formulation (**Table S12**), resulted in significant nuclear accumulation of YAP in most iNPC nuclei (**Fig. 6E and F**) and long-term clonal 2D expansion of iNPCs (**Fig. 6 and S9**), and primary human NPCs (**Fig. S10**) for several months with high purity.

iNPCs cultured in hNPSR-v2 grew quickly with a typical passaging ratio of 1:10 every 3– 4 days (**Fig. 6G and H**). After 3 weeks of culture, more than 95% of cells maintained expression of SIX2/PAX2/WT1/SALL1/ITGA8 (**Fig. 6I and J**). After long-term culture of more than 3 months, all the NPC marker genes were still retained, except for a gradual decrease of *PAX2* starting after 1 month of culture (**Fig. S9C-E**). iNPCs showed a clonal efficiency of 58-70% in reseeding exper-iments (**Figs. 6K and L, S9F and G**). Importantly, the robustness of hNPSR-v2 allowed us to derive iNPC lines without prior FACS enrichment of the SIX2^+^/PAX2^+^ iNPCs (**Fig. 6A, S9H-J and Methods**), suggesting iNPC lines might be derived from any hPSC lines, which greatly broadens this application. Further, primary human NPCs were stably expanded in hNPSR-v2 medium for 100 days with the retention of NPC marker gene expression (**Fig. S10A-J**).

To compare their transcriptomes, bulk RNA-seq transcriptional profiling was performed on iNPCs FACS sorted 10 days after initiation of hPSC differentiation without further culture, and cultured human NPCs (day 15 to day 80) from different sources (iNPC or primary NPC) (**Table S13**). Their transcriptional profiles were compared to recently published bulk RNA-seq data for primary human NPCs freshly isolated from the human fetal kidney^24^. Unsupervised PCA analysis clustered the samples into 5 groups: “SIX2^-^ primary non-NPCs” (SIX2^-^ cells from human fetal kidney^24^); “SIX2^+^ primary NPCs”, “iNPCs with no culture;” “cultured NPCs in v1 medium;” and “cultured NPCs in v2 medium” (**Fig. 6M, N, and S9K**). As expected, the “SIX2^-^ primary non-NPCs” were positioned far from the other NPC groups along the PC1 axis. Interestingly, compared to iNPCs without further culture, or cultured in the v1 medium, NPCs cultured in the v2 medium were closer to the primary NPC state, which largely overlapped with the “SIX2^+^ primary NPCs” in both PC1 and PC3 axes (**Fig. 6M and N**). Consistent with PCA, heatmap of gene expression of NPC marker genes and nephron segment anchor genes also clustered the cultured NPCs closer to the primary NPCs than iNPCs without any culture (**Fig. 6O**). Expression of nephron segment anchor genes was not observed in any NPC group. However, compared to primary NPCs and cultured NPCs, iNPCs without further culture showed significantly lower expression of *SIX2*, *HOXA11*, and *HOXD11* genes, which encode transcription factors essential for NPC identity, suggesting that following 10 days of directed differentiation from hPSCs, the iNPCs might still not be sufficiently programmed to the NPC state, but after further culture, they transition to a state similar to the primary NPCs. Furthermore, we showed that genetic manipulations, including lentiviral gene over-expression and CRISPR-Cas9 based gene knock-in, can be efficiently conducted in the cultured iNPCs, opening new avenues of applications for this system (**Fig. 6P and Q**).

To further validate the nephrogenic potential of the cultured NPCs, we generated nephron organoids from the cultured NPCs, following a protocol that involves a pulse of Wnt activation followed by spontaneous self-organization (**Fig. S10K, and Methods**). Mimicking normal nephrogenesis, clear mesenchymal-to-epithelial transition was observed in the organoid in the first 3 days after Wnt activation, eventually forming numerous renal tubule-like structures with *PAX2*-mCherry reporter expression over a course of 14 days (**Fig. S9L**). Immunostaining further con-firmed the formaton of various nephron segments, including PODXL^+^ glomeruli, LTL^+^/HNF4A^+^ proximal tubule, and PAX2^+^/SLC12A1^+^ loop of Henle (**Fig. 6R and S**). Interestingly, we observed the formation of large numbers of SLC12A3^+^ distal convoluted tubule structures (**Fig. 6T and U**), which are not detected in currently available human kidney organoids^14, 15, 82–84^. Interestingly, iNPCs cultured for a short period of time of 4 days generated significantly fewer SLC12A3^+^ distal tubule structures compared to the iNPCs cultured for 32 days or 42 days (**Fig. 6T and U**). Similarly, signficiantly more AQP1^+^ thin descending limbs of loop of Henle and LRP2^+^ proximal tubules were generated from the long-term cultured iNPCs (**Fig. 6T**), and functional transporters SLC27A2 and SLC34A1 found in the proximal tubule were expressed only from the long-term cultured iNPC-derived organoids (**Fig. 6U and S9M**). Other genes that represent kidney terminal differentiation, such as *SLC22A8*, *SPTSSB*, and *UMOD*, were also expressed at higher levels in the organoids derived from iNPCs cultured long-term rather than short-term (**Fig. S9P**). Based on these observations, and the previous transcriptome analyses, we hypothesized that, upon further culture in hNPSR-v2 medium, hPSC-derived iNPCs are programmed to a state closer to the native NPC state, eventually gaining the more complete nephrogenic potential of generating all major segments of the human nephron, including the distal convoluted tubules.

Current kidney organoids generate a significant amount of off-target cell populations such as neurons and myocytes^14, 15, 82, 83^. In contrast, in the cultured iNPC-derived nephron organoids, MAP2 and NeuN, two neuron genes frequently expressed in current kidney organoids, were not detected (**Fig. S9N and O**), reflecting the high purity of the cultured NPCs, and the restricted developmental potential of NPCs only to the nephron. Similar results were observed in the nephron organoids that were derived from long-term cultured primary human NPCs (**Fig. S10L-N**).

### Rapid, efficient, and scalable PKD modeling directly from genome-edited human NPCs

Based on the robust 2D culture system for long-term expansion of human NPCs, we developed a rapid, efficient, and scalable PKD organoid model directly from genome-edited human NPCs (**Fig. 7A**). The *PKD2* gene was knocked out using CRISPR/Cas9 system in our *SIX2*-GFP reporter hPSC line. Two single cell clones, #10 and #11, with verified biallelic frame-shift mutations (**Fig. 7B-D**), were selected to derive NPC lines in hNPSR-v2 medium. Mini NPC aggregates were generated, followed by shaking culture and nephron induction (**Fig. 7A, E and Methods**). After 8 days of shaking culture, cysts emerged in the *PKD2*^-/-^ NPC-derived nephron organoids, but not in the wild-type NPC-derived con-trol organoids. These cysts continued to grow larger with the majority of the cyst-lining cells expressing LTL and CDH1 (**Fig. 7F-I**). We then verified the responses of these cystic organoids to previously described candidate PKD drug treatments. As expected, CFTRinh172, metformin, AZ505, tubacin^85^, and our newly identified small molecule PTC-209, significantly decreased cyst forming effiencies and cyst diameters, but not tolvaptan (**Fig. 7J-L**). Thus, as proof-of-concept, with this scalable mini PKD organoid model, start-ing from the expandable *PKD2*^-/-^ NPCs, a drug candidate’s effects on cyst development can be quantitiatively determined within 8 days of shaking culture, which is a remarkably faster turnaround than current PKD organoid models starting from hPSCs that require at least 3 weeks^54–58^.

**Figure 7.**
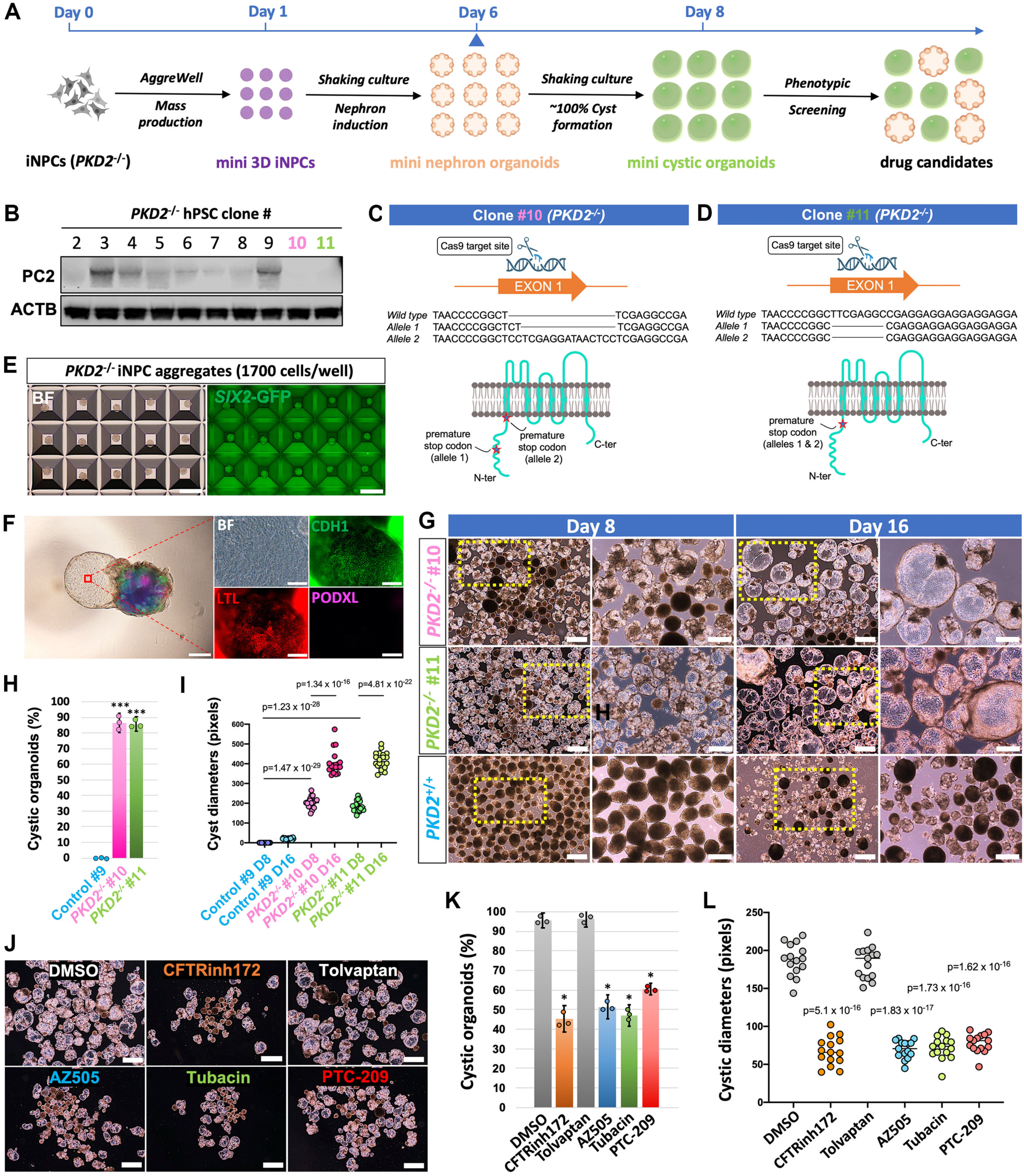
Rapid, efficient, and scalable PKD modeling from genome-edited human NPCs. (A) Schematic showing the experimental procedures for generating PKD organoids directly from expandable *PKD2^-/-^* iNPCs. (B) Western blot analysis of PC2 expression in different candidate *PKD2^-/-^* single cell hPSC clones. (C and D) Schematic showing CRISPR/Cas9-mediated random insertions or deletions (indels) on the first exon of *PKD2* gene, leading to double allele premature termination of *PKD2* transcription in *PKD2^-/-^* clonal iNPC lines #10 and #11. (E) Bright-field and GFP images of mini *PKD2^-/-^ SIX2*-GFP iNPC aggregates upon seeding onto Aggrewell plate overnight. Scale bars, 500 μm. (F) Bright-field (BF) and immunofluorescence images of a cystic nephron organoid derived from *PKD2^-/-^*iNPCs. Right panels show enlarged images of the boxed cystic area from the left panel. Scale bars, 500 μm (left panel) and 100 μm (right panels). (G) Bright-field images showing rapid cyst formation and continuous growth in the mini PKD organoids, derived from *PKD2^-/-^* iNPC lines #10 and #11. Right panels under each time point show enlarged pictures of the boxed areas from the left panels. Scale bars, 500μm and 200 μm (enlarged pictures). (H and I) Quantification of cystic organoid percentages and cyst diameters for samples shown in (G). (J) Bright-field images of *PKD2^-/-^* iNPC-derived mini cystic PKD organoids treated with various known drugs for PKD, and PTC-209. Scale bars, 500 μm. (K and L) Quantification of cystic organoid percentages and cyst diameters for samples shown in (J). Data are presented as mean ± SD. Each column represents counts from three biological replicates (n=3). The significance was determined by two-tailed unpaired Student’s t tests; ns, not significant; *, p<0.05; **, p<0.01; ***, p<0.001.

## DISCUSSION

NPCs play central roles in kidney development and disease and have great potential in kidney regenerative medicine. In this study, we redefine the critical role of p38 and YAP activity in NPC self-renewal. Based on p38 inhibition and YAP activation, we developed chemically-defined cul-ture systems for the clonal, long-term expansion of near pure mouse and human NPCs in 2D culture. The culture system constitutes a synthetic niche that recapitulates developing nephron’s plasticity. Cultured NPCs show gene expression profiles similar to primary NPCs and generate major segments of the nephron, including distal convoluted tubules, on differentiation in culture. Genome-wide CRISPR screen of cultured NPCs demonstrated the potential of the model for ge-nome-scale study of NPC biology, Wilms tumor, and CAKUT. Further, a rapid, efficient, and scal-able PKD organoid model initiated from genome-edited NPCs, highlighted the potential for *de novo* drug identification or validation. We anticipate that, with the simplicity and the robustness of the protocols, these NPC and organoid culture systems will be widely adopted by other research-ers to study different aspects of kidney development, disease, plasticity, and regeneration.

The inhibition of p38 MAPK activity proved to be key in allowing the long-term expansion of mouse and human NPCs *in vitro* in the 2D culture setting. How p38 activity and its downstream regulatory network control NPC self-renewal and differentiation remains to be determined. Simi-larly, how activation of YAP signaling programs hPSC-derived iNPCs to a state similar to the primary NPCs and then stably maintaining this state in culture also remains obscure. Our *in vitro* accessible NPC culture systems will provide an ideal platform to address these important biolog-ical questions.

As observed *in vivo*^25^, we showed that *Wnt4*^+^ nephron cells have developmental plasticity to be reprogrammed back to the NPC state *in vitro* in the NPC culture medium. Of note, *Wnt4*^+^ cells might not be the only plastic cell type in the developing kidney that can be reprogrammed to the NPC state. For example, *Foxd1*^+^ IPCs and NPCs are believed to be derived from the same precursor cells during early kidney development, with interchangeable cell identity^86, 87^. Thus, it is conceivable that IPCs might also be readily reprogrammed to the NPC state. It will also be im-portant to determine whether similar plasticity exists in the human kidney in the future. If so, the mechanisms might be employed to develop *in vivo* reprogramming strategies for kidney regener-ation and disease intervention.

Our observation that hPSC-derived iNPCs are not fully programmed to the NPC stage in existing kidney organoids agrees with a recent report where an improvement in NPC specification in organoid culture led to normalized proximal tubule specification and function though distal con-voluted tubule cells were absent from organoids in this study^16^. As the currently only available human kidney organoid system to generate abundant distal convoluted tubules, our platform will open new avenues to study how human distal nephron fate is specified from NPCs, and allow modeling of diseases that affect the distal nephron of the kidney.

PTC-209 was identified initially as a specific inhibitor of BMI-1, a gene that plays critical roles in the self-renewal of stem cells in normal tissues and cancer^88^. PTC-209 and its derivatives are being tested in preclinical and clinical trials for the treatment of serveral types of cancer^88, 89^. Considering that there are similarities between PKD and cancer pathogenesis^90^, and given the efficacy of PTC-209 in cyst inhibition in our organoid model, we propose that PTC-209 or other BMI-1 inhibitors might be prospective candidates for PKD treatment. Future investigations using animal models are warranted for BMI-1 inhibitors. In addition to PKD, we anticipate other kidney disease models can also be established from kidney organoids derived directly from genome-edited NPCs.

### Limitations of the study

Like all other cell culture systems, cultured cells *in vitro* can recapitulate key features of their counterparts *in vivo*, but may also gain characteristics that reflect *in vitro* culture. These differences usually do not prevent the broad applications of the *in vitro* cultured cells. However, when interpreting the CRISPR screen results, we anticipate a small number of genes might be identified as false positive hits due to differences based on *in vitro* culture. An example we observed is the *Pax2* gene, encoding a transcription factor well-known for maintaining NPC identity^87, 91^. It was expected to be negatively selected, considering that it is essential for NPC self-renewal, but instead, it was identified as a top positively selected gene in the CRISPR screen. Thus, although our CRISPR screen data provide a valuable list of genes that are functionally important *in vitro* in NPC self-renewal, complementary approaches are still needed to validate the functions of specific genes of interest *in vivo*, as in all other CRISPR screen studies.

## ACKNOWLEDGEMENTS

We would like to thank Jeffrey Boyd and Bernadette Masinsin of the USC Flow Cytometry Facility for FACS, Seth Ruffins of the USC Optical Imaging Facility for help with microscopy, Dejerianne Ostrow and David Ruble of the Children’s Hospital Los Angeles Molecular Pathology Genomics Core for RNA-seq, Yibu Chen of the USC Norris Medical Library Bioinformatics Service for help with the RNA-seq computational analysis, Dr. Melissa L. Wilson (Department of Preventive Medicine, University of Southern California) and Family Planning Associates for coordinating fetal tissue collection, and Cristy Lytal for help with editing the manuscript. This work was supported by UKRO foundation funding, a KSOM Dean’s Pilot Award and an NIH Director’s New Innovator Award (DP2DK135739) to Z.L., and NIH DK054364 to A.P.M. Z.Z. was supported by a USC Stem Cell Challenge Award. M.E.S. was supported by a CIRM Bridges Award.

## AUTHOR CONTRIBUTIONS

Conceptualization, B.H. and Z.L.; methodology, investigation and validation, B.H., Z.Z., H.L., X.C., J.G., C.C.Z., M.E.S., A.C.V., T.X., T.P., Y.L., R.K.P., B.D., and J.H.C.; software and formal analysis, B.H., Z.Z., Z.L., and T.F.; resources, M.E.T., B.H.G., Y.D., Y.D., Q.Y., K.G., N.O.L., and A.P.M.; writing – original draft, B.H. and Z.Z.; writing – review and editing, T.F., N.M.P.S., K.R.H., A.P.M. and Z.L.

## STAR METHODS

## KEY RESOURCES TABLE

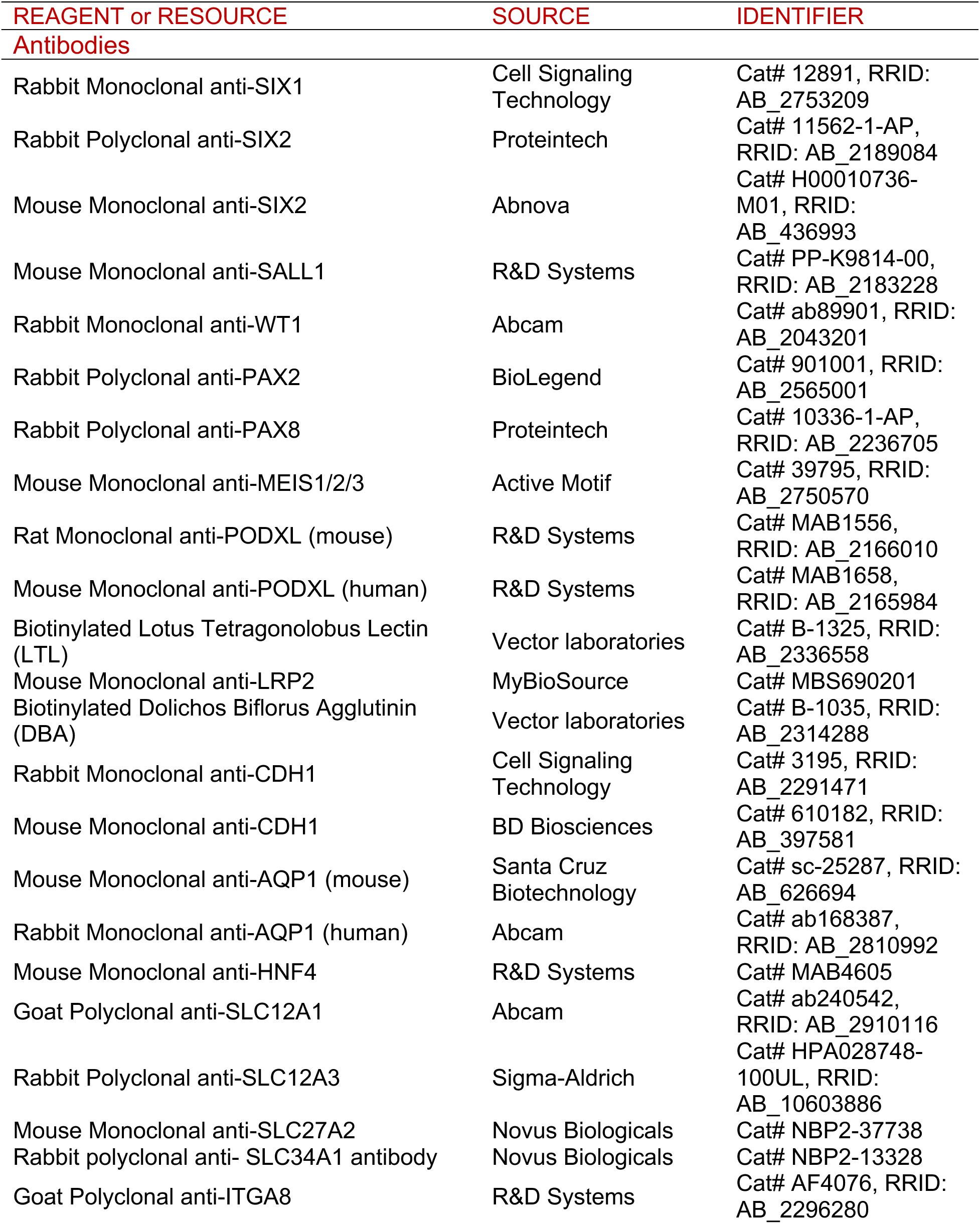
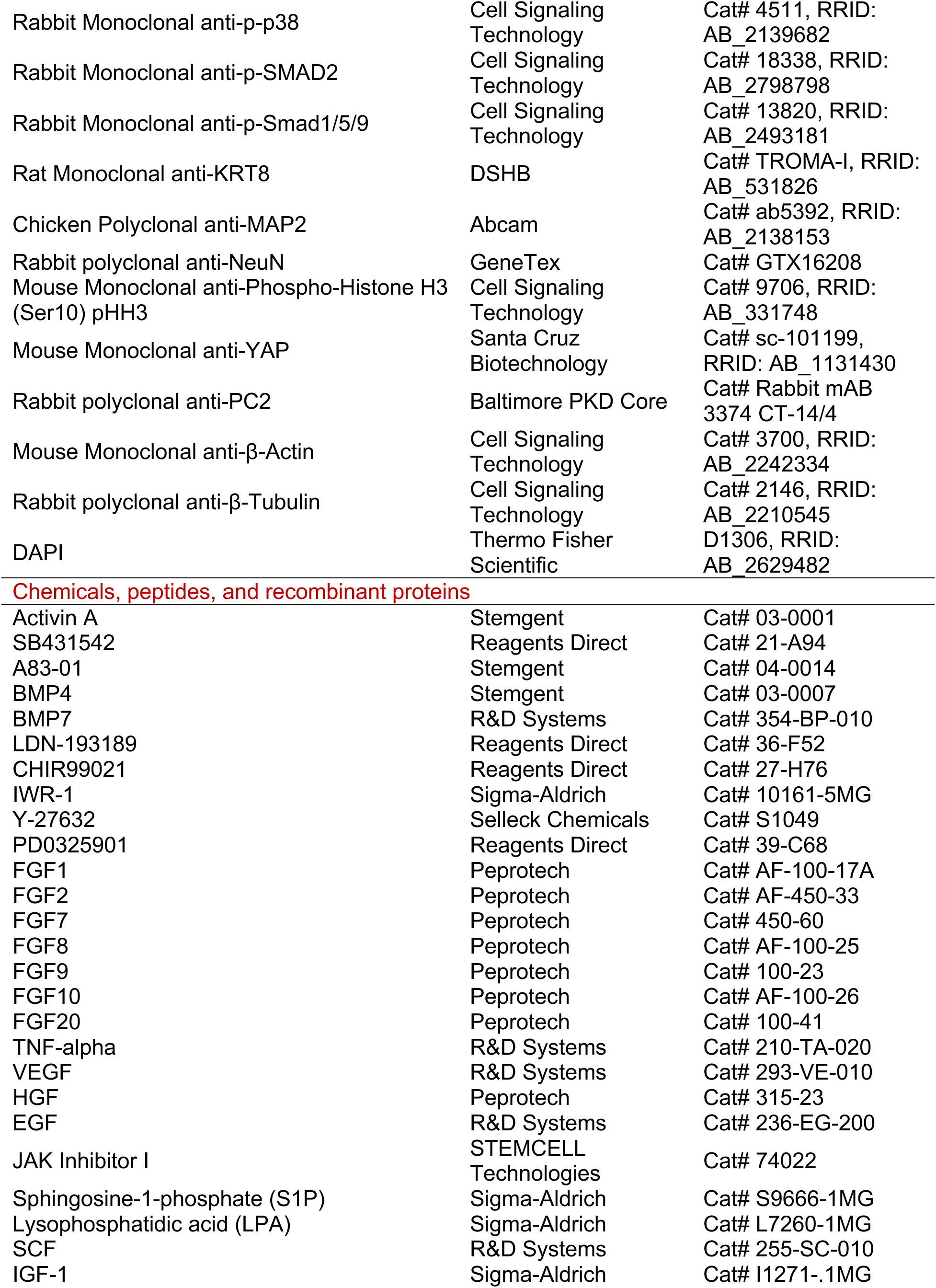
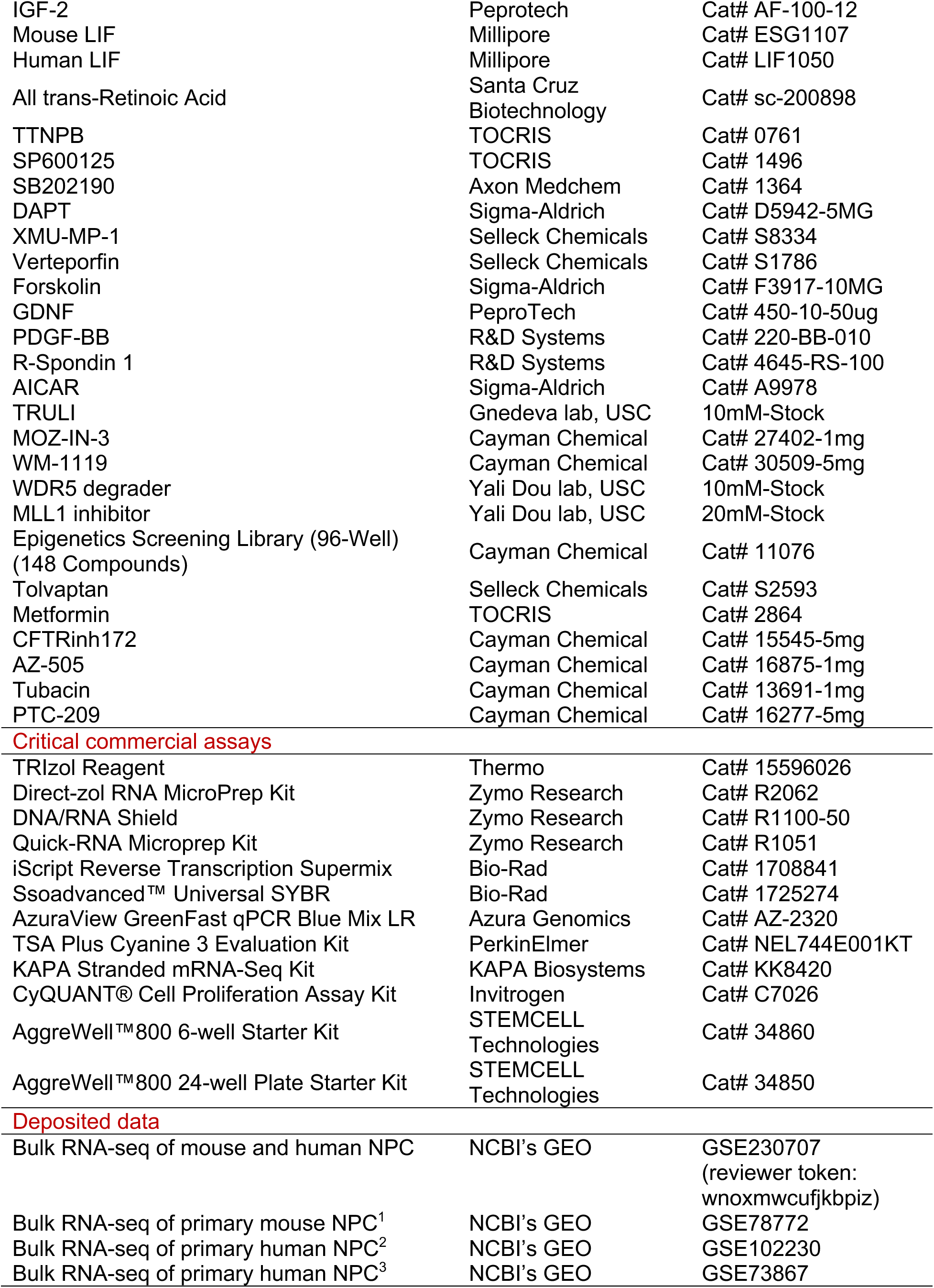
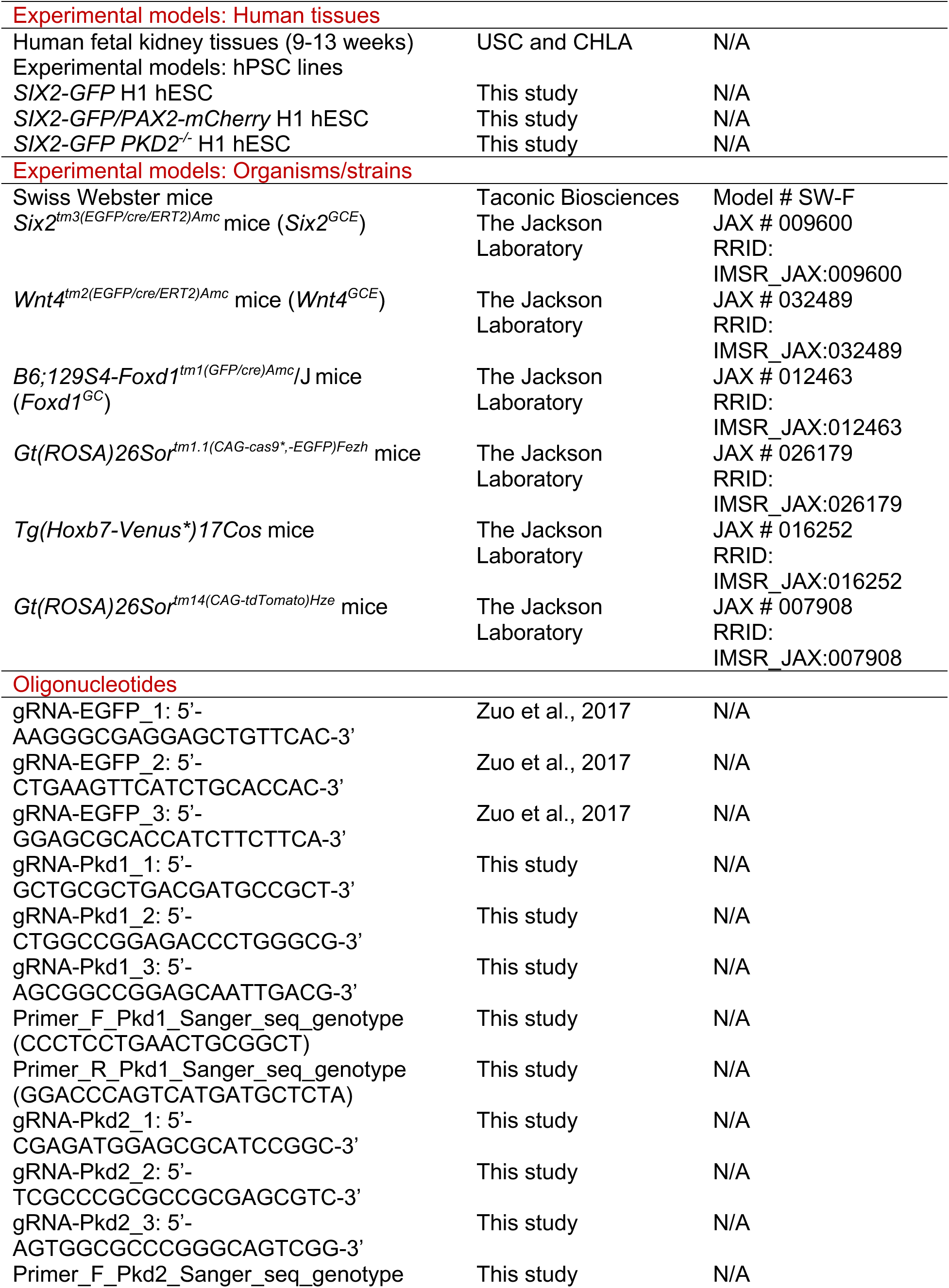
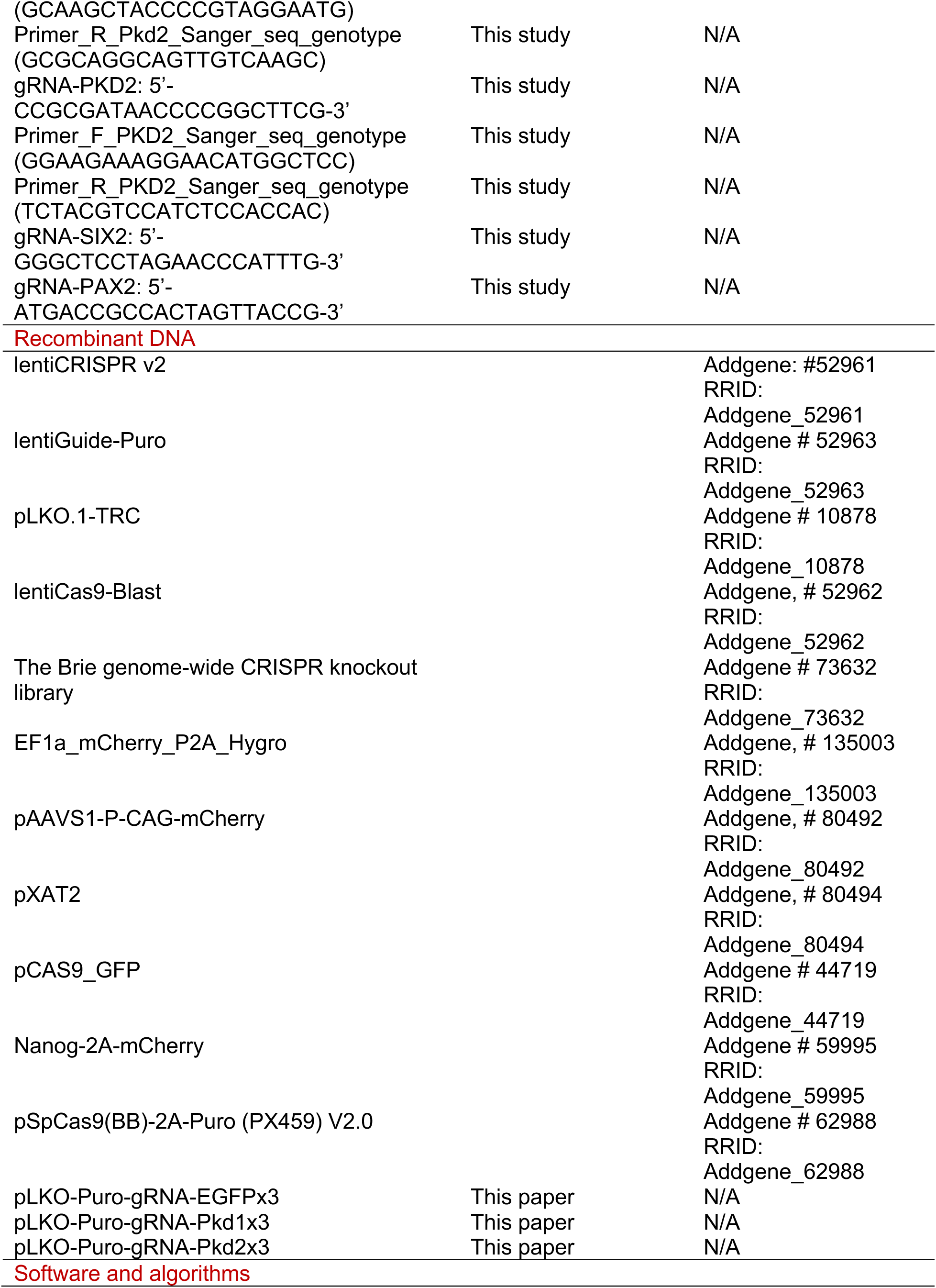
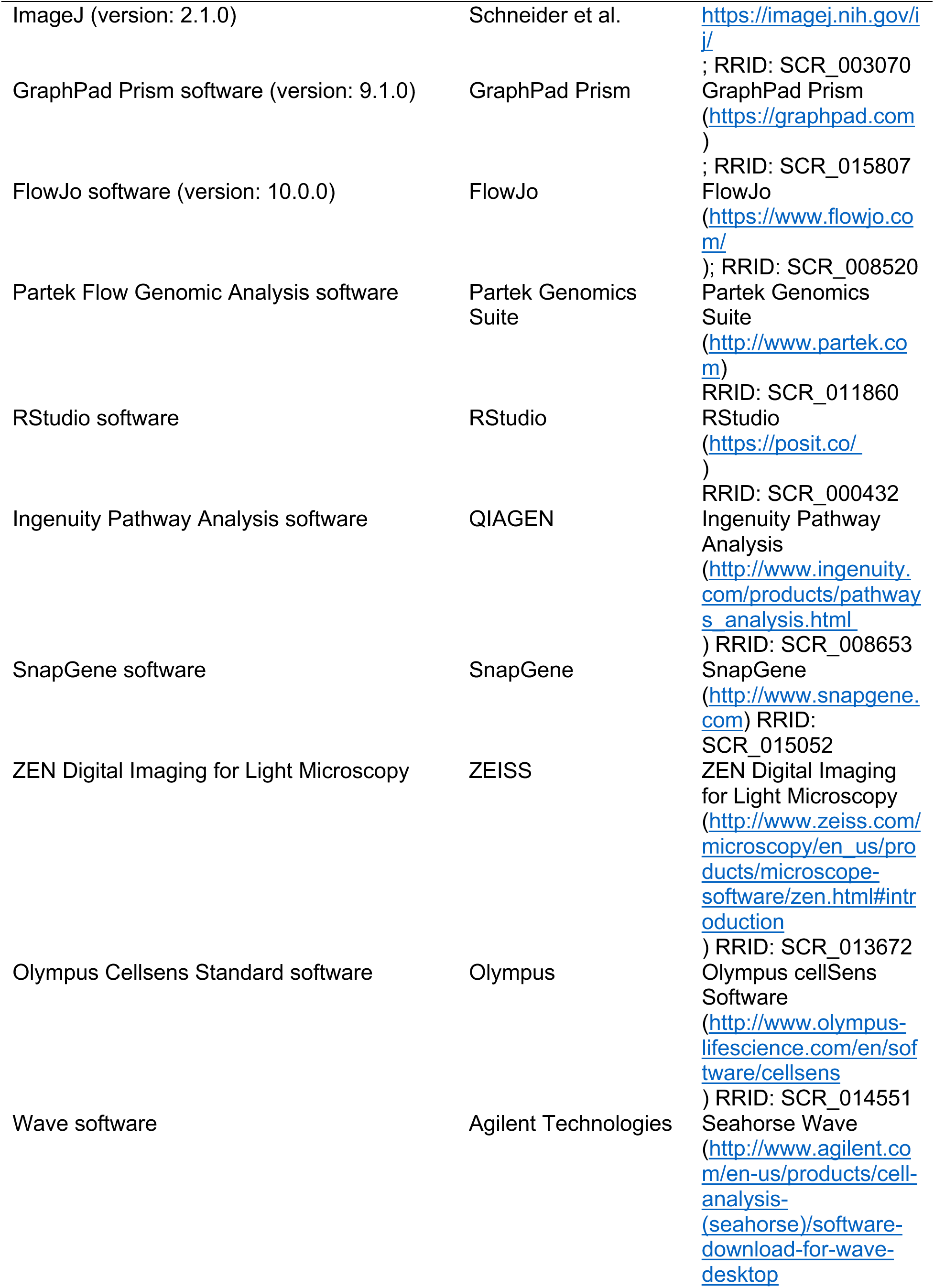
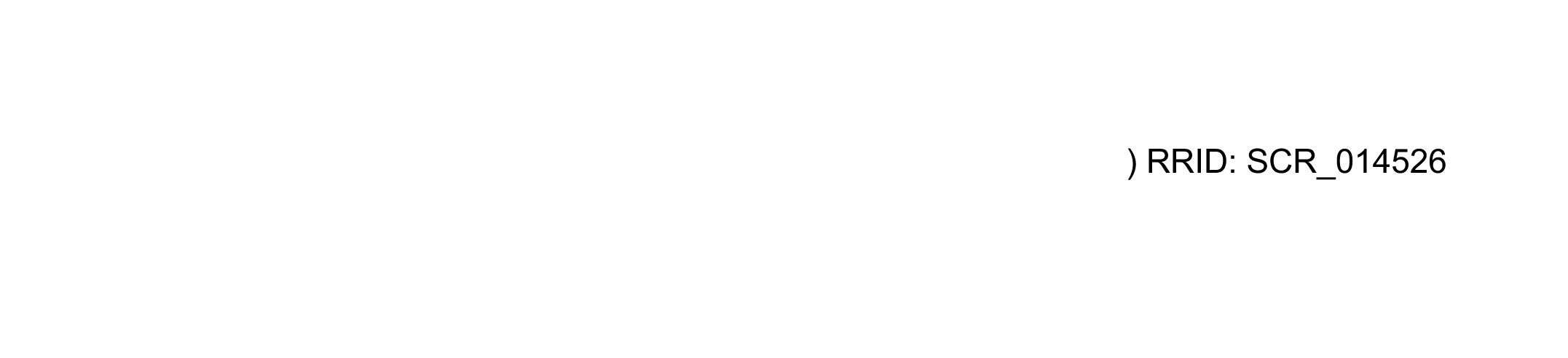

## RESOUCE AVAILABILITY

### Lead contact

Further information and requests for resources and reagents should be directed to and will be fulfilled by the lead contact, Zhongwei Li.

### Materials availability

Cell lines, plasmids, and other unique resources generated in this study are available from the lead contact, Zhongwei Li, with a completed Materials Transfer Agreement.

### Data and code availability

RNA-seq data have been deposited at GEO (GSE230707) with reviewer security token: wnoxmwcufjkbpiz. The data will be made publicly available upon publication. Any additional information required to reanalyze the data reported in this paper is available from the lead contact upon request.

## EXPERIMENTAL MODEL AND SUBJECT DETAILS

### Human tissues

All human fetal kidney samples were collected under Institutional Review Board approval at both the University of Southern California and Children’s Hospital Los Angeles (USC-HS-13-0399 and

CHLA-14-2211). Consent for tissue donation was obtained after the patient had already made the decision for pregnancy termination by Dilation and Curettage or Dilation and Evacuation and was obtained by a different clinical staff member than the physician performing the procedure. All tissues were de-identified, and the only clinical information collected was gestational age and the presence of any maternal or fetal diagnoses.

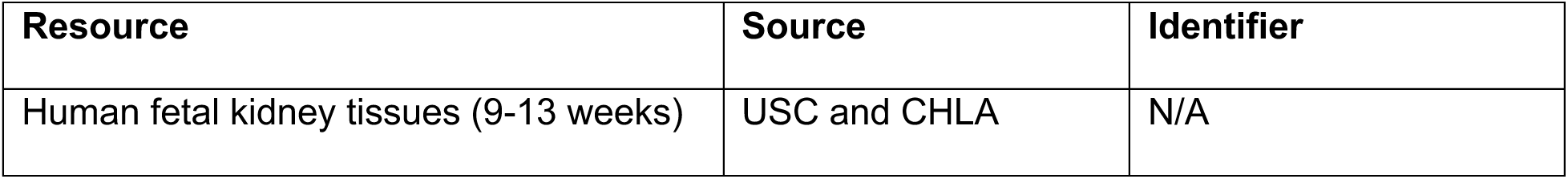

### Mice

All animal work was performed under Institutional Animal Care and Use Committee approval (USC IACUC Protocol # 20829). Swiss Webster mice were purchased from Taconic Biosciences (Model # SW-F). *Six2^tm3(EGFP/cre/ERT2)Amc^* mice (*Six2^GCE^*, JAX # 009600), *Wnt4^tm2(EGFP/cre/ERT2)Amc^* mice (*Wnt4^GCE^*, JAX # 032489), *Foxd1^tm1(GFP/cre)Amc^* mice (*Foxd1^GC^*, JAX# 012463), *Gt(ROSA)26Sor^tm1.1(CAG-cas9*,-EGFP)Fezh^*mice (JAX # 026179), and *Tg(Hoxb7-Venus*)17Cos* mice (JAX # 016252) were kindly shared from Dr. Andrew McMahon. *Gt(ROSA)26Sor^tm14(CAG-tdTomato)Hze^* mice (tdTomato reporter mice, JAX # 007908) were kindly shared from Dr. Kenneth Hallows. *Six2-tdTomato (Six2-tdT)* mice were generated through crossing *Six2^GCE^* mice and tdTomato reporter mice. *Wnt4-tdTomato (Wnt4-tdT)* mice were generated through crossing *Wnt4^GCE^* mice and tdTomato reporter mice.

### hPSC lines

Experiments using hPSCs were approved by the Stem Cell Oversight Committee (SCRO) of University of Southern California under protocol # 2018-2. Human pluripotent stem cells are routinely cultured in mTeSR1 (STEMCELL Technologies #85850) or mTeSR1 Plus (STEMCELL Technologies #100-0276) medium in monolayer culture format coated with Matrigel and passaged using dispase as previously described ^1^, or using Versene Solution (Thermo Fisher # 15040066) following manufacturer’s protocols.

| Resource | Source | Identifier |
| --- | --- | --- |
| *SIX2-GFP* H1 hESC | This study | N/A |
| *SIX2-GFP/PAX2-mCherry* H1 hESC | This study | N/A |
| *SIX2-GFP PKD2<sup>-/-</sup>* H1 hESC | This study | N/A |

## METHOD DETAILS

### Deriving clonal expandable NPC lines from any mouse strain with mNPSR-v2 medium

#### Deriving NPC lines from *Six2-*GFP mice

Timed pregnant E12.5-E18.5 kidneys were isolated from *Six2-*GFP (*Six2^GCE^*, JAX # 009600) mouse embryos, and then the kidneys were minced into small pieces. The minced kidney pieces were transferred into 1.5mL Eppendorf tubes and spun down at 300 g for 3 minutes. (Note that we aliquoted the kidney pieces into multiple tubes to ensure the volume of the tissue pellet after centrifugation was less than 100 µl per tube to ensure best dissociation). The dissection medium was then carefully aspirated. The kidney pieces were then washed once with sterile PBS. 500 µL of pre-warmed Accumax Cell Dissociation Solution (Innovative Cell Technologies, Cat. No. AM-105) was added to the tube to resuspend the kidney pieces. The tube was incubated in 37°C incubator for 20∼22 minutes. 500 µL 10% FBS medium (10% FBS in DMEM) was added to the tube, and then GENTLY pipetted up and down 20 to 25 times to further dissociate the kidney pieces. The tube was spun down and the supernatant was then removed. FACS medium (cold PBS with 2% FBS) was added to resuspend the cell pellet and the cell suspension was filtered through 40 µm cell strainer (Greiner bio-one, Cat. No. 542040) to remove cell clumps before FACS to sort out *Six2-*GFP^+^ NPCs. Purified *Six2-*GFP^+^ NPCs were counted with TC20™ Automated Cell Counter (Bio-Rad, Cat. No. 1450102), and then cultured in mNPSR-v2 medium (Table S2) to establish NPC lines. Take 96-well plate for example, we counted and seeded 5,000 FACS-purified NPCs into one Matrigel-coated well with 100 µl mNPSR-v2 medium. The medium was refreshed 2 days after cell seeding. When NPCs grew to 80-90% confluent (on day 3 or day 4), the culture medium was removed, and the cells were washed once with 100 µl PBS, and then dissociated with 50 µl pre-warmed Accumax. The cells were incubated with Accumax for 8 mins, and then 150 µl 10% FBS medium was added to neutralize Accumax. The medium was pipetted up and down GENTLY for 5-7 times to make single cell suspension, which was then seeded at 1:20-1:30 passage ratio to a new well in 96-well plate. Change medium 2 days after seeding. On day 3 or day 4, the cells grew to 80-90% confluent and can be passaged again using the same protocol described above. (Note, for coating with Matrigel (R&D Systems, # 3433-010-01) in one well of 96-well plate, dissolve 1 mg Matrigel into 25 ml cold DMEM/F12, and then aliquot 100 µl medium into each well. The plate is then incubated in 37°C for at least 1-2 hours before the coating medium is aspirated followed by cell seeding.)

#### Deriving NPC lines from E11.5 metanephric mesenchyme (MM)

E11.5 kidneys were isolated and MM was manually dissected out from the E11.5 kidneys following our previously described protocol to isolate E11.5 UB and MM (Zeng et al., 2021). 20 isolated MM were pooled together and dissociated in 500 µl Accumax for 8-10 minutes (scale it down if less MM were isolated) before 500 µl 10% FBS medium was added to neutralize Accumax. The medium was then pipetted up and down GENTLY for 7-10 times to dissociate the MM into single cells. 5,000 cells were then seeded into one well in a Matrigel-coated 96-well plate (scale up or down based on the surface area, if different culture format was used) in mNPSR-v2 medium to derive NPC line, using similar protocols described above starting from *Six2*-GFP^+^ NPCs.

#### Deriving NPC lines from whole kidneys

E12.5-E15.5 kidneys were isolated, minced into small pieces, and dissociated in Accumax as described above for dissociating E12.5-E18.5 *Six2*-GFP kidneys. 5,000 cells were then seeded into one well in a Matrigel-coated 96-well plate (scale up or down if different culture format was used) in mNPSR-v2 medium to derive NPC line, using similar protocols described above starting from *Six2*-GFP^+^ NPCs. Note that, different from NPC line derivation from *Six2*-GFP^+^ NPCs, or from isolated E11.5 MM, whole kidney cells need to go through 2-3 passages to enrich the NPC population in the culture before a stable NPC line can be established with 90-95% purity.

#### Deriving single cell clonal mouse NPC lines

NPC lines can be derived from FACS-purified *Six2*-GFP+ NPCs, isolated E11.5 MM, or whole kidney cells as described above. When NPC lines are stably established, single cell clonal NPC lines can be generated. For that, when the NPC lines grew to around 80% confluency, the cells were dissociated into single cells following the protocol described above for passaging NPCs. The cells were counted and seeded into Matrigel-coated 96-wells at the density 0.5 cell per well with 100 µl mNPSR-v2 medium so that most wells would have either one single cell or no cell (day 0). On day 3, 50 µl used medium was removed, and 100 µl fresh medium was added. On day 6, the used medium was completely removed and 100 µl fresh medium was added. On day 9, NPC clones were clearly observed under the microscope in about 30% of the wells seeded. Cells in these wells were dissociated following our protocol described above, and all the cells were seeded into another 96-well. On day 11, the cells reached around 80-90% confluency and were passaged routinely thereafter every 3 days as described above. To determine cloning efficiency, 60 wells of a 96-well plate were seeded with single cell NPC following the method mentioned above (D0). On day 3, the wells containing clusters of cells from a single cell were labeled and counted, and the wells without any cell were discontinued. On day 9, the number of wells grown at least 50% confluency was counted. Cloning efficiency was calculated by using the number of wells recorded on day 9 divided by the number of wells recorded on day3, from three independent experiments.

### Generation of mouse nephron organoids from mouse NPC lines

mNPCs cultured in mNPSR-v2 medium were dissociated into single cells using Accumax as described above. 30,000 cells were seeded into one well of a U-bottom 96-well plate (Thermo Fisher Scientific, # 174929) and cultured with 100 μl mNPSR-v2 medium overnight for cells to aggregate. Nephron organoids were formed thereafter following the protocol we described previously^1^. Briefly, on the next day (Day 0), 3D NPC aggregates were transferred onto 6-well format transwell membrane (Corning, # 3450) with 1.2 mL KR5-CF medium at the bottom chamber and cultured for 2 days. On Day 2, change medium to 1.2 mL KR5 medium and change medium every other day. Samples were harvested on Day 7 for various assays.

#### KR5 medium

Basal medium: DMEM/F12 (1:1) (1X), Invitrogen, Cat. No. 11330-032.

Supplements:

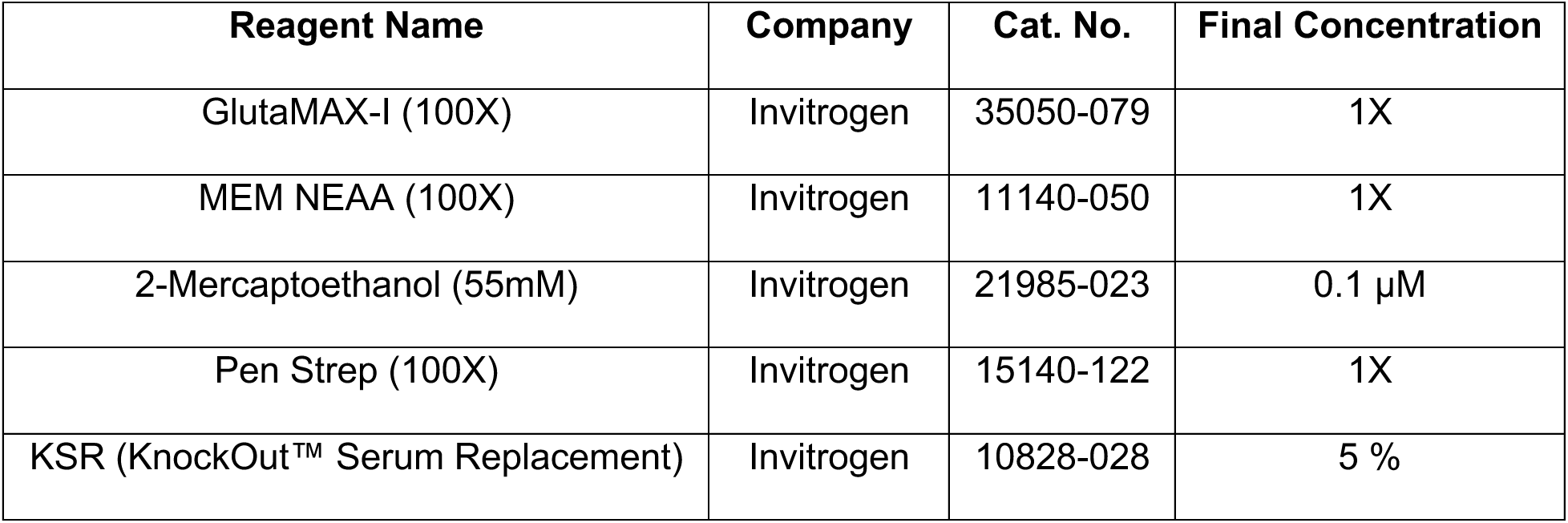

#### KR5-CF medium

KR5 medium with 4.5 μM CHIR99021 (C) and 200 ng/mL FGF2 (F).

### Spinal cord induction assay

After mNPSR-v2 cultured mNPCs reached 80-90% confluency in 2D culture, cells were dissociated into single cells using Accumax Cell Dissociation Solution. 30,000 cells were seeded into each well of U-bottom 96-well plate with 100 μl mNPSR-v2 medium and cultured overnight for re-aggregation. On the next day (Day 0), spinal cord was isolated from E12.5 embryo, and 3D mNPC aggregates were transferred to 6-well format transwell membrane tightly close to spinal cord dorsal part with 1.2mL KR5 in bottom well. Medium was changed every other day with fresh KR5 medium, and the samples were harvested for various assays on day 7.

### Mouse engineered kidney generation from cultured NPCs and UB

Mouse UB (mUB) was cultured as we previously described^4^. The day before mouse kidney reconstruction, mNPCs were dissociated and 50,000 cells were seeded into one well of U-bottom 96-well low-attachment plate with 100µL mNPSR-v2 medium and cultured overnight in 37°C incubator to generate 3D mNPC aggregate. A small piece (with 6-10 branching tips) of day 5-10 cultured mUB organoid was manually dissected out using sterile needles and inserted into a microdissected hole in a 3D mNPC aggregate (sterile needle was used to pierce a hole in the center of mNPC aggregate sphere). This structure was then transferred into a well of a U-bottom 96-well low-attachment plate with 100 µl kidney reconstruction medium (APEL2 + 0.1 µM TTNPB) plus 10 µM Y27632, using a P200 pipette with the top 0.5-1 cm of the tip cut to widen the tip, and cultured in 37°C incubator (day 0). After 24 h (day 1), the reconstructed UB/NPC structure was then transferred onto a 12-well transwell insert membrane (Corning, Cat. No. 3460). 300 µl kidney reconstruction medium was added to the lower chamber of the transwell. Medium was changed every two days for a total of 7 days while the reconstructed kidney branching and maturation progressed.

### Chicken chorioallantoic membrane (CAM) assay

CAM assay was conducted as a cost-effective method to evaluate the nephrogenic potential of cultured NPCs *in vivo*. Briefly, *Cas9*-GFP mNPC lines were dissociated into single cells, and then 30,000 cells were seeded into 96-well U-bottom low-attachment plate (Thermo Fisher Scientific, # 174929) in mNPSR-v2 medium to generate 3D NPC aggregate. On the next day (D0), change medium with 100ul KR5-CF medium and follow by continuous two days culture (D0-D2). On day 2, transfer 3D NPC aggregates into chicken chorioallantoic membrane. On day 5 and day 7, use microscope to take videos to record the vasculature system in implants. On day 7, harvest the samples for various assays.

### Dissociation of postnatal P3-P7 mouse kidneys and further culture in mNPSR-v2

P3-P7 pups were euthanized, and the kidneys were dissected. Mince one kidney into small pieces using blade (Cincinnati Surgical, # 75870-580). Collect small amount of kidney pieces into an Eppendorf tube (after spinning down, the pellet volume should be less than 30 µl). Spin down at 300g for 5min, remove the supernatant and resuspend the pellet with 500 µl warmed-up FRESH 1X Collagenase IV (Thermo Fisher Scientific, # 17104019). Put the Eppendorf tube into rotating shaker (Eppendorf ThermoMixer® F2.0) set at 37C with 800rpm shaking speed. Every 10min, take out the tube and pipet up and down the samples for 10 times. Dissociation in Col IV for a total of 30min. After 30min and the 3^rd^ pipetting, spin the cells down at 300g for 1min to collect the kidney tubules and glomerulus as the pellet. Resuspend the pellet with warmed-up 500ul Accumax. Put the Eppendorf tube into rotating shaker set at 37C with 800rpm shaking speed. Every 10min, take out the tube and pipet up and down the samples for 10 times. Dissociation in Accumax a total of 30min. After 30min, add 500 µl 10% FBS to neutralize the dissociation. Then start the 3rd pipetting for 20 times. Spin down the cells at 300g for 5min, remove supernatant, and resuspend the pellet with 200 µl ACK lysing buffer (Thermo Fisher Scientific, # A1049201), to remove the blood cells (room temperature for 3min). Add 1ml 10% FBS and spin the cells down at 300g for 5min. Remove the 1.2ml supernatant and resuspend the cells into 100 µl 10% FBS medium for cell counting (use Trypan Blue to determine the live cell percentage). Usually, the live cell percentage of dissociated cells was 50-75%. Dissociated whole kidney cells were further cultured with mNPSR-v2 medium to induce *Six2*+ NPC-like cells. Note that for this culture, iMatrix-511 (Nacalai USA, # 892021), but not Matrigel, was used for coating plates, to enhance the attachment of the postnatal kidney cells to the plate.

### Generation of *SIX2-*GFP*/PAX2-*mCherry knock-in dual reporter hPSC line

CRISPR-Cas9 based genome editing was used to insert 2A-EGFP-FRT-PGK-Neo-FRT or 2A-mCherry-loxP-PGK-Neo-loxP cassette downstream of the stop codon (removed) of endogenous *SIX2* or *PAX2* gene, respectively. DNA sequences ∼1 kb upstream and ∼1 kb downstream of the stop codon for endogenous *SIX2* (upstream F: CCGGAATTCTGCCCAGTTTGGAGCTACAG; upstream R: TACGAGCTCGGAGCCCAGGTCCACGAGGTT; downstream F: CGCGTCGACAACCCATTTGCCTTGATGAG; downstream R: CCCAAGCTTCCCGAAGAACATTCACATGAGG) or *PAX2* (upstream F: GAAGTCGACTTTCCACCCATTAGGGGCCA; up-stream R: TATGCTAGCGTGGCGGTCATAGGCAGCGG; downstream F: TATAC-GCGTTTACCGCGGGGACCACATCA; downstream R: GACGGTACCAGTAACTGCTGGAG-GAAGAC) were cloned as homology arms upstream and downstream of 2A-EGFP-FRT-PGK-Neo-FRT (for *SIX2*-GFP) or 2A-mCherry-loxP-PGK-Neo-loxP cassette (for PAX2-mCherry), respectively, to facilitate homologous recombination. 2A-EGFP fragment was cloned from pCAS9_GFP (Addgene # 44719) and the FRT-PGK-Neo-FRT cassette was cloned from pZero-FRT-Neo3R (kindly provided by Dr. Keiichiro Suzuki). 2A-mCherry-loxP-Neo-loxP fragment was cloned from Nanog-2A-mCherry plasmid (Addgene # 59995). The different fragments were then cloned to a modified pUC19 plasmid with additional restriction sites inserted, to make the complete donor plasmids for both knock-in experiments. Oligos for making sgRNA-expressing plasmid for *SIX2* knockin (F: CACCGGGGCTCCTAGAACCCATTTG; R: AAAC CAAATGGGTTCTAGGAGCCCC) or *PAX2* knockin (F: CACCGATGACCGCCACTAGTTACCG; R: AAACCGGTAACTAGTGGCGGTCATC) were synthesized, annealed, and cloned into the pSpCas9(BB)-2A-Puro (PX459) V2.0 plasmid (Addgene # 62988). Both donor and sgRNA plasmids for *SIX2* reporter knockin were transfected into the H1 hESCs using the Lipofectamine 3000 Transfection Reagent (Invitrogen, Cat. No. L3000015). Neomycin-resistant single cell colonies were picked up manually and genotyping was performed based on PCR. Clones with biallelic knock-in of *SIX2*-GFP were chosen for second round screen where plasmid encoding flippase was delivered via transfection to allow the transient expression of flippase, whose activities excise the FRT-flanked PGK-Neo cassette from the *SIX2*-GFP knock-in alleles. PCR was performed to identify single cell clones in which PGK-Neo cassettes were excised from both alleles. Then the same strategy was used to knock in *PAX2* reporter based on the successful biallelic *SIX2*-GFP knock-in clones.

### *PKD2* knock-out in *SIX2*-GFP knock-in reporter hPSC line

*SIX2-*GFP knock-in reporter hPSC line was generated using the method described above. We further knocked out *PKD2* gene in the reporter hPSC line using CRISPR/Cas9 based genome editing. For that, sgRNA was designed to target the first exon of human *PKD2* gene with sgRNA targeting sequence: CCGCGATAACCCCGGCTTCG. sgRNA was inserted into lentiCRISPR v2 plasmid (Addgene # 52961) and lentivirus was produced and then infected into *SIX2*-GFP hPSC line. After puromycin selection followed by clonal expansion of hPSCs, 11 single cell clones were picked up and expanded. Proteins were extracted from the 11 single cell clones and Western blot was performed to identify the candidate *PKD2*^-/-^ clones #10 and #11. PCR-based genotyping from the genomic DNA, following by Sanger sequencing, further confirmed the generation of frame-shift mutations in both alleles of *PKD2* gene from #10 and #11 clones.

### Deriving iNPC lines from human pluripotent stem cells

#### Deriving iNPC lines from FACS-purified *SIX2*-EGFP/*PAX2*-mCherry iNPCs

Directed differentiation from *SIX2*-GFP/*PAX2*-mCherry knock-in dual reporter hPSCs into iNPCs was performed following a previously published protocol^5^, with minor modifications. Briefly, hPSCs were dissociated into single cells with Accumax and 40,000 cells were seeded into one well in a 12-well plate with 1ml mTeSR medium plus 10 µM Y27632. Medium was changed daily with fresh 1ml mTeSR medium without Y27632 for another 2 days. At this time (day 0), hPSCs formed small colonies and were ready for directed differentiation. Phase 1, day 0 to day 4 (D0-D4), hPSCs were cultured with 1 ml Advanced RPMI 1640 Medium (Thermo Fisher Scientific, # 12633-012) supplemented with 8 µM CHIR99021 and 10nM LDN193189; medium was refreshed on D2 and D3. Phase 2 (D4-D7), medium was changed to Advanced RPMI 1640 Medium supplemented with 10 µM Y27632 and 10 ng/ml activin A; medium was refreshed daily. Phase 3 (D7-D10), medium was changed to Advanced RPMI 1640 Medium supplemented with 50 ng/ml FGF9; medium was changed daily till D10. On D10, cells were dissociated into single cells with pre-warmed Accumax, and *SIX2*-GFP/*PAX2*-mCherry iNPCs were sorted out through BD FACSAria™ III Cell Sorter. Sorted iNPCs were counted and 10,000 cells were seeded into one well in a 96-well plated coated with Matrigel and cultured with 100 µl hNPSR-v2 medium (Table S12). The medium was refreshed 2 days after cell seeding. When iNPCs grew to 80-90% confluent (on day 3 or day 4), the culture medium was removed, and the cells were washed once with 100 µl PBS, and then dissociated with 50 µl pre-warmed Accumax. The cells were incubated with Accumax for 8 mins, and then 150 µl 10% FBS medium was added to neutralize Accumax. The medium was pipetted up and down GENTLY for 5-7 times to make single cell suspension, which was then seeded at 1:10 passage ratio to a new well in 96-well plate. Change medium 2 days after seeding. On day 3 or day 4, the cells grew to 80-90% confluent and can be passaged again using the same protocol described above. (Note, Matrigel coating protocol for iNPC culture is the same as the one described for mNPC culture.)

#### Deriving iNPC lines without prior FACS-based purification of iNPCs

hPSCs were differentiated following the same protocol described above to generate iNPCs. On D10 of differentiation, instead of using FACS to purify the iNPCs based on *SIX2*-GFP; *PAX2*-mCherry dual reporter system, the dissociated whole cells were counted and then 10,000 cells were seeded into one well of 96-well coated with Matrigel and cultured with 100 µl hNPSR-v2 medium. Thereafter, after 2 weeks of continuous culture and passage in hNPSR-v2 medium following protocol described above, stable iNPC line was established with 90-95% purity of iNPCs.

#### Deriving single cell clonal iNPC lines

When iNPC lines are stably established, single cell clonal iNPC lines can be generated. For that, when the iNPC lines grew to around 80% confluency, the cells were dissociated into single cells following the protocol described above for passaging iNPCs. The cells were counted and seeded into Matrigel-coated 96-wells at the density 0.5 cell per well with 100 µl hNPSR-v2 medium so that most wells would have either one single cell or no cell (day 0). On day 3, 50 µl used medium was removed, and 100 µl fresh medium was added. On day 6, the used medium was completely removed and 100 µl fresh medium was added. On day 9, NPC clones were clearly observed under the microscope in about 30% of the wells seeded. Cells in these wells were dissociated following our protocol described above, and all the cells were seeded into another 96-well. On day 12-14, the cells reached around 80-90% confluency and were passaged routinely every 3 or 4 days at the ratio of 1:10 as described above for bulk iNPC culture. To determine cloning efficiency, 60 wells of a 96-well plate were seeded with single cell NPC following the method mentioned above (D0). On day 3, the wells containing clusters of cells from a single cell were labeled and counted, and the wells without any cell were discontinued. On day 9, the number of wells grown at least 40% confluency was counted. Cloning efficiency was calculated by using the number of wells recorded on day 9 divided by the number of wells recorded on day3, from three independent experiments.

#### Deriving hNPC lines from human fetal kidneys

Tweezers were utilized to dissect nephrogenic zones from 9 to 12 week-old human fetal kidneys. Nephrogenic zones were minced into small pieces, and transferred into 1.5 mL Eppendorf tubes, and spun down at 300 g for 3 minutes. (Note that we aliquoted the kidney pieces into multiple tubes to ensure the volume of the tissue pellet after centrifugation was less than 100 µl per tube to ensure best dissociation). The dissection medium was then carefully aspirated. The kidney pieces were then washed once with sterile PBS. 500 µL of pre-warmed Accumax was added to the tube to resuspend the kidney pieces. The tube was incubated in 37°C incubator for 20∼22 minutes. 500 µL 10% FBS medium (10% FBS in DMEM) was added to the tube, and then GENTLY pipetted up and down 20 to 25 times to further dissociate the kidney pieces. The tube was spun down and the supernatant was then removed. FACS medium (cold PBS with 2% FBS) was added to resuspend the cell pellet and the cell suspension was filtered through 40 µm cell strainer (Greiner bio-one, Cat. No. 542040) to remove cell clumps. Cells were counted and diluted to 1 million cells per 100 µl in FACS medium, and ITGA8 antibody (R&D Systems, # AF4076) was added at the ratio of 1:200 for live cell staining. The incubation with ITGA8 antibody was performed on ice for 1 hour, followed by wash with 1 ml FACS medium and incubation for 1 hour with fluorescence protein-conjugated secondary antibody (Donkey anti-Goat IgG (H+L) Highly Cross-Adsorbed Secondary Antibody, Alexa Fluor Plus 488 or 568, Thermo Scientific, # A11055 or A11057) protected from light. After that, another wash with 1 ml FACS medium was performed and the cell suspension was then subjected to FACS to sort out ITGA8^+^ hNPC through BD FACSAria™ III Cell Sorter. Purified ITGA8^+^ hNPCs (70-90% SIX2^+^/PAX2^+^) were cultured with the same protocol as described for purified *SIX2*-GFP; *PAX2*-mCherry iNPCs. After 2 to 3 passages in hNPSR-v2 medium, stable hNPC line with 90-95% hNPCs were established.

### Lentiviral expression of mCherry in iNPCs

30,000 *SIX2-*GFP^+^*/PAX2-*mCherry^+^ iNPCs were seeded into one well in 24-well plate. 24 hours later, iNPCs were infected with lentivirus expressing mCherry driven by EF1α promoter (EF1a_mCherry_P2A_Hygro, Addgene, # 135003) using spinfection method described above. 24 hours after infection, 150 μg/ml Hygromycin B (Thermo Scientific, Cat. No. 10687010) was added to the medium to select for iNPCs that have been successfully infected. More than 95% of the iNPCs showed bright mCherry expression after 1 week of Hygromycin B selection. (Note that exogenous mCherry is much brighter than endogenous mCherry from the *PAX2*-mCherry reporter, allowing the separation of these two mCherry signals.)

### Targeted genome editing at the *AAVS1* loci in iNPCs

30,000 *SIX2-*GFP^+^*/PAX2-*mCherry^+^ iNPCs were seeded into one well in 24-well plate. 24 hours later, iNPCs were transfected with a mixture of two plasmids that provide donor DNA for targeted knockin of CAG promoter-driven mCherry expression cassette at the AAVS1 loci (pAAVS1-P-CAG-mCherry, Addgene, # 80492), and that express Cas9 and sgRNA (pXAT2, Addgene, # 80494), at the ratio of 3:1, using Lipofectamine 3000 transfection reagent. 24 hours after transfection, 0.3 μg/ml puromycin was added to the medium to select for iNPCs that have been successfully gene edited. More than 97% of iNPCs showed bright mCherry expression after 1 week of puromycin selection. (Note that exogenous mCherry is much brighter than endogenous mCherry from the *PAX2*-mCherry reporter, allowing the separation of these two mCherry signals.)

### Generation of human nephron organoids from human NPC lines

iNPCs or hNPCs cultured in hNPSR-v2 medium were dissociated into single cells using Accumax as described above. 30,000 cells were seeded into one well of a U-bottom 96-well plate (Thermo Fisher Scientific, # 174929) and cultured with 100 μl hNPSR-v2 medium overnight for cells to aggregate. On the next day (Day 0), 3D NPC aggregates were transferred onto 6-well format transwell membrane (Corning, # 3450) with 6 μM CHIR99021 in 1.2 mL STEMdiff™ APEL™2 medium (STEMCELL Technologies, # 05270) at the bottom of the chamber for 1-hour. Then medium on the bottom chamber was removed as much as possible, and 1.2mL STEMdiff™ APEL™2 medium with 50 ng/mL FGF9 and 1 μg/mL heparin was added for continuous culture. This medium was refreshed every other day. On Day 5, medium was changed to 1.2mL STEMdiff™ APEL™2 medium without any other factors. The medium was refreshed every other day till Day 14, when the samples were harvested for various assays.

### Growth curve analysis of NPCs cultured in mNPSR-v2 or hNPSR-v2 media

For the growth curve analysis, 5,000 mNPCs, iNPCs, or hNPCs were seeded into each well of a 96-well plate coated with Matrigel and cultured in their corresponding mNPSR-v2 or hNPSR-v2 media. For each experiment, 16 wells were prepared for each line tested. Every 24 hours, 4 wells were dissociated, and the cell number was counted till 4 days after initial cell seeding, to determine the cell growth from day 0 to day 4. Three independent experiments were performed for each NPC line, serving as three biological replicates, to calculate the average cell numbers and the error bars at each time point.

### Bulk RNA sequencing

Cultured mouse and human NPC samples were collected and lysed in TRIzol reagent or DNA/RNA Shield and stored at -80°C. Total RNA was extracted using Direct-zol RNA MicroPrep Kit (Zymo) or Quick-RNA Microprep Kit (Zymo). Bulk RNA-Sequencing was then performed through the Molecular Pathology Genomics Core of Children’s Hospital Los Angeles (CHLA), or through Novogene USA Inc. For sequencing through CHLA, cDNA library was prepared using KAPA Stranded mRNA-Seq Kit (KAPA Biosystems) and sequenced using Illumina HiSeq 2500. For sequencing through Novogene, library prep and sequencing were performed using standard Novogene bulk RNA-seq pipeline.

### Bulk RNA-seq data analysis

RNA sequencing data was analyzed using Partek Flow Genomic Analysis Software. In addition to the new data generated from mouse and human NPC lines described in this study, legendary bulk RNA-seq data of primary mouse and human NPCs were used as positive controls, and primary kidney cells that are not NPCs were used as negative controls. These include E12.5 Six2-negative primary mouse non-NPCs, E11.5, E12.5, E13.5, E16.5 and P0 primary mouse NPCs ^1^, SIX2-negative primary human non-NPCs, SIX2+ human NPCs, SIX2+/MEIS1+ human NPCs, ^6^ and ITGA8+ human NPCs ^3^. FASTQ files were trimmed from both ends based on a minimum read length of 25 bps and an end minimum quality score (Phred) of 20 or higher. For mouse samples, reads were aligned to mm39 using STAR 2.7.8a. For human samples, reads were aligned to hg38 using STAR 2.7.8a. Aligned reads were quantified to the Partek E/M annotation model. Gene counts were normalized using DESeq2 Median ratio. DESeq2 Median ratio values can be found in Table S4 (mouse NPCs) and Table S13 (human NPCs). 972 mouse NPC signature genes were identified by comparing primary Six2^+^ E12.5 NPCs and primary Six2^-^ E12.5 non-NPCs^1^, with cutoff of fold change > 1.5 or < -1.5, and p-value < 0.01 using DESeq2 (Table S4). 1058 human NPC signature genes were identified by comparing 5 datasets of primary ITGA8^+^, SIX2^+^ or SIX2^+^/MEIS1^+^ NPCs with 2 datasets of primary SIX2^-^ non-NPCs^2, 3^, with cutoff of fold change > 1.5 or < -1.5, and FDR < 0.05 using DESeq2 (Table S13). Principle component analysis (PCA) was performed using the mouse or human NPC signature genes identified. Hierarchical clustering of selected genes was produced based on the genes’ DESeq2 Median ratio values, clustering samples and features with average linkage cluster distance and Euclidean point distance.

### Genome-wide CRISPR screen

The Brie genome-wide CRISPR knockout library^7^ (Addgene # 73632) was introduced to cultured NPCs via lentiviral infection in two different NPC lines as two biological replicates. This library contains 4 sgRNAs for each of the 19,674 protein-coding genes of the mouse genome, and 1,000 non-targeting control sgRNAs, totaling 78,637 unique sgRNAs (Table S5). The infection was carried out at a low multiplicity of infection (MOI) of 0.3, to ensure the majority of the NPCs express only one sgRNA. For each experiment, twenty-five million NPCs were used for the initial infection so that at least 100 cells would carry the same sgRNA to protect against random loss of sgRNA if lower cell number was used. 0.3 µg/ml puromycin was added 48 hours after lentiviral infection, to select for the successfully infected cells, which were further cultured continuously for a total of 3 weeks since lentiviral infection. Genomic DNA was extracted after 3 weeks of culture, and targeted PCR was performed to amplify the sgRNA integrated into the genome for next-generation sequencing, following Sequencing Protocol provided by Addgene (“Broad Institute PCR of sgRNAs for Illumina sequencing”). Next-generation sequencing was performed from the Molecular Pathology Genomics Core of Children’s Hospital Los Angeles using Illumina HighSeq 2500.

### Genome-wide CRISPR screen data analysis

Normalized read counts for each individual sgRNA in the plasmid library, before and after CRISPR screen (Table S5), CRISPR screen beta scores (Table S6), and the scatterplots of beta scores, were generated using MAGeCKFlute ^8^. 1798 genes from CRISPR screen replicate #1, and 1627 genes from CRISPR screen replicate #2, with beta scores > 1.5 or < -1.5, and p-values < 0.05, were identified as potential hits (Table S7) for further analyses using Canonical Pathway Analysis tools of the Ingenuity Pathway Analysis (IPA) platform (Table S8). Full list of reference FGF, Wnt, and LIF signaling pathway-related genes, and the genes identified in the screens, were summarized in Table S9. Full list of NPC-specific genes identified by comparisons between primary E12.5 NPC and primary E12.5 non-NPC^1^, or between primary E16.5 NPC and primary E16.5 IPC^2^, and the 64 highly confident NPC self-renewal hits, were summarized in Table S10. Full list of reference CAKUT or Wilms tumor-related genes, and the genes identified in the screens, were summarized in Table S11.

### KMT2A and KAT6A inhibitor treatment experiments

20,000 mNPCs were seeded into each Matrigel-coated well of 96-well plates, and cultured in (1) mNPSR-v2 + DMSO, (2) mNPSR-v2 + 100 nM WDR5 degrader, (3) mNPSR-v2 + 1 µM MLL1 (inh), (4) mNPSR-v2 + 5 µM MOZ-IN-3, or (5) mNPSR-v2 + 5 µM WM-1119. Medium was changed every two days and samples were harvested for immunofluorescence staining and qRT-PCR assay on day 8. Three independent experiments were conducted for each group as biological replicates.

### Establishment of one-step multiplexed CRISPR/Cas9 knockout lentiviral plasmids

Three different sgRNAs (sgRNA-A, sgRNA-B, and sgRNA-C) were designed to target *EGFP* coding sequences^9^, or the first exons of *Pkd1*, or *Pkd2 genes*. These sgRNAs were inserted individually into lentiGuide-Puro plasmid (Addgene # 52963) following the cloning protocols provided from the plasmid depositor. Then, for each target gene, the three sgRNA expression cassettes were subcloned one by one from the three lentiGuide-Puro plasmids into a modified pLKO.1-TRC plasmid (additional multiple cloning site BclI-EsrGI-MluI-NheI-PstI-SalI-XbaI-XmaI was inserted between the original PpuMI and EcoRI sites in the pLKO.1-TRC plasmid (Addgene # 10878)) to make tandem sgRNA expression cassettes in the same lentiviral vector.

sgRNA targeting sequences:

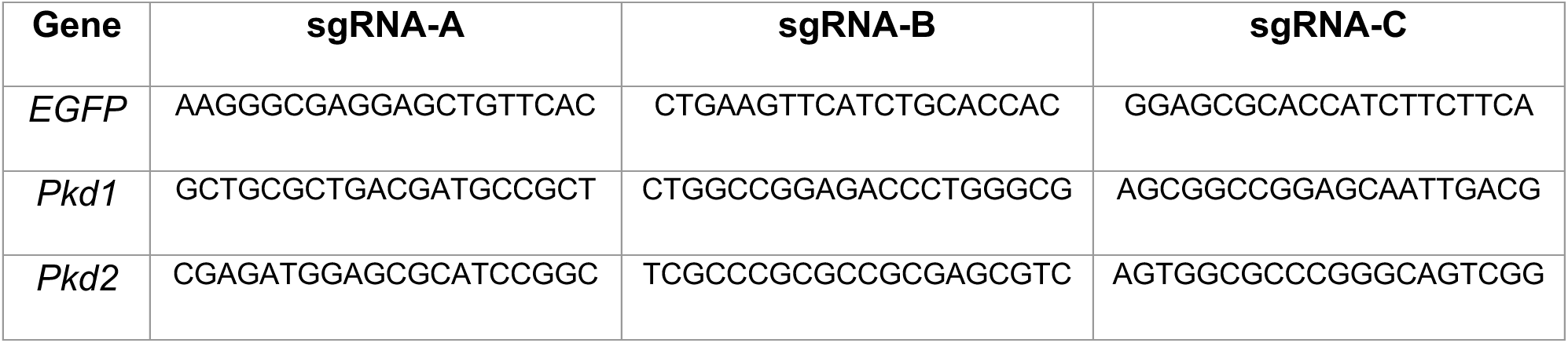

### One-step gene knockout in mouse NPCs with multiplexed CRISPR/Cas9 KO system

Multiplexed CRISPR/Cas9 KO plasmids targeting *EGFP*, *Pkd1,* or *Pkd2* were generated as described above. Lentivirus was produced from these plasmids and used to infect mNPCs. For that, lentivirus was first packaged following protocols we described previously^1^ and was then concentrated 100x using Lenti-X Concentrator kit (Takara, # 631231). Concentrated lentivirus was aliquoted and stored in -80°C before use. The lentivirus was used at 1x final concentration together with 10 μg/ml polybrene (Sigma-Aldrich, Cat. No. TR-1003-G) diluted in mNPSR-v2. Lentiviral infection was conducted in mNPC cultured in wells of 96-well plate. (1) For gene editing in *Cas9-*EGFP mNPC line, used culture medium was removed from each well and 100 µl lentivirus-polybrene-mNPSR-v2 mixture was added to the well. The plate was then centrifuged at 800 g for 15 minutes at room temperature. After the spinfection, the lentivirus-polybrene-mNPSR-v2 mixture was removed and the infected mNPCs were washed three times gently with 150-200 µl pre-warmed PBS, then cultured in 100 µl fresh mNPSR-v2 medium. 24 hours after infection, 0.3 μg/ml puromycin was added to the medium to select for NPCs that have been successfully infected. (2) For gene editing in *Six2-*GFP mNPC lines or wildtype mNPC lines cultured in 96-well plate, cells were first infected with lentivirus containing lentiCas9-Blast plasmid (Addgene, # 52962) using similar spinfection protocol described above. 24 hours after infection, 2 μg/ml blasticidin was added to the medium to select for NPCs that have been successfully infected. 1 week after selection, these Cas9-expressing mNPCs were infected with lentivirus expressing multiplexed CRISPR/Cas9 KO system as described above to generate gene KO starting from the wild-type mNPC line. Once bulk *Pkd1^-/-^* or *Pkd2^-/-^* mNPC lines were generated, single cell clonal *Pkd1^-/-^* or *Pkd2^-/-^*mNPC lines were generated following the same protocols we described above for generating clonal mNPC lines from mNPCs.

### Genotyping of *Pkd1^-/-^* or *Pkd2^-/-^* single cell clonal mNPC lines

Genomic DNA of *Pkd1^-/-^*or *Pkd2^-/-^* single cell clonal mNPC lines were extracted using QuickExtract^TM^ DNA Extraction Solution (Lucigen, # QE09050). PCR primers were designed to flank the sgRNA targeting sites of *Pkd1* or *Pkd2* genes (listed below). PCR were performed and PCR amplicons were used to conduct gel running and purified by DNA Clean & Concentrator kit (Zymo Research, # D4031). A-tailing were then performed on the purified PCR products following the NEB protocol: https://www.neb.com/protocols/2013/11/01/a-tailing-with-taq-polymerase. A-tailed PCR products were ligated with digested linear pGEM®-T Vector (Promega, Cat. No. A3600), followed by transformation with One Shot™ TOP10 Chemically Competent E. coli (Thermo Fisher Scientific, # C404010). Transformed E. coli were evenly daubed on IPTG/X-gal/Amp agarose plates and incubated at 37°C overnight. White bacterial clones were then picked up, inoculated, followed by mini-prep plasmid extraction (Zymo Research, # D4020). Extracted plasmids were sent to for Sanger sequencing. SnapGene software was utilized to analyze sequencing data.

Primers for PCR-based *Pkd1^-/-^* or *Pkd2^-/-^* genotyping:

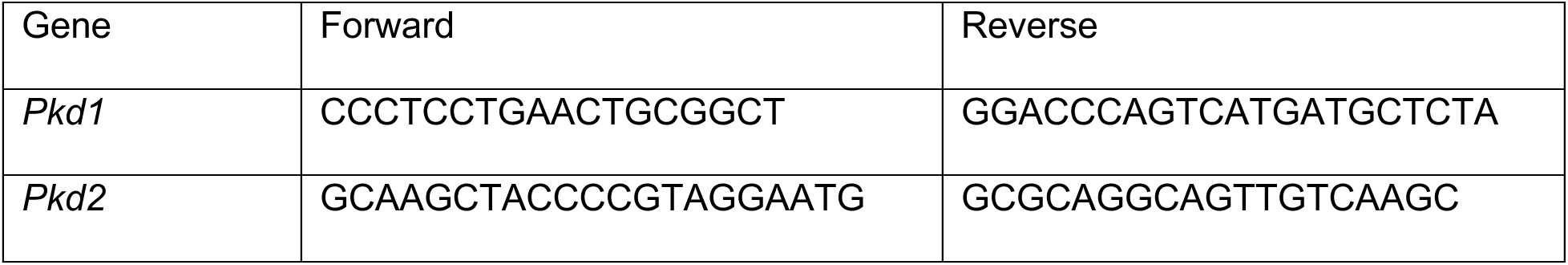

### Optimization of mini *Pkd2^-/-^* cystic organoid generation

Clonal *Pkd2^-/-^*mNPC lines and *GFP^-/-^* mNPC lines were used to optimize mini organoids model. For the first round of optimization, after mNPSR-v2 cultured NPCs reached 80-90% confluency in 2D culture, cells were dissociated into single cells using Accumax Cell Dissociation Solution. 500,000 cells were seeded into each well of AggreWell™800 24-well Plate (STEMCELL Technologies, Cat. No. 34850) with 2 mL mNPSR-v2 medium and cultured overnight. On the next day (Day 0), mini 3D NPC aggregates formed, and they were transferred into 12-well plates with around 30 mini aggregates/well in 1mL medium. mini 3D NPC aggregates were treated with different concentrations of CHIR99021 at 3.0 μM, 4.5 μM or 6.0 μM in KR5 or hBI medium under shaking culture at 120 rpm (VWR Orbital Shaker Model 1000) for 2 days (D0-D2), and then followed by culture with KR5 or hBI for 5 days (D2-D7). For the second round of optimization, mini 3D NPC aggregates were transferred into 12-well plates or ultra-low attachment plates under shaking culture or suspension culture, and treated with CHIR99021 at 4.5 μM in KR5 medium with 1% MTG or without MTG for 2 days (D0-D2), and then followed by culture in KR5 with 1% MTG or without MTG for 5 days (D2-D7). Cystic organoid formation efficiency was quantified based on 3-4 independent experiments and cyst diameters were measured by the measure tools of Olympus Cellsens Standard software. See also Figure S6 for more information.

#### hBI medium

Basal medium: DMEM/F12 (1:1) (1X), Invitrogen, Cat. No. 11330-032.

Supplements:

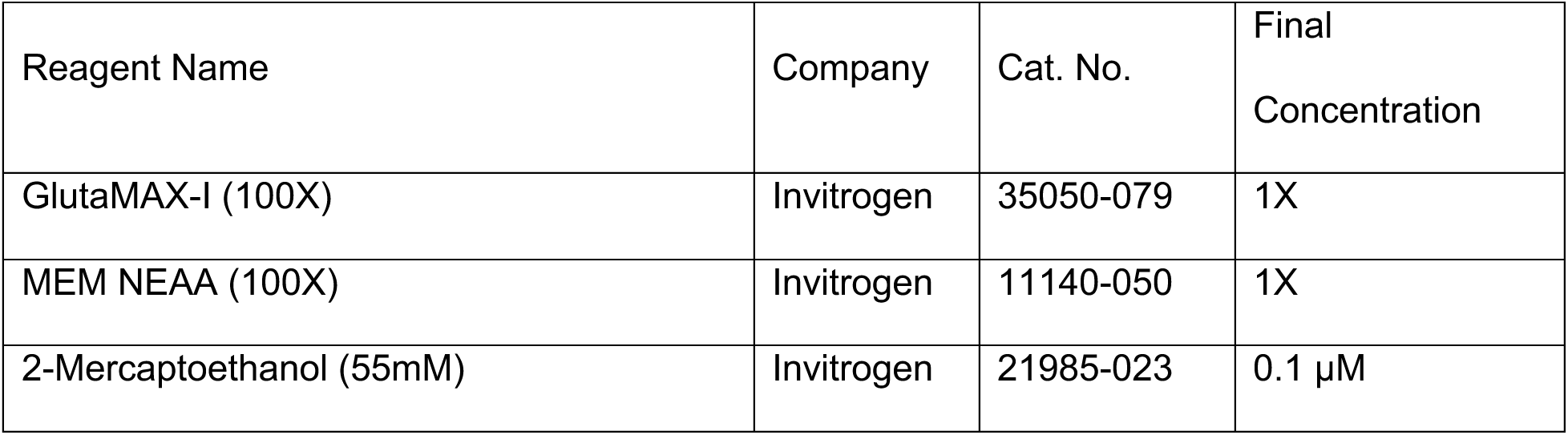
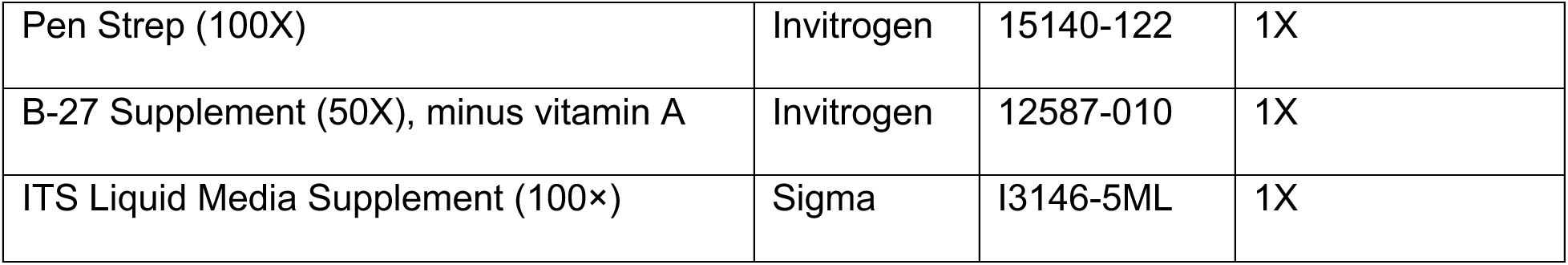

### Generation of PKD organoid models from mouse and human NPC lines

#### Traditional PKD organoid models

Clonal *Pkd2^-/-^* mNPC lines or *PKD2^-/-^* iNPC lines were used to generate corresponding mouse or human ADPKD models. The protocols for generation of mouse and human nephron organoids on transwell plates from NPC lines were described above. On day 7 (mouse) or day 14 (human) of organoid differentiation, each organoid was cut into 6 pieces using needles and subjected to shaking culture at 120 rpm (VWR Orbital Shaker Model 1000) in KR5 medium to develop cysts. Cystic organoid formation efficiency was quantified based on 3-4 independent experiments and cyst diameters were measured by the measure tools of Olympus Cellsens Standard software. For drug testing using this traditional PKD organoid model, for each drug treatment group, 30 small organoid pieces from fully developed mouse or human nephron organoids were pooled together in one well of a 12-well plate, cultured in KR5 medium with shaking, and supplemented with the drug to be tested. Drugs were freshly added to the medium and were refreshed with KR5 medium every two days. 4 days after drug treatment, cystic organoids were recorded through imaging and cyst formation efficiency and cyst diameter were quantified. Concentrations of drugs used in this study were determined based on previous publications ^10–13^. Cystic organoid formation efficiency was quantified based on 3-4 independent experiments and cyst diameters were measured by the measure tools of Olympus Cellsens Standard software.

#### Scalable mini PKD organoid models

Clonal *Pkd2^-/-^* mNPC lines or *PKD2^-/-^* iNPC lines were used to generate corresponding mouse or human ADPKD mini organoid models. For that, 500,000 mNPCs or iNPCs were seeded into each well of AggreWell™800 24-well Plate (STEMCELL Technologies, # 34850) with 2 mL mNPSR-v2 medium (mouse) or hNPSR-v2 medium (human), and cultured overnight to form mini aggregates (∼300 mini aggregates per well, or ∼7,200 mini aggregates per plate). On the next day (Day 0), mini 3D NPC aggregates from one well were transferred into 2 wells of 6-well plate with 2.5 mL KR5-CF medium in each well with shaking at 120 rpm. These aggregates were treated with KR5-CF medium for 2 days (mouse) or 3 days (human), then the medium was changed to 2.5 mL KR5 medium with medium refreshed every other day till harvested. Obvious PKD cysts typically emerged 4 days (mouse) or 8 days (human) after shaking culture. Cystic organoid formation efficiency was quantified based on 3-4 independent experiments and cyst diameters were measured by the measure tools of Olympus Cellsens Standard software.

### Drug screening with scalable mini PKD organoid models

Commercially available small molecule library (Cayman, # 11076) targeting major epigenetic processes was used for drug screening in the scalable mini PKD organoid models. Following protocols described above, thousands of mini NPC aggregates were generated using a full plate of AggreWell™800 24-well Plate starting from 12 million *Pkd2*^-/-^ clonal NPC line #4, or #5, as two biological replicates. These mini NPC aggregates were seeded into 12-well plates with around 30 mini NPC aggregates per well (Day 0). These plates were then subjected to shaking culture with 1 mL KR5-CF medium per well for 2 days. Medium was changed to 1 mL KR5 medium per well after 2 days (Day 2). On Day 3, small molecules from the epigenetic library were added individually into each well at the concentration of 1 µM, with controls that have DMSO, or different concentrations of metformin or tolvaptan. Cystic organoid percentages and cyst diameters were quantified on Day 5 as described above. Similar protocols were used when individual hits were further validated rather than from the initial screen. For drug testing in the human mini PKD organoid models, the KR5-CF step was extended to 3 days (Day 0 to Day 3), followed by KR5 step with medium refreshed every two days. Drug candidates were added on Day 6, and data were analyzed on Day 8. Cystic organoid formation efficiency was quantified based on 3-4 independent experiments and cyst diameters were measured by the measure tools of Olympus Cellsens Standard software.

### Seahorse assays on *GFP^-/-^* or *Pkd2^-/-^* NPCs and *GFP^-/-^* or *Pkd2^-/-^* nephron organoids

The XFp Extracellular Flux Analyzer (Agilent) was used for extracellular flux measurements. For *GFP^-/-^* mNPCs vs *Pkd2^-/-^* mNPCs, 20,000 cells per well were seeded in iMatrix-511 (Nacalai USA, # 892021) coated Seahorse miniplates 18-20 hours before the assay. For *GFP^-/-^* nephron organoids vs *Pkd2^-/-^* nephron organoids, on Day 4 of shaking culture following the mini PKD organoid protocol, one single nephron organoid was seeded into each well of poly-l-lysine coated Seahorse miniplate 1 hour before the assay. The XF sensor cartridges were hydrated overnight at 37°C without carbon dioxide in the Seahorse XF calibrant solution, as recommended by the manufacturer protocol. For the cell energy phenotype test, cell culture medium was replaced with XF assay medium (unbuffered DMEM [pH7.4] with 17.5 mM glucose, 0.5 mM pyruvate, and 4.5 mM glutamine). Microplate with cells was placed in a 37°C incubator without carbon dioxide for one hour. Oligomycin and FCCP were added to the ports at the final concentration of 1 µM. Standard XFp cell energy phenotype test (3 cycles of baseline measurements and 5 cycles of Oligomycin + FCCP with 3 minutes mixing and 3 minutes measuring) were performed for 1hour. Measurements were recorded from three to six independent experiments with three technical replicates per group in each independent experiment. Data were either normalized to cell numbers for NPCs (stained and counted with Hoechst 33342), or normalized to total genomic DNA amounts for organoids (CyQUANT® Cell Proliferation Assay, Invitrogen, C7026). Data were analyzed using Wave software.

### Immunofluorescence staining

#### Whole-mount staining

Samples were fixed in 4% PFA (4% Paraformaldehyde Aqueous Solution, Electron Microscopy Sciences, #157-4) for 15 minutes (kidney organoids, cystic organoids, or engineered kidneys) in Eppendorf tubes or tissue culture plates at room temperature. They were then washed four times in 1X PBS (Corning, Cat. No. 21-040-CV) for total 30 minutes. After the washes, samples were blocked in blocking solution (0.3% PBST containing 3% BSA) for 30 minutes at room temperature or 4°C overnight followed by primary antibody staining (primary antibodies were diluted in blocking solution) at 4°C overnight. On the second day, samples were washed four times with 0.3% PBST for total 60 minutes at room temperature. Secondary antibodies diluted in blocking solution were added and samples were incubated at 4°C overnight. On the third day, samples were washed four times with 0.3% PBST for total 60 minutes at room temperature then mounted for imaging. Confocal Microscope Zeiss LSM 800: AxioObserver.M2 was used for imaging recording in this study.

#### Cryo-section staining

Samples were fixed in 4% PFA for 30 minutes (human fetal kidneys, mouse embryonic kidneys, or mouse postnatal kidneys) in Eppendorf tubes or tissue culture plates at room temperature and then washed four times in 1X PBS for total 30 minutes. Fixed kidneys were transferred and incubated in 30% sucrose overnight at 4°C. After these kidneys sunk to the bottom in the sucrose on the next day, they were then transferred into a plastic mold and embedded in OCT Compound (Scigen, Cat. No. 4586K1) and froze in -80°C for 24 hours to make a cryo-block. The cryo-blocks were sectioned using Leica CM1800 Cryostat. For staining, these sectioned slides were blocked with blocking solution for 30 minutes followed by 2 hours of primary antibodies staining, all at room temperature. After primary staining, wash the slides four times with 0.3% PBST for a total of 15 minutes, then secondary staining for one hour at room temperature. After secondary staining, the slides were washed four times with 0.3% PBST for a total of 15 minutes and mounted with mounting medium (Southern Biotech, Fluoromount-G® Mounting Medium, #0100-01). For the Cryo-section staining of human fetal kidneys and mouse embryonic kidneys with p-p38, p-Smad2 or p-Smad1/5/9 antibodies, we used TSA-based immunocytofluorescence staining with TSA Plus Cyanine 3 Evaluation Kit (PerkinElmer, # NEL744E001KT) to enhance the phosphorylation signals. Slides were first incubated with 3% H_2_O_2_ reagent at room temperature for 10 minutes, rinsed with PBS, then blocked for 30 mins with blocking buffer. After blocking, these slides were incubated with diluted primary antibody overnight at 4°C (p-Smad2 dilution ratio-1:4000, p-Smad1/5/9 dilution ratio-1:4000, p-p38 dilution ratio-1:8000, in blocking buffer). On the second day, slides were rinsed with 0.3% PBST, incubated with anti-rabbit or anti-mouse HRP (1:2,000 in blocking buffer) for 2 hours at room temperature, rinsed again with 0.3% PBST, then incubated with TSA working solution (1:100 dilution) for 15 minutes at room temperature followed by a final rinse with 0.3% PBST. After the completion of TSA-based staining, proceed to standard IF staining.

#### Immunofluorescence staining in 96-well plates

NPCs cultured in 96-well plates were stained directly in the plates to determine various NPC marker gene expression. For that, the used culture medium was first removed from each well and 50 µl 4% PFA was added to fix the samples in the plates for 10 minutes at room temperature. Fixed samples were then gently washed three times in 1X PBS (Corning, Cat. No. 21-040-CV) 3 times for total 15 minutes, blocked in 100 µl blocking solution (0.1% PBST containing 3% BSA) for 30 minutes at room temperature, then followed by primary antibody staining at room temperature for 2 hours. Then, samples were gently washed two times with PBST for 10 minutes and secondary staining was conducted for one hour at room temperature. After the secondary staining, samples were gently washed three times with PBST for 15 minutes, and the PBST from the last wash was kept in the well to prevent samples from drying out. Samples were then ready for observation and recording. Note: NPCs cultured on Matrigel-coated 96-well plates can easily detach from the plate during staining, it is important to add/remove reagents gently in the process of staining. Validated primary antibodies in this study can be found in “Key Resources Table”.

#### Imaging data quantification

For immunostaining quantification, 3-4 different fields of view were randomly selected to count the number of positively stained cell numbers and total cell numbers (as determined by DAPI signals). At least 500 cells per field of view were included. Error bars represent standard derivation between different fields of views.

### RNA isolation, reverse transcription, qRT-PCR, and immunoblotting

Samples were dissolved in 100 µl TRIzol (Invitrogen, Cat. No. 15596018) or 100 µl DNA/RNA Shield (Zymo Research, Cat. No. R1100-50) and kept in -80°C freezer. RNA isolation was performed using the Direct-zol RNA MicroPrep Kit (Zymo Research, Cat. No. R2062) or Quick-RNA Microprep Kit (Zymo Research, Cat. No. R1051) according to the manufacturer’s instructions. Reverse transcription was performed using the iScript Reverse Transcription Supermix (Bio-Rad, Cat. No. 1708841) following the manufacturer’s instructions. qRT-PCR was performed using SsoAdvanced Universal SYBR® Green Supermix (Bio-Rad, Cat. No. 1725274) or AzuraView GreenFast qPCR Blue Mix LR (Azura Genomics, Cat. No. AZ-2320) and carried out on an Applied Biosystems Vii 7 RT-PCR system (Thermo Scientific P/N 4453552). Validated gene-specific primers in this study can be found in Table S14. Fold changes were calculated from ΔCt using Gapdh as a housekeeping gene as previously described^14^. Immunoblotting experiments were performed as described previously^14^.

## FACS

Cells were dissociated/prepared as described above. FACS sorting was performed on a BD FACSAria™ III Cell Sorter, operated by experienced core facility staff at USC Stem Cell’s FACS core. Sorted cells were collected in 1.5 ml Eppendorf tubes with 500 µl dissection medium on ice.

## QUANTIFICATION AND STATISTICAL ANALYSIS

Data are presented as mean ± SD from at least three biological replicates (n=3). Statistical significance was determined by two-tailed unpaired Student’s t tests; ns, not significant; *, p<0.05; **, p<0.01; ***, p<0.001.

## REFERENCES

1 McMahon, A. P. Development of the Mammalian Kidney. Curr Top Dev Biol 117, 31–64 (2016). https://doi.org:10.1016/bs.ctdb.2015.10.010

2 Schnell, J., Achieng, M. & Lindström, N. O. Principles of human and mouse nephron development. Nat Rev Nephrol (2022). https://doi.org:10.1038/s41581-022-00598-5

3 Kobayashi, A. et al. Six2 defines and regulates a multipotent self-renewing nephron progenitor population throughout mammalian kidney development. Cell Stem Cell 3, 169–181 (2008). https://doi.org:10.1016/j.stem.2008.05.020

4 van der Ven, A. T., Vivante, A. & Hildebrandt, F. Novel Insights into the Pathogenesis of Monogenic Congenital Anomalies of the Kidney and Urinary Tract. J Am Soc Nephrol 29, 36–50 (2018). https://doi.org:10.1681/ASN.2017050561

5 Treger, T. D., Chowdhury, T., Pritchard-Jones, K. & Behjati, S. The genetic changes of Wilms tumour. Nat Rev Nephrol 15, 240–251 (2019). https://doi.org:10.1038/s41581-019-0112-0

6 Taguchi, A. et al. Redefining the in vivo origin of metanephric nephron progenitors enables generation of complex kidney structures from pluripotent stem cells. Cell Stem Cell 14, 53–67 (2014). https://doi.org:10.1016/j.stem.2013.11.010

7 Freedman, B. S. et al. Modelling kidney disease with CRISPR-mutant kidney organoids derived from human pluripotent epiblast spheroids. Nat Commun 6, 8715 (2015). https://doi.org:10.1038/ncomms9715

8 Morizane, R. et al. Nephron organoids derived from human pluripotent stem cells model kidney development and injury. Nat Biotechnol 33, 1193–1200 (2015). https://doi.org:10.1038/nbt.3392

9 Takasato, M. et al. Kidney organoids from human iPS cells contain multiple lineages and model human nephrogenesis. Nature 526, 564–568 (2015). https://doi.org:10.1038/nature15695

10 Brown, A. C., Muthukrishnan, S. D. & Oxburgh, L. A synthetic niche for nephron progenitor cells. Dev Cell 34, 229–241 (2015). https://doi.org:10.1016/j.devcel.2015.06.021

11 Tanigawa, S., Taguchi, A., Sharma, N., Perantoni, A. O. & Nishinakamura, R. Selective In Vitro Propagation of Nephron Progenitors Derived from Embryos and Pluripotent Stem Cells. Cell Rep 15, 801–813 (2016). https://doi.org:10.1016/j.celrep.2016.03.076

12 Li, Z. et al. 3D Culture Supports Long-Term Expansion of Mouse and Human Nephrogenic Progenitors. Cell Stem Cell 19, 516–529 (2016). https://doi.org:10.1016/j.stem.2016.07.016

13 Little, M. H. & Combes, A. N. Kidney organoids: accurate models or fortunate accidents. Genes Dev 33, 1319-1345 (2019). https://doi.org:10.1101/gad.329573.119

14 Wu, H. et al. Comparative Analysis and Refinement of Human PSC-Derived Kidney Organoid Differentiation with Single-Cell Transcriptomics. Cell Stem Cell 23, 869–881.e868 (2018). https://doi.org:10.1016/j.stem.2018.10.010

15 Combes, A. N., Zappia, L., Er, P. X., Oshlack, A. & Little, M. H. Single-cell analysis reveals congruence between kidney organoids and human fetal kidney. Genome Med 11, 3 (2019). https://doi.org:10.1186/s13073-019-0615-0

16 Vanslambrouck, J. M. et al. Enhanced metanephric specification to functional proximal tubule enables toxicity screening and infectious disease modelling in kidney organoids. Nat Commun 13, 5943 (2022). https://doi.org:10.1038/s41467-022-33623-z

17 Shalem, O. et al. Genome-scale CRISPR-Cas9 knockout screening in human cells. Science 343, 84–87 (2014). https://doi.org:10.1126/science.1247005

18 Wang, T., Wei, J. J., Sabatini, D. M. & Lander, E. S. Genetic screens in human cells using the CRISPR-Cas9 system. Science 343, 80–84 (2014). https://doi.org:10.1126/science.1246981

19 Doench, J. G. Am I ready for CRISPR? A user’s guide to genetic screens. Nat Rev Genet 19, 67–80 (2018). https://doi.org:10.1038/nrg.2017.97

20 Tanigawa, S. et al. Activin Is Superior to BMP7 for Efficient Maintenance of Human iPSC-Derived Nephron Progenitors. Stem Cell Reports 13, 322–337 (2019). https://doi.org:10.1016/j.stemcr.2019.07.003

21 Li, Z., Araoka, T. & Belmonte, J. C. I. Gene Editing in 3D Cultured Nephron Progenitor Cell Lines. Methods Mol Biol 1926, 151–159 (2019). https://doi.org:10.1007/978-1-4939-9021-4_13

22 Zeng, Z. et al. Generation of patterned kidney organoids that recapitulate the adult kidney collecting duct system from expandable ureteric bud progenitors. Nat Commun 12, 3641 (2021). https://doi.org:10.1038/s41467-021-23911-5

23 Garreta, E. et al. Fine tuning the extracellular environment accelerates the derivation of kidney organoids from human pluripotent stem cells. Nat Mater 18, 397–405 (2019). https://doi.org:10.1038/s41563-019-0287-6

24 Lindström, N. O. et al. Conserved and Divergent Features of Mesenchymal Progenitor Cell Types within the Cortical Nephrogenic Niche of the Human and Mouse Kidney. J Am Soc Nephrol 29, 806–824 (2018). https://doi.org:10.1681/ASN.2017080890

25 Lawlor, K. T. et al. Nephron progenitor commitment is a stochastic process influenced by cell migration. Elife 8 (2019). https://doi.org:10.7554/eLife.41156

26 Przybyla, L. & Gilbert, L. A. A new era in functional genomics screens. Nat Rev Genet 23, 89–103 (2022). https://doi.org:10.1038/s41576-021-00409-w

27 Doench, J. G. et al. Optimized sgRNA design to maximize activity and minimize off-target effects of CRISPR-Cas9. Nat Biotechnol 34, 184–191 (2016). https://doi.org:10.1038/nbt.3437

28 Li, W. et al. Quality control, modeling, and visualization of CRISPR screens with MAGeCK-VISPR. Genome Biol 16, 281 (2015). https://doi.org:10.1186/s13059-015-0843-6

29 Wang, B. et al. Integrative analysis of pooled CRISPR genetic screens using MAGeCKFlute. Nat Protoc 14, 756-780 (2019). https://doi.org:10.1038/s41596-018-0113-7

30 Makayes, Y. et al. Increasing mTORC1 Pathway Activity or Methionine Supplementation during Pregnancy Reverses the Negative Effect of Maternal Malnutrition on the Developing Kidney. J Am Soc Nephrol 32, 1898–1912 (2021). https://doi.org:10.1681/ASN.2020091321

31 Volovelsky, O. et al. Hamartin regulates cessation of mouse nephrogenesis independently of Mtor. Proc Natl Acad Sci U S A 115, 5998–6003 (2018). https://doi.org:10.1073/pnas.1712955115

32 Marrone, A. K. et al. MicroRNA-17∼92 is required for nephrogenesis and renal function. J Am Soc Nephrol 25, 1440–1452 (2014). https://doi.org:10.1681/ASN.2013040390

33 Ho, J. et al. The pro-apoptotic protein Bim is a microRNA target in kidney progenitors. J Am Soc Nephrol 22, 1053–1063 (2011). https://doi.org:10.1681/ASN.2010080841

34 Cargill, K. et al. Von Hippel-Lindau Acts as a Metabolic Switch Controlling Nephron Progenitor Differentiation. J Am Soc Nephrol 30, 1192–1205 (2019). https://doi.org:10.1681/ASN.2018111170

35 Nishinakamura, R. et al. Murine homolog of SALL1 is essential for ureteric bud invasion in kidney development. Development 128, 3105–3115 (2001).

36 Kanda, S. et al. Sall1 maintains nephron progenitors and nascent nephrons by acting as both an activator and a repressor. J Am Soc Nephrol 25, 2584–2595 (2014). https://doi.org:10.1681/ASN.2013080896

37 Basta, J. M., Robbins, L., Kiefer, S. M., Dorsett, D. & Rauchman, M. Sall1 balances self-renewal and differentiation of renal progenitor cells. Development 141, 1047–1058 (2014). https://doi.org:10.1242/dev.095851

38 Motamedi, F. J. et al. WT1 controls antagonistic FGF and BMP-pSMAD pathways in early renal progenitors. Nat Commun 5, 4444 (2014). https://doi.org:10.1038/ncomms5444

39 Motojima, M., Tanaka, M. & Kume, T. Foxc1 and Foxc2 are indispensable for the maintenance of nephron and stromal progenitors in the developing kidney. J Cell Sci 135 (2022). https://doi.org:10.1242/jcs.260356

40 Pan, X., Karner, C. M. & Carroll, T. J. Myc cooperates with β-catenin to drive gene expression in nephron progenitor cells. Development 144, 4173–4182 (2017). https://doi.org:10.1242/dev.153700

41 Xu, J. et al. Eya1 interacts with Six2 and Myc to regulate expansion of the nephron progenitor pool during nephrogenesis. Dev Cell 31, 434–447 (2014). https://doi.org:10.1016/j.devcel.2014.10.015

42 Chen, Z. et al. Jxc1/Sobp, encoding a nuclear zinc finger protein, is critical for cochlear growth, cell fate, and patterning of the organ of corti. J Neurosci 28, 6633–6641 (2008). https://doi.org:10.1523/JNEUROSCI.1280-08.2008

43 Tavares, A. L. P., Jourdeuil, K., Neilson, K. M., Majumdar, H. D. & Moody, S. A. Sobp modulates the transcriptional activation of Six1 target genes and is required during craniofacial development. Development 148 (2021). https://doi.org:10.1242/dev.199684

44 Atlasi, Y. & Stunnenberg, H. G. The interplay of epigenetic marks during stem cell differentiation and development. Nat Rev Genet 18, 643–658 (2017). https://doi.org:10.1038/nrg.2017.57

45 Avgustinova, A. & Benitah, S. A. Epigenetic control of adult stem cell function. Nat Rev Mol Cell Biol 17, 643–658 (2016). https://doi.org:10.1038/nrm.2016.76

46 Huang, B., Liu, Z., Vonk, A., Zeng, Z. & Li, Z. Epigenetic regulation of kidney progenitor cells. Stem Cells Transl Med 9, 655–660 (2020). https://doi.org:10.1002/sctm.19-0289

47 Liu, H. et al. Histone deacetylases 1 and 2 regulate the transcriptional programs of nephron progenitors and renal vesicles. Development 145 (2018). https://doi.org:10.1242/dev.153619

48 Denner, D. R. & Rauchman, M. Mi-2/NuRD is required in renal progenitor cells during embryonic kidney development. Dev Biol 375, 105–116 (2013). https://doi.org:10.1016/j.ydbio.2012.11.018

49 Liu, H. et al. The polycomb proteins EZH1 and EZH2 co-regulate chromatin accessibility and nephron progenitor cell lifespan in mice. J Biol Chem 295, 11542–11558 (2020). https://doi.org:10.1074/jbc.RA120.013348

50 Basta, J. M. et al. The core SWI/SNF catalytic subunit Brg1 regulates nephron progenitor cell proliferation and differentiation. Dev Biol 464, 176–187 (2020). https://doi.org:10.1016/j.ydbio.2020.05.008

51 van der Ven, A. T. et al. Whole-Exome Sequencing Identifies Causative Mutations in Families with Congenital Anomalies of the Kidney and Urinary Tract. J Am Soc Nephrol 29, 2348–2361 (2018). https://doi.org:10.1681/ASN.2017121265

52 Yu, X. et al. A selective WDR5 degrader inhibits acute myeloid leukemia in patient-derived mouse models. Sci Transl Med 13, eabj1578 (2021). https://doi.org:10.1126/scitranslmed.abj1578

53 Bergmann, C. et al. Polycystic kidney disease. Nat Rev Dis Primers 4, 50 (2018). https://doi.org:10.1038/s41572-018-0047-y

54 Cruz, N. M. et al. Organoid cystogenesis reveals a critical role of microenvironment in human polycystic kidney disease. Nat Mater 16, 1112–1119 (2017). https://doi.org:10.1038/nmat4994

55 Czerniecki, S. M. et al. High-Throughput Screening Enhances Kidney Organoid Differentiation from Human Pluripotent Stem Cells and Enables Automated Multidimensional Phenotyping. Cell Stem Cell 22, 929–940.e924 (2018). https://doi.org:10.1016/j.stem.2018.04.022

56 Kuraoka, S. et al. -Dependent Renal Cystogenesis in Human Induced Pluripotent Stem Cell-Derived Ureteric Bud/Collecting Duct Organoids. J Am Soc Nephrol 31, 2355–2371 (2020). https://doi.org:10.1681/ASN.2020030378

57 Shimizu, T. et al. A novel ADPKD model using kidney organoids derived from disease-specific human iPSCs. Biochem Biophys Res Commun 529, 1186–1194 (2020). https://doi.org:10.1016/j.bbrc.2020.06.141

58 Tran, T. et al. A scalable organoid model of human autosomal dominant polycystic kidney disease for disease mechanism and drug discovery. Cell Stem Cell 29, 1083–1101.e1087 (2022). https://doi.org:10.1016/j.stem.2022.06.005

59 Zuo, E. et al. One-step generation of complete gene knockout mice and monkeys by CRISPR/Cas9-mediated gene editing with multiple sgRNAs. Cell Res 27, 933–945 (2017). https://doi.org:10.1038/cr.2017.81

60 Platt, R. J. et al. CRISPR-Cas9 knockin mice for genome editing and cancer modeling. Cell 159, 440–455 (2014). https://doi.org:10.1016/j.cell.2014.09.014

61 Yang, B., Sonawane, N. D., Zhao, D., Somlo, S. & Verkman, A. S. Small-molecule CFTR inhibitors slow cyst growth in polycystic kidney disease. J Am Soc Nephrol 19, 1300–1310 (2008). https://doi.org:10.1681/ASN.2007070828

62 Takiar, V. et al. Activating AMP-activated protein kinase (AMPK) slows renal cystogenesis. Proc Natl Acad Sci U S A 108, 2462–2467 (2011). https://doi.org:10.1073/pnas.1011498108

63 Pastor-Soler, N. M. et al. Metformin improves relevant disease parameters in an autosomal dominant polycystic kidney disease mouse model. Am J Physiol Renal Physiol 322, F27–F41 (2022). https://doi.org:10.1152/ajprenal.00298.2021

64 Li, L. X. et al. Lysine methyltransferase SMYD2 promotes cyst growth in autosomal dominant polycystic kidney disease. J Clin Invest 127, 2751–2764 (2017). https://doi.org:10.1172/JCI90921

65 Li, L. X. et al. Cross talk between lysine methyltransferase Smyd2 and TGF-β-Smad3 signaling promotes renal fibrosis in autosomal dominant polycystic kidney disease. Am J Physiol Renal Physiol 323, F227–F242 (2022). https://doi.org:10.1152/ajprenal.00452.2021

66 Torres, V. E. et al. Tolvaptan in patients with autosomal dominant polycystic kidney disease. N Engl J Med 367, 2407–2418 (2012). https://doi.org:10.1056/NEJMoa1205511

67 Reif, G. A. et al. Tolvaptan inhibits ERK-dependent cell proliferation, Cl⁻ secretion, and in vitro cyst growth of human ADPKD cells stimulated by vasopressin. Am J Physiol Renal Physiol 301, F1005–1013 (2011). https://doi.org:10.1152/ajprenal.00243.2011

68 Shillingford, J. M. et al. The mTOR pathway is regulated by polycystin-1, and its inhibition reverses renal cystogenesis in polycystic kidney disease. Proc Natl Acad Sci U S A 103, 5466–5471 (2006). https://doi.org:10.1073/pnas.0509694103

69 Tao, Y., Kim, J., Schrier, R. W. & Edelstein, C. L. Rapamycin markedly slows disease progression in a rat model of polycystic kidney disease. J Am Soc Nephrol 16, 46–51 (2005). https://doi.org:10.1681/ASN.2004080660

70 Cai, J. et al. A RhoA-YAP-c-Myc signaling axis promotes the development of polycystic kidney disease. Genes Dev 32, 781–793 (2018). https://doi.org:10.1101/gad.315127.118

71 Trudel, M., D’Agati, V. & Costantini, F. C-myc as an inducer of polycystic kidney disease in transgenic mice. Kidney Int 39, 665–671 (1991). https://doi.org:10.1038/ki.1991.80

72 Padovano, V., Podrini, C., Boletta, A. & Caplan, M. J. Metabolism and mitochondria in polycystic kidney disease research and therapy. Nat Rev Nephrol 14, 678–687 (2018). https://doi.org:10.1038/s41581-018-0051-1

73 Rowe, I. et al. Defective glucose metabolism in polycystic kidney disease identifies a new therapeutic strategy. Nat Med 19, 488–493 (2013). https://doi.org:10.1038/nm.3092

74 Menezes, L. F., Lin, C. C., Zhou, F. & Germino, G. G. Fatty Acid Oxidation is Impaired in An Orthologous Mouse Model of Autosomal Dominant Polycystic Kidney Disease. EBioMedicine 5, 183–192 (2016). https://doi.org:10.1016/j.ebiom.2016.01.027

75 Lin, C. C. et al. A cleavage product of Polycystin-1 is a mitochondrial matrix protein that affects mitochondria morphology and function when heterologously expressed. Sci Rep 8, 2743 (2018). https://doi.org:10.1038/s41598-018-20856-6

76 Cao, Y. et al. Chemical modifier screen identifies HDAC inhibitors as suppressors of PKD models. Proc Natl Acad Sci U S A 106, 21819–21824 (2009). https://doi.org:10.1073/pnas.0911987106

77 Zhou, X. et al. Therapeutic targeting of BET bromodomain protein, Brd4, delays cyst growth in ADPKD. Hum Mol Genet 24, 3982-3993 (2015). https://doi.org:10.1093/hmg/ddv136

78 Zanconato, F., Cordenonsi, M. & Piccolo, S. YAP/TAZ at the Roots of Cancer. Cancer Cell 29, 783–803 (2016). https://doi.org:10.1016/j.ccell.2016.05.005

79 Yu, F. X., Zhao, B. & Guan, K. L. Hippo Pathway in Organ Size Control, Tissue Homeostasis, and Cancer. Cell 163, 811–828 (2015). https://doi.org:10.1016/j.cell.2015.10.044

80 Das, A. et al. Stromal-epithelial crosstalk regulates kidney progenitor cell differentiation. Nat Cell Biol 15, 1035–1044 (2013). https://doi.org:10.1038/ncb2828

81 Kastan, N. et al. Small-molecule inhibition of Lats kinases may promote Yap-dependent proliferation in postmitotic mammalian tissues. Nat Commun 12, 3100 (2021). https://doi.org:10.1038/s41467-021-23395-3

82 Subramanian, A. et al. Single cell census of human kidney organoids shows reproducibility and diminished off-target cells after transplantation. Nat Commun 10, 5462 (2019). https://doi.org:10.1038/s41467-019-13382-0

83 Phipson, B. et al. Evaluation of variability in human kidney organoids. Nat Methods 16, 79–87 (2019). https://doi.org:10.1038/s41592-018-0253-2

84 Tran, T. et al. In Vivo Developmental Trajectories of Human Podocyte Inform In Vitro Differentiation of Pluripotent Stem Cell-Derived Podocytes. Dev Cell 50, 102–116.e106 (2019). https://doi.org:10.1016/j.devcel.2019.06.001

85 Cebotaru, L. et al. Inhibition of histone deacetylase 6 activity reduces cyst growth in polycystic kidney disease. Kidney Int 90, 90–99 (2016). https://doi.org:10.1016/j.kint.2016.01.026

86 Kobayashi, A. et al. Identification of a multipotent self-renewing stromal progenitor population during mammalian kidney organogenesis. Stem Cell Reports 3, 650–662 (2014). https://doi.org:10.1016/j.stemcr.2014.08.008

87 Naiman, N. et al. Repression of Interstitial Identity in Nephron Progenitor Cells by Pax2 Establishes the Nephron-Interstitium Boundary during Kidney Development. Dev Cell 41, 349–365.e343 (2017). https://doi.org:10.1016/j.devcel.2017.04.022

88 Kreso, A. et al. Self-renewal as a therapeutic target in human colorectal cancer. Nat Med 20, 29–36 (2014). https://doi.org:10.1038/nm.3418

89 Jia, L., Zhang, W. & Wang, C. Y. BMI1 Inhibition Eliminates Residual Cancer Stem Cells after PD1 Blockade and Activates Antitumor Immunity to Prevent Metastasis and Relapse. Cell Stem Cell 27, 238–253.e236 (2020). https://doi.org:10.1016/j.stem.2020.06.022

90 Seeger-Nukpezah, T., Geynisman, D. M., Nikonova, A. S., Benzing, T. & Golemis, E. A. The hallmarks of cancer: relevance to the pathogenesis of polycystic kidney disease. Nat Rev Nephrol 11, 515–534 (2015). https://doi.org:10.1038/nrneph.2015.46

91 Bouchard, M., Souabni, A., Mandler, M., Neubüser, A. & Busslinger, M. Nephric lineage specification by Pax2 and Pax8. Genes Dev 16, 2958–2970 (2002). https://doi.org:10.1101/gad.240102

## SUPPLEMENTAL REFERENCES

1. Li, Z., Araoka, T., Wu, J., Liao, H.-K., Li, M., Lazo, M., Zhou, B., Sui, Y., Wu, M.-Z., and Tamura, I. (2016). 3D culture supports long-term expansion of mouse and human nephrogenic progenitors. Cell stem cell 19, 516–529.

2. Lindström, N.O., Guo, J., Kim, A.D., Tran, T., Guo, Q., De Sena Brandine, G., Ransick, A., Parvez, R.K., Thornton, M.E., Baskin, L., et al. (2018). Conserved and Divergent Features of Mesenchymal Progenitor Cell Types within the Cortical Nephrogenic Niche of the Human and Mouse Kidney. J Am Soc Nephrol 29, 806–824. 10.1681/ASN.2017080890.

3. O’Brien, L.L., Guo, Q., Lee, Y., Tran, T., Benazet, J.-D., Whitney, P.H., Valouev, A., and McMahon, A.P. (2016). Differential regulation of mouse and human nephron progenitors by the Six family of transcriptional regulators. Development 143, 595–608.

4. Zeng, Z., Huang, B., Parvez, R.K., Li, Y., Chen, J., Vonk, A.C., Thornton, M.E., Patel, T., Rutledge, E.A., and Kim, A.D. (2021). Generation of patterned kidney organoids that recapitulate the adult kidney collecting duct system from expandable ureteric bud progenitors. Nature communications 12, 1–15.

5. Morizane, R., Lam, A.Q., Freedman, B.S., Kishi, S., Valerius, M.T., and Bonventre, J.V. (2015). Nephron organoids derived from human pluripotent stem cells model kidney development and injury. Nat Biotechnol 33, 1193–1200. 10.1038/nbt.3392.

6. Lindström, N.O., Guo, J., Kim, A.D., Tran, T., Guo, Q., Brandine, G.D.S., Ransick, A., Parvez, R.K., Thornton, M.E., and Basking, L. (2018). Conserved and divergent features of mesenchymal progenitor cell types within the cortical nephrogenic niche of the human and mouse kidney. Journal of the American Society of Nephrology 29, 806–824.

7. Doench, J.G., Fusi, N., Sullender, M., Hegde, M., Vaimberg, E.W., Donovan, K.F., Smith, I., Tothova, Z., Wilen, C., Orchard, R., et al. (2016). Optimized sgRNA design to maximize activity and minimize off-target effects of CRISPR-Cas9. Nat Biotechnol 34, 184–191. 10.1038/nbt.3437.

8. Wang, B., Wang, M., Zhang, W., Xiao, T., Chen, C.-H., Wu, A., Wu, F., Traugh, N., Wang, X., and Li, Z. (2019). Integrative analysis of pooled CRISPR genetic screens using MAGeCKFlute. Nature protocols 14, 756–780.

9. Zuo, E., Cai, Y.J., Li, K., Wei, Y., Wang, B.A., Sun, Y., Liu, Z., Liu, J., Hu, X., Wei, W., et al. (2017). One-step generation of complete gene knockout mice and monkeys by CRISPR/Cas9-mediated gene editing with multiple sgRNAs. Cell Res 27, 933–945. 10.1038/cr.2017.81.

10. Low, J.H., Li, P., Chew, E.G.Y., Zhou, B., Suzuki, K., Zhang, T., Lian, M.M., Liu, M., Aizawa, E., and Esteban, C.R. (2019). Generation of human PSC-derived kidney organoids with patterned nephron segments and a de novo vascular network. Cell stem cell 25, 373–387. e379.

11. Tran, T., Song, C.J., Nguyen, T., Cheng, S.-Y., McMahon, J.A., Yang, R., Guo, Q., Der, B., Lindström, N.O., and Lin, D.C.-H. (2022). A scalable organoid model of human autosomal dominant polycystic kidney disease for disease mechanism and drug discovery. Cell Stem Cell 29, 1083–1101. e1087.

12. Li, L.X., Fan, L.X., Zhou, J.X., Grantham, J.J., Calvet, J.P., Sage, J., and Li, X. (2017). Lysine methyltransferase SMYD2 promotes cyst growth in autosomal dominant polycystic kidney disease. The Journal of clinical investigation 127, 2751–2764.

13. Takiar, V., Nishio, S., Seo-Mayer, P., King Jr, J.D., Li, H., Zhang, L., Karihaloo, A., Hallows, K.R., Somlo, S., and Caplan, M.J. (2011). Activating AMP-activated protein kinase (AMPK) slows renal cystogenesis. Proceedings of the National Academy of Sciences 108, 2462–2467.

14. Huang, B., Wang, B., Lee, W.Y.-W., Leung, K.T., Li, X., Liu, Z., Chen, R., cheng Lin, J., Tsang, L.L., and Liu, B. (2019). KDM3A and KDM4C regulate mesenchymal stromal cell senescence and bone aging via condensin-mediated heterochromatin reorganization. Iscience 21, 375–390.

