## Supplemental Information for "Modeling kidney development, disease, and plasticity with clonal expandable nephron progenitor cells and nephron organoids"

### Figure S1

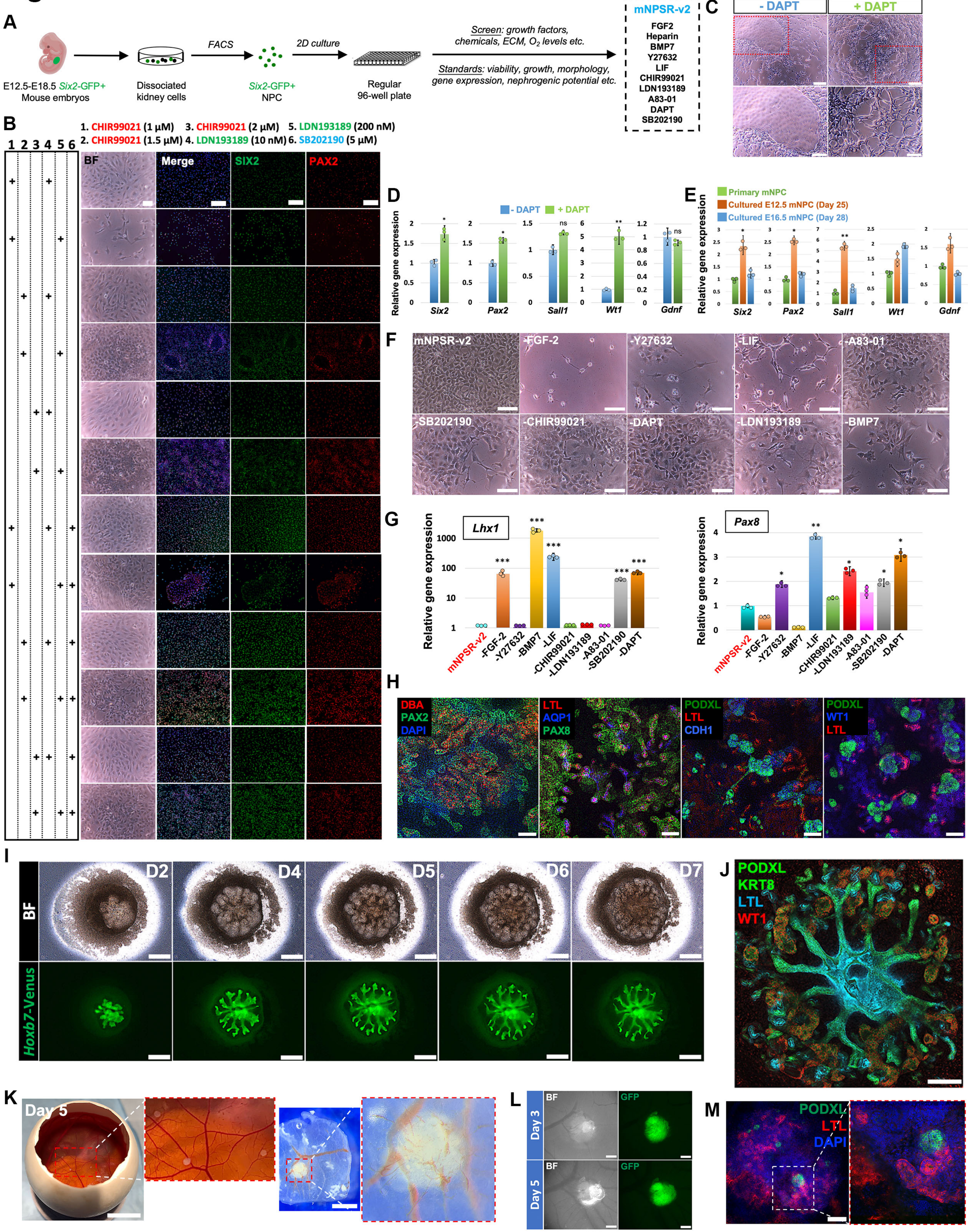
A
E12.5-E18.5 Six2-GFP+
Mouse embryos
Dissociated
kidney cells
FACS
Six2-GFP+
NPC
2D culture
Regular
96-well plate
Screen: growth factors,
chemicals, ECM, O2 levels etc.
Standards: viability, growth, morphology,
gene expression, nephrogenic potential etc.
mNPSR-v2
FGF2
Heparin
BMP7
Y27632
LIF
CHIR99021
LDN193189
A83-01
DAPT
SB202190
C
- DAPT
+ DAPT
B
1. CHIR99021 (1 µM) 3. CHIR99021 (2 µM) 5. LDN193189 (200 nM)
2. CHIR99021 (1.5 µM) 4. LDN193189 (10 nM) 6. SB202190 (5 µM)
BF
Merge
SIX2
PAX2
D
- DAPT
+ DAPT
Relative gene expression
Six2
Pax2
Sall1
Wt1
Gdnf
E
Primary mNPC
Cultured E12.5 mNPC (Day 25)
Cultured E16.5 mNPC (Day 28)
F
mNPSR-v2
-FGF-2
-Y27632
-LIF
-A83-01
-SB202190
-CHIR99021
-DAPT
-LDN193189
-BMP7
G
Lhx1
Pax8
H
DBA
PAX2
DAPI
LTL
AQP1
PAX8
PODXL
LTL
CDH1
PODXL
WT1
LTL
I
D2
D4
D5
D6
D7
BF
Hoxb7-Venus
J
PODXL
KRT8
LTL
WT1
K
Day 5
L
Day 3
Day 5
BF
GFP
M
PODXL
LTL
DAPI

**Figure S1. Development of mNPSR-v2 medium and characterization of mNPCs cultured in mNPSR-v2 medium. Related to Figure 1.**

(A) Schematic of the experimental procedure for the development of mNPSR-v2 medium.

(B) Bright-field (BF) and immunofluorescence images of primary mNPCs after 4 days of culture in media containing different concentrations of CHIR99021, LDN193189, and SB202190 as indicated, on top of mNPSR medium. Scale bars, 50 μm (bright field) and 100 μm (fluorescence).

(C) Bright-field images of primary mNPCs cultured in mNPSR-v2 medium with or without DAPT for 4 days. Lower panels show enlarged pictures of the boxed areas in the upper panels, highlighting the differentiated morphology from the condition without DAPT. Scale bars, 100 μm (upper panels), 50 μm (lower panels).

(D) qRT-PCR analysis of mNPCs cultured for 4 days in mNPSR-v2 medium with or without DAPT for NPC marker genes as indicated.

(E) qRT-PCR analysis of two long-term cultured mNPC lines, derived from NPCs at two developmental stages (E12.5 and E16.5), for NPC marker gene expression as indicated. Primary E12.5 mNPC sample was used as control.

(F) Bright-field images of established mNPC line upon withdrawal of each individual mNPSR-v2 medium component as indicated for 2 days. mNPCs cultured in complete mNPSR-v2 medium were used as control. Scale bars, 50 μm.

(G) qRT-PCR of mNPCs cultured in different media as shown in (F), for *Lhx1* and *Pax8*, two marker genes representing early nephron induction.

(H) Whole-mount immunofluorescence analyses of nephron organoids derived from E13.5 NPC line (Day 28 of culture) for various marker genes representing glomerulus (PODXL and WT1), proximal tubule (LTL, PAX2, PAX8), loop of Henle (AQP1, PAX2, PAX8), and distal tubule (DBA, PAX2 and PAX8). The nephron organoids were generated using chemically-defined medium protocol. Scale bars, 100 μm.

(I) Time-course bright-field (BF) and *Hoxb7*-Venus fluorescence images of an engineered kidney, reconstructed from cultured mNPCs and cultured mouse UB (labeled with *Hoxb7*-Venus) and cultured on Transwell air-liquid interface, from day 2 (D2) through day 7 (D7). Scale bars, 200 μm.

(J) Whole-mount immunofluorescence analysis of the D7 engineered kidney in (I) for various nephron marker genes as indicated. UB-derived structures were labeled with KRT8. Scale bar, 100 μm.

(K) Chick chorioallantoic membrane (CAM) assay showing host chicken vasculature invasion into cultured mNPCs, 5 days after transplantation. Scale bars, 1 cm.

(L) Bright-field (BF) and GFP images of the mNPC transplant in CAM assay, showing the gradual formation of vasculature around the *Cas9*-GFP mNPCs from 3 days (D3) to 5 days (D5) upon transplantation. Scale bars, 200 μm.

(M) Whole-mount immunofluorescence images showing the formation of nephron structures 5 days after mNPCs were transplanted to CAM. Scale bar, 100 μm.

Data are presented as mean ± SD. Each column represents counts from three biological replicates (n=3). The significance was determined by two-tailed unpaired Student's t tests; ns, not significant; *, $p<0.05$; **, $p<0.01$; ***, $p<0.001$.

### Figure S2

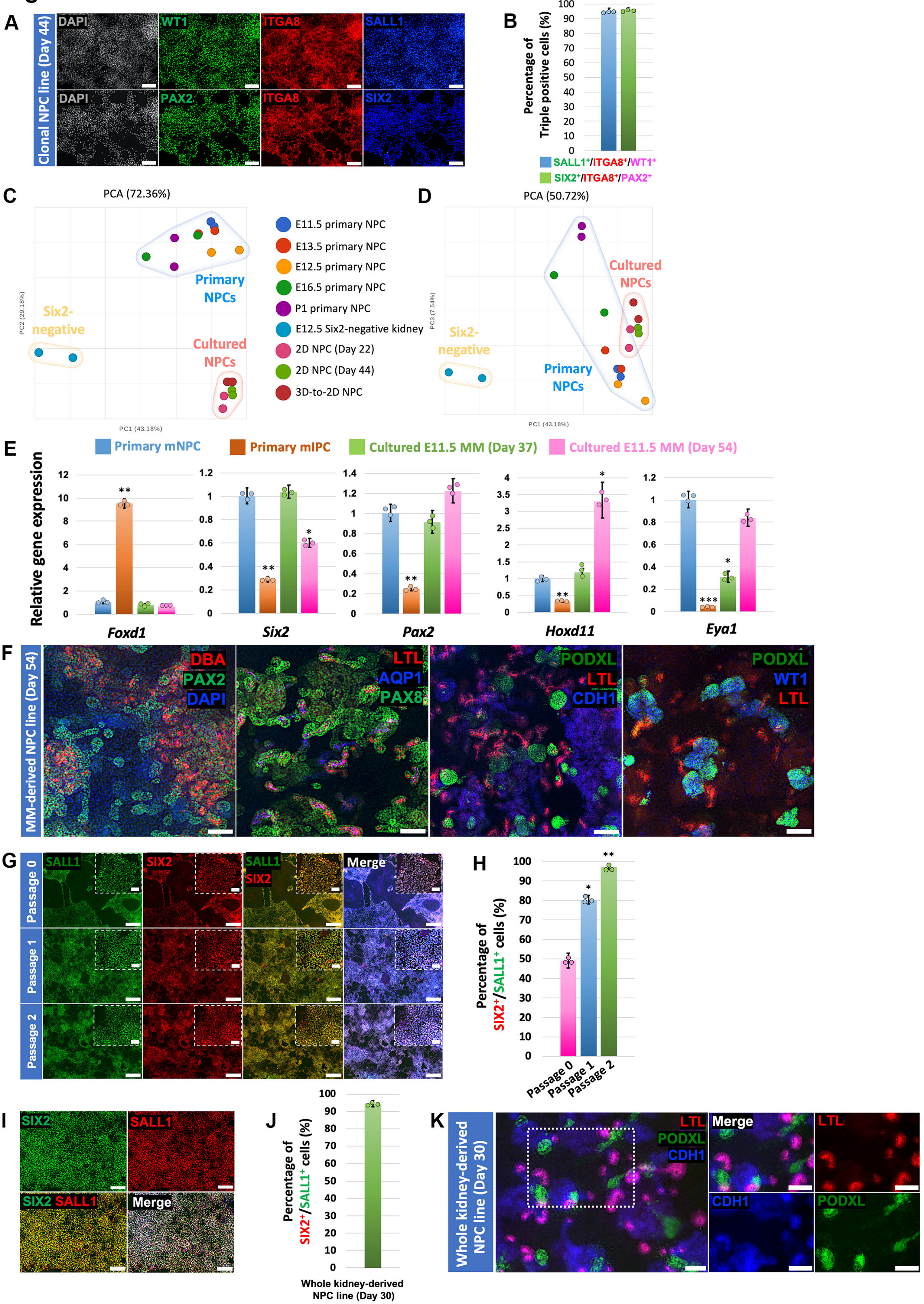

A
Clonal NPC line (Day 44)
DAPI
WT1
ITGA8
SALL1
PAX2
SIX2
B
Percentage of Triple positive cells (%)
SALL1+/ITGA8+/WT1+
SIX2+/ITGA8+/PAX2+
C
PCA (72.36%)
Primary NPCs
Six2-negative
Cultured NPCs
PC1 (43.18%)
PC2 (29.18%)
E11.5 primary NPC
E13.5 primary NPC
E12.5 primary NPC
E16.5 primary NPC
P1 primary NPC
E12.5 Six2-negative kidney
2D NPC (Day 22)
2D NPC (Day 44)
3D-to-2D NPC
D
PCA (50.72%)
PC1 (43.18%)
PC3 (7.54%)
E
Primary mNPC
Primary mIPC
Cultured E11.5 MM (Day 37)
Cultured E11.5 MM (Day 54)
Relative gene expression
Foxd1
Six2
Pax2
Hoxd11
Eya1
F
MM-derived NPC line (Day 54)
DBA
PAX2
DAPI
LTL
AQP1
PAX8
PODXL
CDH1
WT1
G
Passage 0
Passage 1
Passage 2
SALL1
SIX2
Merge
H
Percentage of SIX2+/SALL1+ cells (%)
I
J
Whole kidney-derived NPC line (Day 30)
K
Whole kidney-derived NPC line (Day 30)

**Figure S2. Derivation of mNPC Lines from E11.5 MM and whole embryonic kidneys. Related to Figure 1.**

(A and B) Immunofluorescence images (A) and quantification (B) of a representative clonal mNPC line derived from parental mNPCs cultured for 44 days in mNPSR-v2, for various NPC marker gene expression as indicated. Scale bars, 100 μm.

(C and D) Two-dimensional presentation of principal component analysis (PCA) of bulk RNA-seq data of various primary NPCs isolated from different developmental stages, NPCs cultured in mNPSR-v2 medium, and a negative control *Six2*-negative population from E12.5 kidney.

(E) qRT-PCR analyses of long-term cultured E11.5 MM-derived mNPC lines (on day 37 and day 54) for NPC marker genes *Six2*, *Pax2*, *Hoxd11*, *Eya1*, and IPC marker gene *Foxd1*. Primary E12.5 mNPCs, and primary *Foxd1*$^+$ mIPCs (freshly FACS purified from E13.5 kidneys of *Foxd1*-GFP reporter mice) were used as controls. Note the lack of *Foxd1* expression in the MM-derived mNPC lines, suggesting the selective expansion of NPCs, but not IPCs, from the mNPSR-v2 medium.

(F) Whole-mount immunofluorescence analyses of nephron organoids derived from E11.5 MM-derived NPC line (Day 54 of culture) for various marker genes representing glomerulus (PODXL and WT1), proximal tubule (LTL, PAX2, PAX8), loop of Henle (AQP1, PAX2, PAX8), and distal tubule (DBA, PAX2, and PAX8). The nephron organoids were generated using chemically-defined medium protocol. Scale bars, 100 μm.

(G and H) Immunofluorescence analyses (G) and quantification (H) of E13.5 whole kidney cells cultured in mNPSR-v2 medium for the first three passages for various NPC marker genes as indicated. Scale bars, 100 μm.

(I and J) Immunofluorescence analyses (I) and quantification (J) of E13.5 whole kidney cell-derived mNPC line, after 30 days of culture in mNPSR-v2 medium, for NPC marker genes as indicated. Scale bars, 100 μm.

(K) Whole-mount immunofluorescence analyses of nephron organoids derived from E13.5 whole kidney-derived NPC line (Day 30 of culture) for various nephron marker genes as indicated. The nephron organoids were generated using chemically-defined medium protocol. Scale bars, 100 μm.

Data are presented as mean ± SD. Each column represents counts from three biological replicates (n=3). The significance was determined by two-tailed unpaired Student’s t tests; ns, not significant; *, $p<0.05$; **, $p<0.01$; ***, $p<0.001$.

#### Figure S3

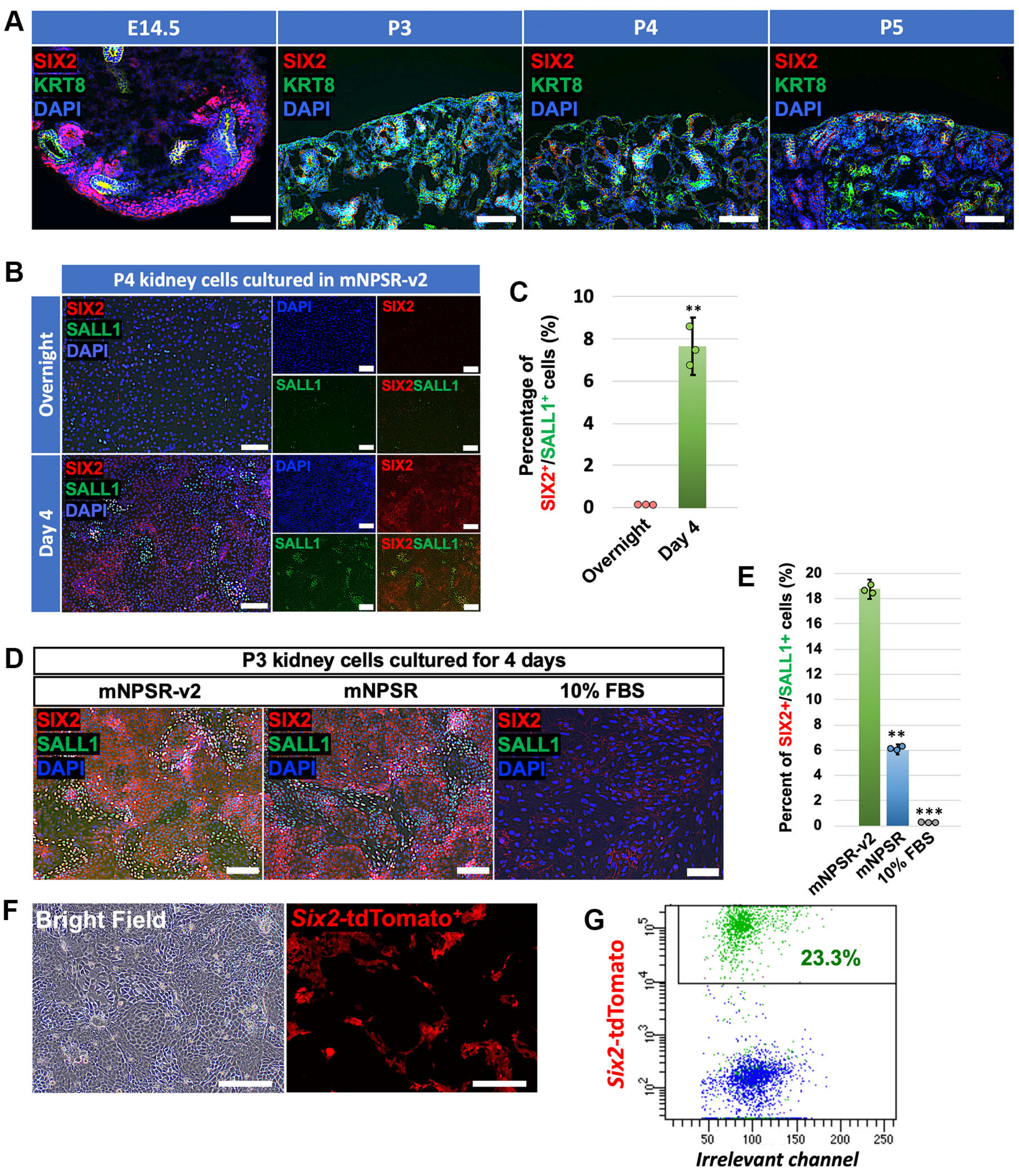

A
E14.5
P3
P4
P5
SIX2
KRT8
DAPI
B
P4 kidney cells cultured in mNPSR-v2
Overnight
Day 4
SIX2
SALL1
DAPI
SIX2SALL1
C
Percentage of SIX2+/SALL1+ cells (%)
10
8
6
4
2
0
**
Overnight
Day 4
D
P3 kidney cells cultured for 4 days
mNPSR-v2
mNPSR
10% FBS
E
Percent of SIX2+/SALL1+ cells (%)
20
18
16
14
12
10
8
6
4
2
0
**
***
mNPSR-v2
mNPSR
10% FBS
F
Bright Field
Six2-tdTomato+
G
Six2-tdTomato
23.3%
50
100
150
200
250
Irrelevant channel

**Figure S3. Plasticity of developing kidney cell types *in vitro*. Related to Figure 2.**

(A) Immunofluorescence images of E14.5, P3, P4, and P5 kidney sections for NPC (SIX2) and UB (KRT8) markers. Note that only E14.5 kidney section showed $SIX2^{+}$ cells (nucleus localization) surrounding the $KRT8^{+}$ UB tips. Membrane-bound SIX2 signals from P3, P4 and P5 kidney sections were from the non-specific binding of the SIX2 primary antibody to some renal tubules. Scale bars, 100 μm.

(B and C) Immunofluorescence images (B) and quantification (C) of P4 whole kidney cells cultured in mNPSR-v2 medium for overnight or 4 days, for NPC marker genes SIX2 and SALL1. Note that only fluorescence signals in the nucleus were true SIX2 signals. Membrane-bound signals were from the non-specific binding of the SIX2 primary antibody. Scale bars, 100 μm.

(D and E) Immunofluorescence staining (D) and quantification (E) of P3 whole kidney cells cultured in mNPSR-v2, mNPSR, and 10% FBS media for 4 days, for various NPC marker genes as indicated. Note that only fluorescence signals in the nucleus were true SIX2 signals. Membrane-bound signals were from the non-specific binding of the SIX2 primary antibody. Scale bars, 100 μm.

(F) Bright-field and *Six2*-tdTomato fluorescence images of P3 whole kidney cells cultured in mNPSR-v2 for 4 days following experimental procedures described in Fig. 2E. Scale bars, 100 μm.

(G) Flow cytometry analysis of P3 whole kidney cells cultured in mNPSR-v2 for 4 days following experimental procedures described in Fig. 2E.

Data are presented as mean ± SD. Each column represents counts from three biological replicates (n=3). The significance was determined by two-tailed unpaired Student's t tests; ns, not significant; *, $p<0.05$; **, $p<0.01$; ***, $p<0.001$.

### Figure S4

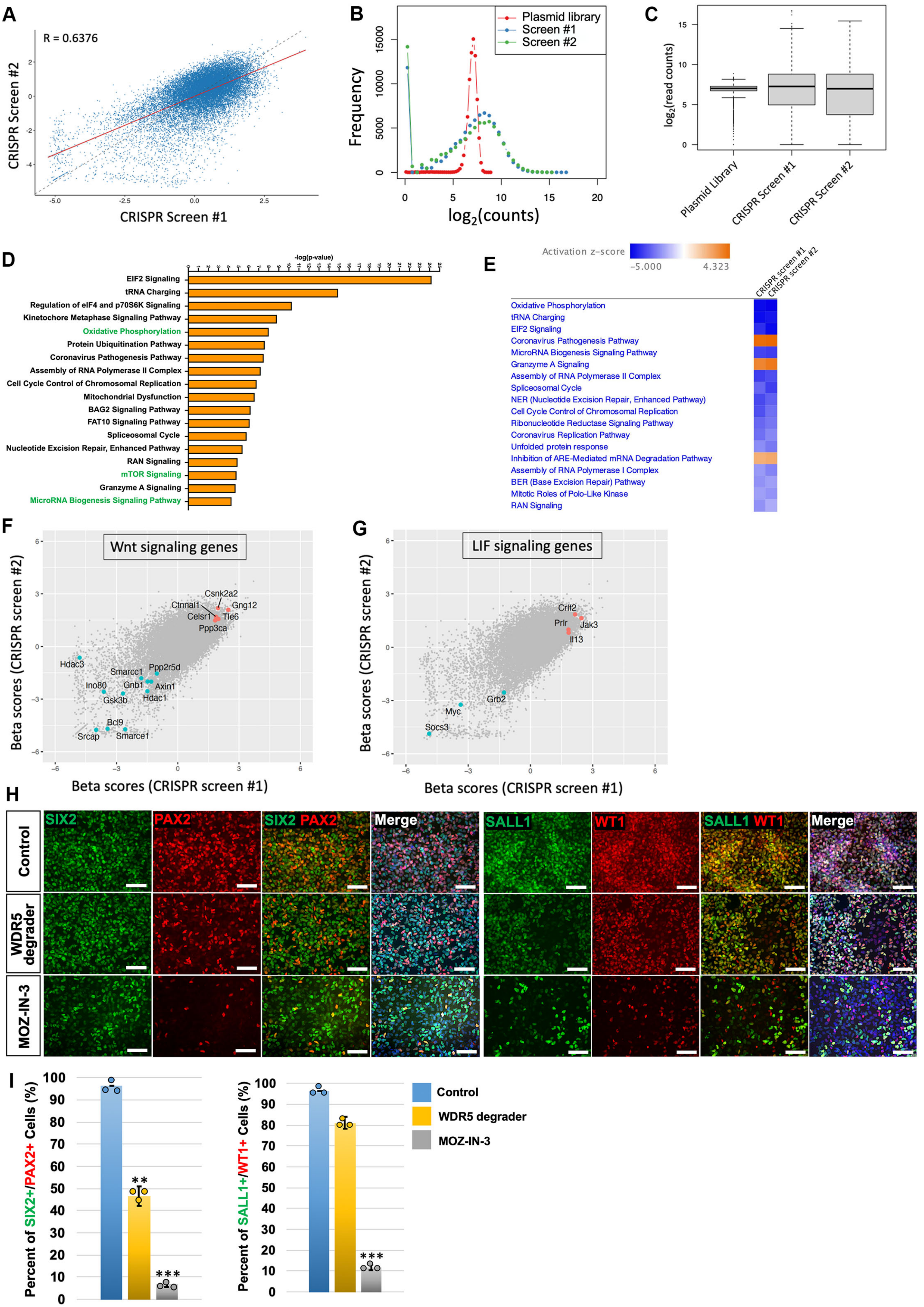
A
R = 0.6376
CRISPR Screen #2
CRISPR Screen #1
B
Plasmid library
Screen #1
Screen #2
Frequency
log2(counts)
C
log2(read counts)
Plasmid Library
CRISPR Screen #1
CRISPR Screen #2
D
-log(p-value)
EIF2 Signaling
tRNA Charging
Regulation of eIF4 and p70S6K Signaling
Kinetochore Metaphase Signaling Pathway
Oxidative Phosphorylation
Protein Ubiquitination Pathway
Coronavirus Pathogenesis Pathway
Assembly of RNA Polymerase II Complex
Cell Cycle Control of Chromosomal Replication
Mitochondrial Dysfunction
BAG2 Signaling Pathway
FAT10 Signaling Pathway
Spliceosomal Cycle
Nucleotide Excision Repair, Enhanced Pathway
RAN Signaling
mTOR Signaling
Granzyme A Signaling
MicroRNA Biogenesis Signaling Pathway
E
Activation z-score
-5.000
4.323
CRISPR screen #1
CRISPR screen #2
Oxidative Phosphorylation
tRNA Charging
EIF2 Signaling
Coronavirus Pathogenesis Pathway
MicroRNA Biogenesis Signaling Pathway
Granzyme A Signaling
Assembly of RNA Polymerase II Complex
Spliceosomal Cycle
NER (Nucleotide Excision Repair, Enhanced Pathway)
Cell Cycle Control of Chromosomal Replication
Ribonucleotide Reductase Signaling Pathway
Coronavirus Replication Pathway
Unfolded protein response
Inhibition of ARE-Mediated mRNA Degradation Pathway
Assembly of RNA Polymerase I Complex
BER (Base Excision Repair) Pathway
Mitotic Roles of Polo-Like Kinase
RAN Signaling
F
Wnt signaling genes
Csnk2a2
Ctnnal1
Gng12
Celsr1
Tle6
Ppp3ca
Hdac3
Smarcc1
Ppp2r5d
Ino80
Gnb1
Axin1
Gsk3b
Hdac1
Bcl9
Srcap
Smarce1
Beta scores (CRISPR screen #2)
Beta scores (CRISPR screen #1)
G
LIF signaling genes
Crlf2
Prlr
Jak3
Il13
Grb2
Myc
Socs3
Beta scores (CRISPR screen #2)
Beta scores (CRISPR screen #1)
H
Control
WDR5 degrader
MOZ-IN-3
SIX2
PAX2
SIX2 PAX2
Merge
SALL1
WT1
SALL1 WT1
Merge
I
Percent of SIX2+/PAX2+ Cells (%)
Percent of SALL1+/WT1+ Cells (%)
**
***
***
Control
WDR5 degrader
MOZ-IN-3

**Figure S4. Genome-wide CRISPR screen in mouse NPC lines. Related to Figure 3.**

(A) MAGeCKFlute scatterplot of beta scores from two biological replicates of mNPC lines on all genes. Grey line means y=x, and red line is the fitted line.

(B) Distribution of normalized sgRNA read counts in the plasmid library, and in two CRISPR screen samples.

(C) Box plot showing the distribution of normalized sgRNA read counts in the CRISPR screen plasmid library, and in two CRISPR screen samples. Boxes, 25th to 75th percentiles; whiskers, 1st to 99th percentiles.

(D) Top 18 enriched ingenuity pathway analysis (IPA) canonical pathways from CRISPR screen replicate #2. Pathways shown in green color are well-established pathways in NPC self-renewal.

(E) Common enriched IPA canonical pathways identified from CRISPR screen #1 and #2.

(F and G) MAGeCKFlute scatterplots of beta scores from two CRISPR screen replicates showing Wnt (F) and LIF (G) signaling related genes.

(H) Immunofluorescence analyses of the expression of NPC marker genes SIX2, PAX2, SALL1, and WT1 in mNPCs after treated with KMT2A inhibitor WDR5 degrader, or KAT6A inhibitor MOZ-IN-3, for 8 days. Scale bars, 50 μm.

(I) Quantification of images in (H).

Data are presented as mean ± SD. Each column represents counts from three biological replicates (n=3). The significance was determined by two-tailed unpaired Student's t tests; ns, not significant; *, $p<0.05$; **, $p<0.01$; ***, $p<0.001$.

### Figure S5

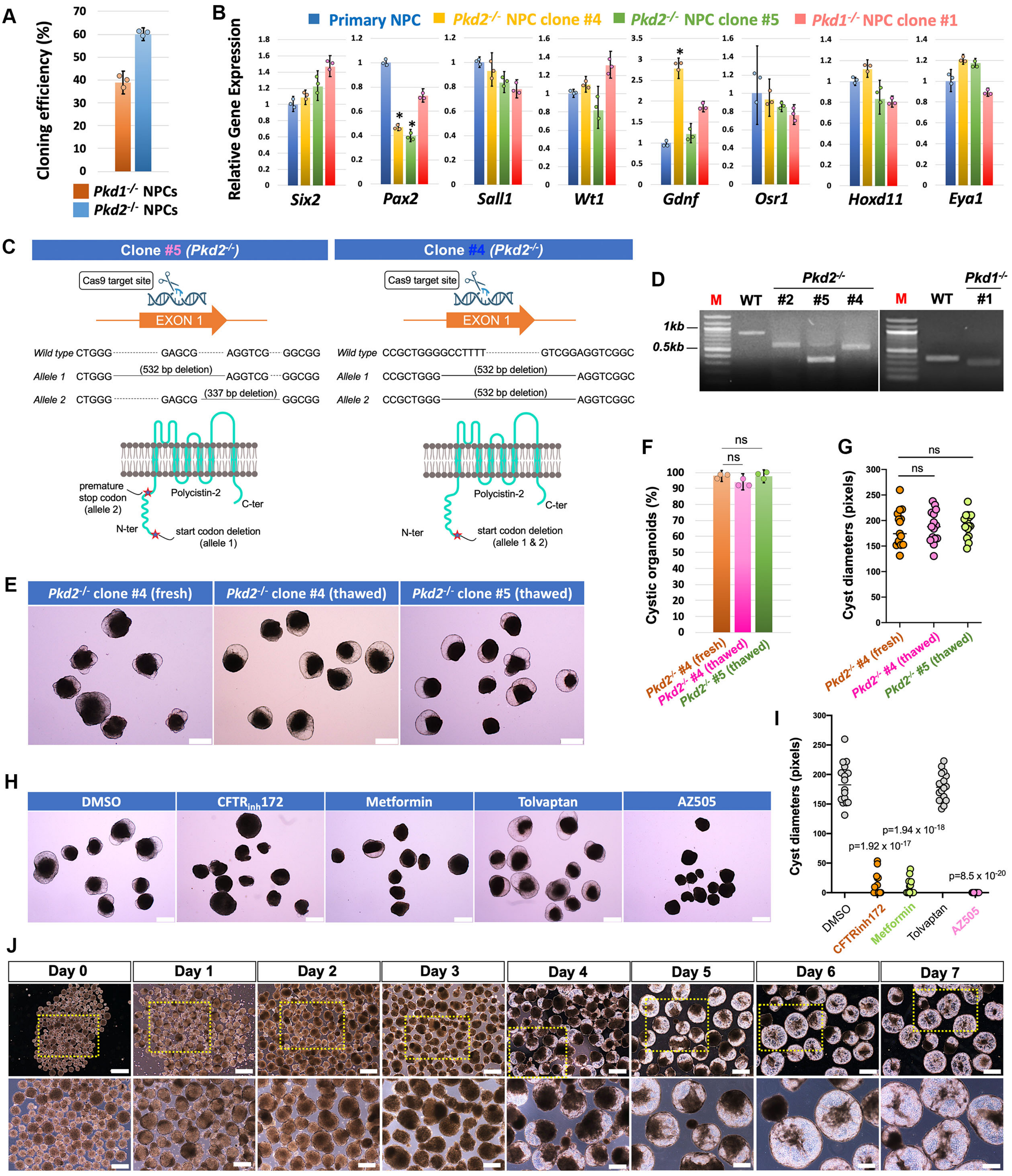

A
Cloning efficiency (%)
Pkd1-/- NPCs
Pkd2-/- NPCs
B
Relative Gene Expression
Primary NPC
Pkd2-/- NPC clone #4
Pkd2-/- NPC clone #5
Pkd1-/- NPC clone #1
Six2
Pax2
Sall1
Wt1
Gdnf
Osr1
Hoxd11
Eya1
C
Clone #5 (Pkd2-/-)
Clone #4 (Pkd2-/-)
Cas9 target site
EXON 1
Wild type CTGGG GAGCG AGGTCG GGCGG
Allele 1 CTGGG (532 bp deletion) AGGTCG GGCGG
Allele 2 CTGGG GAGCG (337 bp deletion) GGCGG
Wild type CCGCTGGGGCCTTTT GTCGGAGGTCGGC
Allele 1 CCGCTGGG (532 bp deletion) AGGTCGGC
Allele 2 CCGCTGGG (532 bp deletion) AGGTCGGC
premature stop codon (allele 2)
Polycistin-2
C-ter
N-ter
start codon deletion (allele 1)
start codon deletion (allele 1 & 2)
D
Pkd2-/-
Pkd1-/-
M WT #2 #5 #4
M WT #1
1kb
0.5kb
E
Pkd2-/- clone #4 (fresh)
Pkd2-/- clone #4 (thawed)
Pkd2-/- clone #5 (thawed)
F
Cystic organoids (%)
ns
G
Cyst diameters (pixels)
Pkd2-/- #4 (fresh)
Pkd2-/- #4 (thawed)
Pkd2-/- #5 (thawed)
H
DMSO
CFTRinh172
Metformin
Tolvaptan
AZ505
I
Cyst diameters (pixels)
p=1.92 x 10-17
p=1.94 x 10-18
p=8.5 x 10-20
J
Day 0
Day 1
Day 2
Day 3
Day 4
Day 5
Day 6
Day 7

**Figure S5. PKD modeling from *Pkd1*$^{-/-}$ or *Pkd2*$^{-/-}$ clonal mNPC line-derived nephron organoids. Related to Figure 4.**

(A) Cloning efficiency of deriving *Pkd1*$^{-/-}$ or *Pkd2*$^{-/-}$ clonal mNPC lines from bulk *Pkd1*$^{-/-}$ or *Pkd2*$^{-/-}$ mNPCs.

(B) qRT-PCR analyses of long-term cultured (26 days) *Pkd1*$^{-/-}$ and *Pkd2*$^{-/-}$ clonal mNPC lines for various NPC marker genes as indicated. Primary NPC sample was used as control.

(C) Schematic showing the multiplexed CRISPR/Cas9-mediated deletions on the first exon of *Pkd2* gene, leading to double allele premature termination of *Pkd2* transcription and thus successful gene knockout of *Pkd2*, in *Pkd2*$^{-/-}$ clonal mNPC lines #4 and #5.

(D) PCR-based genotyping for *Pkd1*$^{-/-}$ or *Pkd2*$^{-/-}$ clonal mNPC lines. Wild-type (WT) mNPC line without genome editing was used as control.

(E) Bright-field images of PKD organoids derived from fresh *Pkd2*$^{-/-}$ nephron organoids, or *Pkd2*$^{-/-}$ nephron organoids after freeze-thaw process, following experimental procedure as described in Figure 4G. Scale bars, 200 μm.

(F and G) Quantification of cystic organoid percentages (F) and cyst diameters (G) for samples shown in (E).

(H) Bright-field images of *Pkd2*$^{-/-}$ clonal mNPC line #4 -derived nephron organoids treated with DMSO (control), or various previously reported PKD drugs: $CFTR_{inh}172$, metformin, tolvaptan, and AZ505. Scale bars, 200 μm.

(I) Quantification of the cyst diameters in samples shown in (H).

(J) Time-course bright-field images showing the development of cystic mini nephron organoids from the *Pkd2*$^{-/-}$ mNPC clonal line, following experimental procedures described in Figure 4K. Enlarged pictures of the boxed areas in the upper panels are shown in the lower panels. Scale bars, 500 μm (upper panels) and 200 μm (lower panels).

Data are presented as mean ± SD. Each column represents counts from three biological replicates (n=3). The significance was determined by two-tailed unpaired Student's t tests; ns, not significant; *, $p<0.05$; **, $p<0.01$; ***, $p<0.001$.

### Figure S6

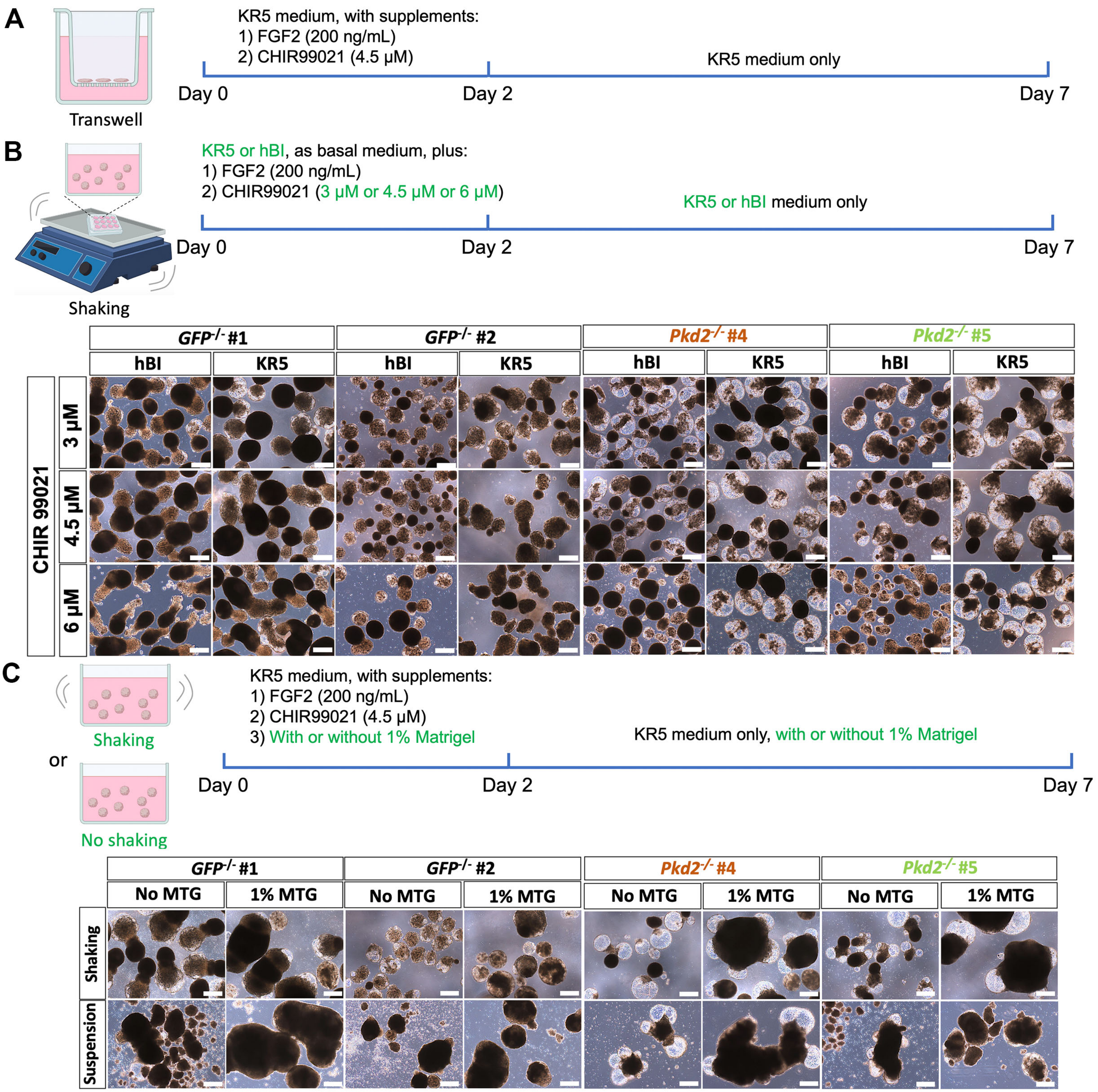


D

| | *GFP$^{-/-}$* #1 | | *GFP$^{-/-}$* #2 | | *Pkd2$^{-/-}$* #4 | | *Pkd2$^{-/-}$* #5 | |
|---|---|---|---|---|---|---|---|---|
| | hBI | KR5 | hBI | KR5 | hBI | KR5 | hBI | KR5 |
| CHIR (3 μM) | Some small non-typical cysts formation | Some small non-typical cysts formation | Some small non-typical cysts formation | Some small non-typical cysts formation | Typical cyst formation | Typical cyst formation | Typical cyst formation | Typical cyst formation |
| CHIR (4.5 μM) | No cyst formation | No cyst formation | No cyst formation | No cyst formation | Typical cyst formation | Typical cyst formation | Typical cyst formation | Typical cyst formation |
| CHIR (6 μM) | Irregular organoid formation | Irregular organoid formation | No cyst formation | No cyst formation | Irregular organoid formation | Typical cyst formation | Irregular organoid formation | Typical cyst formation |

| | *GFP$^{-/-}$* #1 | | *GFP$^{-/-}$* #2 | | *Pkd2$^{-/-}$* #4 | | *Pkd2$^{-/-}$* #5 | |
|---|---|---|---|---|---|---|---|---|
| | Shaking | Suspension | Shaking | Suspension | Shaking | Suspension | Shaking | Suspension |
| No MTG | Some small non-typical cysts formation | No cyst formation; Organoids unhealthy or dead; Organoids fusion | No cyst formation | No cyst formation; Organoids unhealthy or dead; Organoids fusion | Typical cyst formation | Typical cyst formation; Organoids fusion | Typical cyst formation | No cyst formation; Organoids unhealthy or dead; Organoids fusion |
| 1% MTG | Some small non-typical cysts formation; Organoids fusion | No cyst formation; Organoids unhealthy or dead; Organoids fusion | No cyst formation; Organoids fusion | No cyst formation; Organoids unhealthy or dead; Organoids fusion | Typical cyst formation; Organoids fusion | Typical cyst formation; Organoids fusion | Typical cyst formation; Organoids fusion | No cyst formation; Organoids unhealthy or dead; Organoids fusion |

- ● Typical cyst formation
- ◣ No cyst formation
- ▲ Some small non-typical cysts formation
- ◆ Organoids unhealthy or dead
- ⬟ Irregular organoid formation
- ■ Organoids fusion

**Figure S6. Optimization of clonal *Pkd2*$^{-/-}$ mNPC-derived mini PKD organoid model. Related to Figure 4.**

(A) Schematic of the chemically-defined differentiation protocol to generate nephron organoid from cultured mNPCs on Transwell air-liquid-interface.

(B) Upper panel shows the schematic of the experimental design for optimizing cystic PKD organoid generation via manipulation of basal medium and different doses of CHIR99021. Green color indicates the variables tested. Lower panel shows the results of the experiments, in which bright-field images of mini nephron organoids derived from two clonal *GFP*$^{-/-}$ mNPC lines (#1 and #2) or two *Pkd2*$^{-/-}$ mNPC lines (#4 and #5) after 5 days of differentiation in shaking culture are shown. Two different basal media, hBI or KR5 (see also Methods for the detailed medium components), were tested. Under each basal medium, different concentrations of CHIR99021 (3 μM, 4.5 μM, 6 μM) were added together with 200 ng/ml FGF2, for 2 days, followed by continuous culture in the same basal medium for another 3 days. Note that the results indicated that KR5 is superior to hBI in supporting cyst development while different concentrations of CHIR99021 did not have dramatic differences in KR5 basal medium. Scale bars, 200 μm.

(C) Upper panel shows the schematic of the experimental design for further optimizing cystic PKD organoid generation, based on the selected best culture condition identified in (B), in which KR5 medium with 200 ng/ml FGF2 and 4.5 μM CHIR99021 was added for the first 2 days, followed by KR5 medium only for another 3 days. *Pkd2*$^{-/-}$ mini nephron organoids were generated in this culture condition with or without the addition of 1% Matrigel (MTG), and under shaking culture (120 rpm) or static suspension culture. Lower panel shows the results of the experiments. Note that shaking culture without 1% MTG generated the best cystic organoids with limited organoid fusion and was selected as the finalized protocol. Scale bars, 200 μm.

(D) Summary of observations in (B), upper panel, and (C), lower panel, in terms of cyst formation, organoid fusion, and health status of the *GFP*$^{-/-}$ or *Pkd2*$^{-/-}$ mini nephron organoids generated in different conditions tested.

### Figure S7

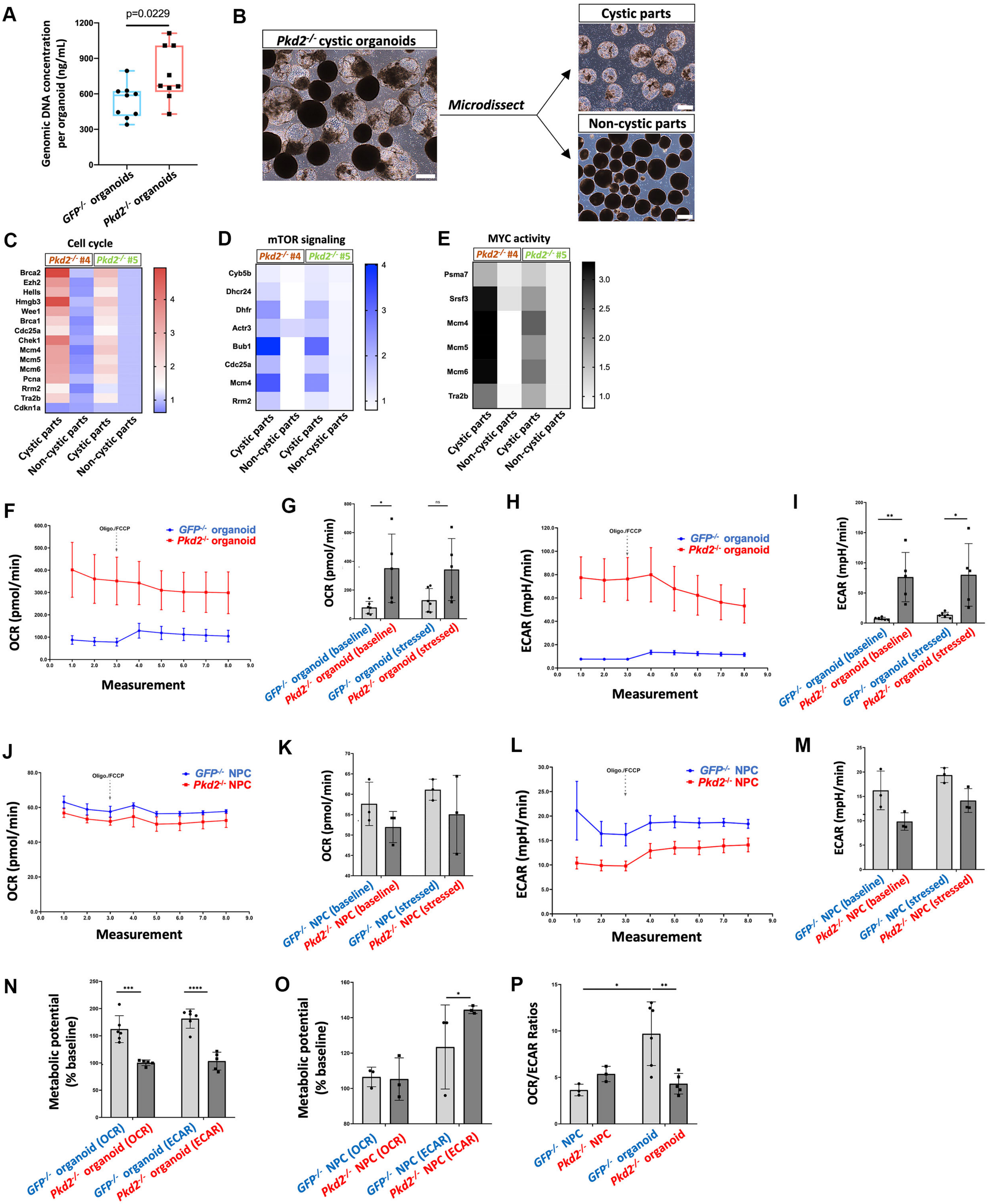
A
p=0.0229
Genomic DNA concentration per organoid (ng/mL)
GFP-/- organoids
Pkd2-/- organoids
B
Pkd2-/- cystic organoids
Microdissect
Cystic parts
Non-cystic parts
C
Cell cycle
Pkd2-/- #4
Pkd2-/- #5
Brca2
Ezh2
Hells
Hmgb3
Wee1
Brca1
Cdc25a
Chek1
Mcm4
Mcm5
Mcm6
Pcna
Rrm2
Tra2b
Cdkn1a
Cystic parts
Non-cystic parts
D
mTOR signaling
Cyb5b
Dhcr24
Dhfr
Actr3
Bub1
Cdc25a
Mcm4
Rrm2
E
MYC activity
Psma7
Srsf3
Mcm4
Mcm5
Mcm6
Tra2b
F
Oligo./FCCP
GFP-/- organoid
Pkd2-/- organoid
OCR (pmol/min)
Measurement
G
OCR (pmol/min)
GFP-/- organoid (baseline)
Pkd2-/- organoid (baseline)
GFP-/- organoid (stressed)
Pkd2-/- organoid (stressed)
H
ECAR (mpH/min)
I
J
GFP-/- NPC
Pkd2-/- NPC
K
GFP-/- NPC (baseline)
Pkd2-/- NPC (baseline)
GFP-/- NPC (stressed)
Pkd2-/- NPC (stressed)
L
M
N
Metabolic potential (% baseline)
GFP-/- organoid (OCR)
Pkd2-/- organoid (OCR)
GFP-/- organoid (ECAR)
Pkd2-/- organoid (ECAR)
O
GFP-/- NPC (OCR)
Pkd2-/- NPC (OCR)
GFP-/- NPC (ECAR)
Pkd2-/- NPC (ECAR)
P
OCR/ECAR Ratios
GFP-/- NPC
Pkd2-/- NPC
GFP-/- organoid
Pkd2-/- organoid

**Figure S7. *Pkd2$^{-/-}$* mini PKD organoids recapitulate molecular, cellular, and metabolic features of PKD. Related to Figure 5.**

(A) Genomic DNA concentrations from single mini nephron organoids derived from *EGFP$^{-/-}$* or *Pkd2$^{-/-}$* mNPCs. The same number of *EGFP$^{-/-}$* or *Pkd2$^{-/-}$* mNPCs were seeded to form organoids. Note the significantly higher genomic DNA concentrations from the *Pkd2$^{-/-}$* organoids, suggesting higher cell proliferation in the cystic organoids.

(B) Bright-field images showing the separation of cystic and non-cystic parts from the *Pkd2$^{-/-}$* organoids through microdissection. Scale bars, 200 μm.

(C–E) Heat map presentation of cell cycle (C), mTOR signaling (D), and MYC activity (E) associated gene expression levels as determined by qRT-PCR, in cystic parts and non-cystic parts of *Pkd2$^{-/-}$* mini PKD organoids.

(F–I) Measurement of oxygen consumption rate (OCR, F and G) and extracellular acidification rate (ECAR, H and I) in *GFP$^{-/-}$* or *Pkd2$^{-/-}$* mini nephron organoids under baseline or oligo/FCCP treatment-induced stressed state using Seahorse XFp assays. F and H show experimental tracings, and G and I show summary data.

(J–M) Measurement of OCR (J and K) and ECAR (L and M) in *GFP$^{-/-}$* or *Pkd2$^{-/-}$* NPCs under baseline or oligo/FCCP treatment-induced stressed state using Seahorse XFp assays. J and L show experimental tracings, and K and M show summary data.

(N and O) Quantification of metabolic potential of *GFP$^{-/-}$* or *Pkd2$^{-/-}$* mini nephron organoids (N), or *GFP$^{-/-}$* or *Pkd2$^{-/-}$* NPCs (O), under baseline or oligo/FCCP treatment-induced stressed state.

(P) Quantification of baseline OCR/ECAR ratios in *GFP$^{-/-}$* or *Pkd2$^{-/-}$* mini nephron organoids, and in *GFP$^{-/-}$* or *Pkd2$^{-/-}$* NPCs.

Data are presented as mean ± SD. Each column represents counts from three biological replicates (n=3). The significance was determined by two-tailed unpaired Student's t tests; ns, not significant; *, $p<0.05$; **, $p<0.01$; ***, $p<0.001$.

### Figure S8

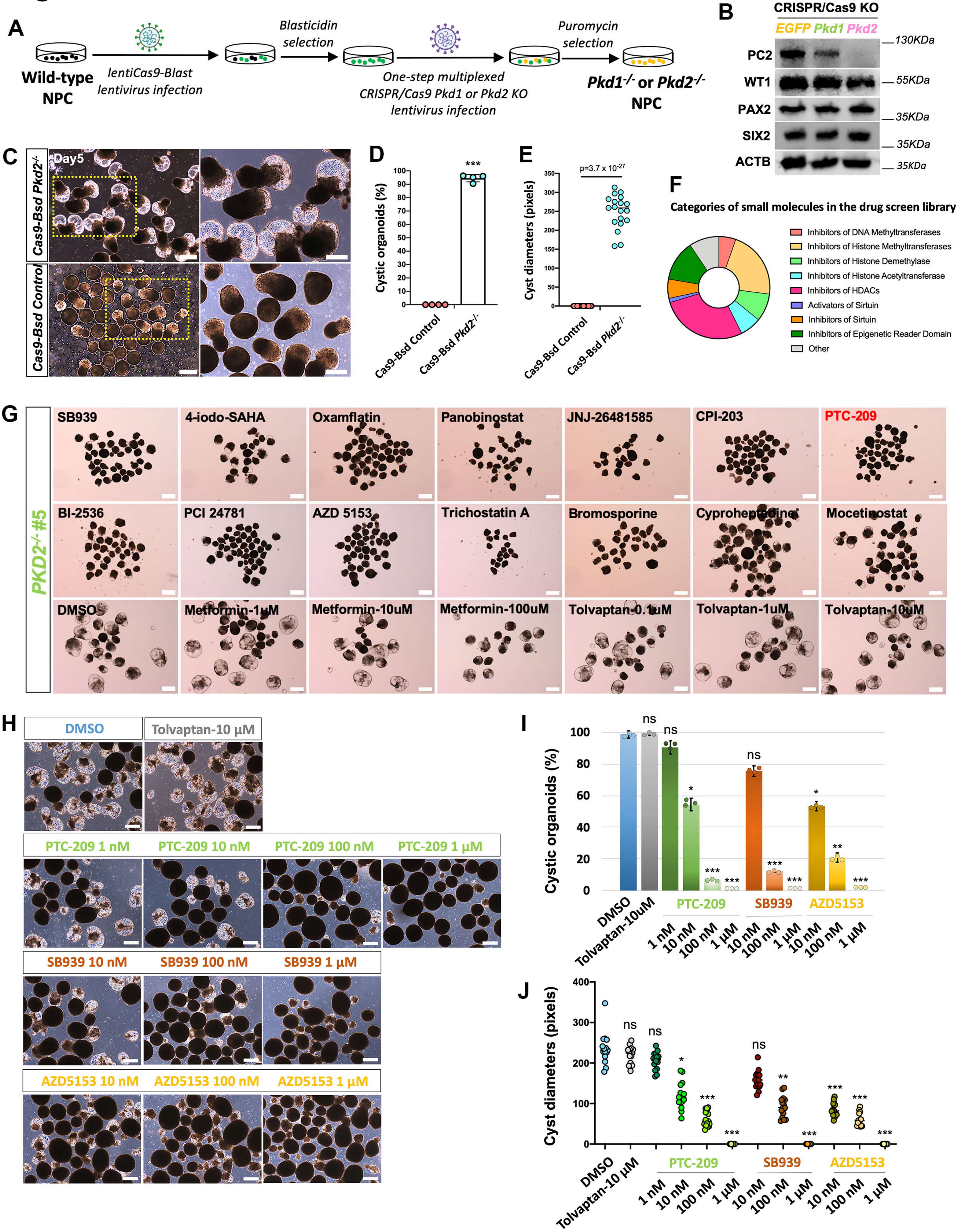

A
Wild-type NPC
lentiCas9-Blast lentivirus infection
Blasticidin selection
One-step multiplexed CRISPR/Cas9 Pkd1 or Pkd2 KO lentivirus infection
Puromycin selection
Pkd1-/- or Pkd2-/- NPC
B
CRISPR/Cas9 KO
EGFP Pkd1 Pkd2
PC2
WT1
PAX2
SIX2
ACTB
130KDa
55KDa
35KDa
35KDa
35KDa
C
Day5
Cas9-Bsd Pkd2-/-
Cas9-Bsd Control
D
***
Cystic organoids (%)
Cas9-Bsd Control
Cas9-Bsd Pkd2-/-
E
p=3.7 x 10-27
Cyst diameters (pixels)
Cas9-Bsd Control
Cas9-Bsd Pkd2-/-
F
Categories of small molecules in the drug screen library
Inhibitors of DNA Methyltransferases
Inhibitors of Histone Methyltransferases
Inhibitors of Histone Demethylase
Inhibitors of Histone Acetyltransferase
Inhibitors of HDACs
Activators of Sirtuin
Inhibitors of Sirtuin
Inhibitors of Epigenetic Reader Domain
Other
G
PKD2-/- #5
SB939
4-iodo-SAHA
Oxamflatin
Panobinostat
JNJ-26481585
CPI-203
PTC-209
BI-2536
PCI 24781
AZD 5153
Trichostatin A
Bromosporine
Cyproheptadine
Mocetinostat
DMSO
Metformin-1uM
Metformin-10uM
Metformin-100uM
Tolvaptan-0.1uM
Tolvaptan-1uM
Tolvaptan-10uM
H
DMSO
Tolvaptan-10 μM
PTC-209 1 nM
PTC-209 10 nM
PTC-209 100 nM
PTC-209 1 μM
SB939 10 nM
SB939 100 nM
SB939 1 μM
AZD5153 10 nM
AZD5153 100 nM
AZD5153 1 μM
I
Cystic organoids (%)
ns
ns
ns
*
***
***
***
***
*
**
***
DMSO
Tolvaptan-10uM
PTC-209
SB939
AZD5153
1 nM
10 nM
100 nM
1 μM
J
Cyst diameters (pixels)
ns
ns
*
***
***
ns
**
***
***
***
***
DMSO
Tolvaptan-10 μM
PTC-209
SB939
AZD5153
1 nM
10 nM
100 nM
1 μM

**Figure S8. PKD modeling from any mNPC line and validation of PKD drug candidate PTC-209. Related to Figure 5.**

(A) Schematic of the experimental protocol for deriving *Pkd1*$^{-/-}$ or *Pkd2*$^{-/-}$ mNPC lines from any mNPC lines using stepwise lentiviral expression of Cas9 (lentiCas9-Blast) followed by multiplexed CRISPR/Cas9 KO (described in Figure 4A).

(B) Western blot analysis of *GFP*$^{-/-}$, *Pkd1*$^{-/-}$, and *Pkd2*$^{-/-}$ mNPC lines (day 28 of culture), derived from wildtype mNPCs following protocol described in (A), for the expression of PC2 and NPC marker genes WT1, PAX2, and SIX2. Beta-actin (ACTB) was used as an internal control.

(C) Bright-field images of mini nephron organoids derived from mNPC line expressing Cas9 but not sgRNA (Cas9-Bsd Control), or mNPC line expressing both Cas9 and multiplexed sgRNAs against *Pkd2* gene (Cas9-Bsd *Pkd2*$^{-/-}$). Enlarged pictures of the boxed areas from the left panels were shown on the right panels. Scale bars, 500 μm (left panels) and 200 μm (right panels).

(D and E) Quantification of cystic organoid percentages and cyst diameters for samples in (C).

(F) Pie chart showing the categories of 148 small molecules in the drug screen library targeting major epigenetic regulatory pathways.

(G) Bright-field images of *Pkd2*$^{-/-}$ clonal NPC line #5-derived mini nephron organoids upon treatment with representative PKD drug candidates identified during the drug screening. Treatment with DMSO, and different doses of metformin and tolvaptan, were used as controls. Scale bars, 500 μm.

(H) Bright-field images of *Pkd2*$^{-/-}$ mini nephron organoids treated with different concentrations of PTC-209 (drug candidate), SB939 (representative HDAC inhibitor, positive control) and AZD5153 (representative BRD4 inhibitor, positive control). Scale bars, 200 μm (>30 mini nephron organoids/group).

(I and J) Quantification of cystic organoid percentages (I) and cyst diameters (J) for samples in (H).

Data are presented as mean ± SD. Each column represents counts from three biological replicates (n=3). The significance was determined by two-tailed unpaired Student's t tests; ns, not significant; *, $p<0.05$; **, $p<0.01$; ***, $p<0.001$.

### Figure S9

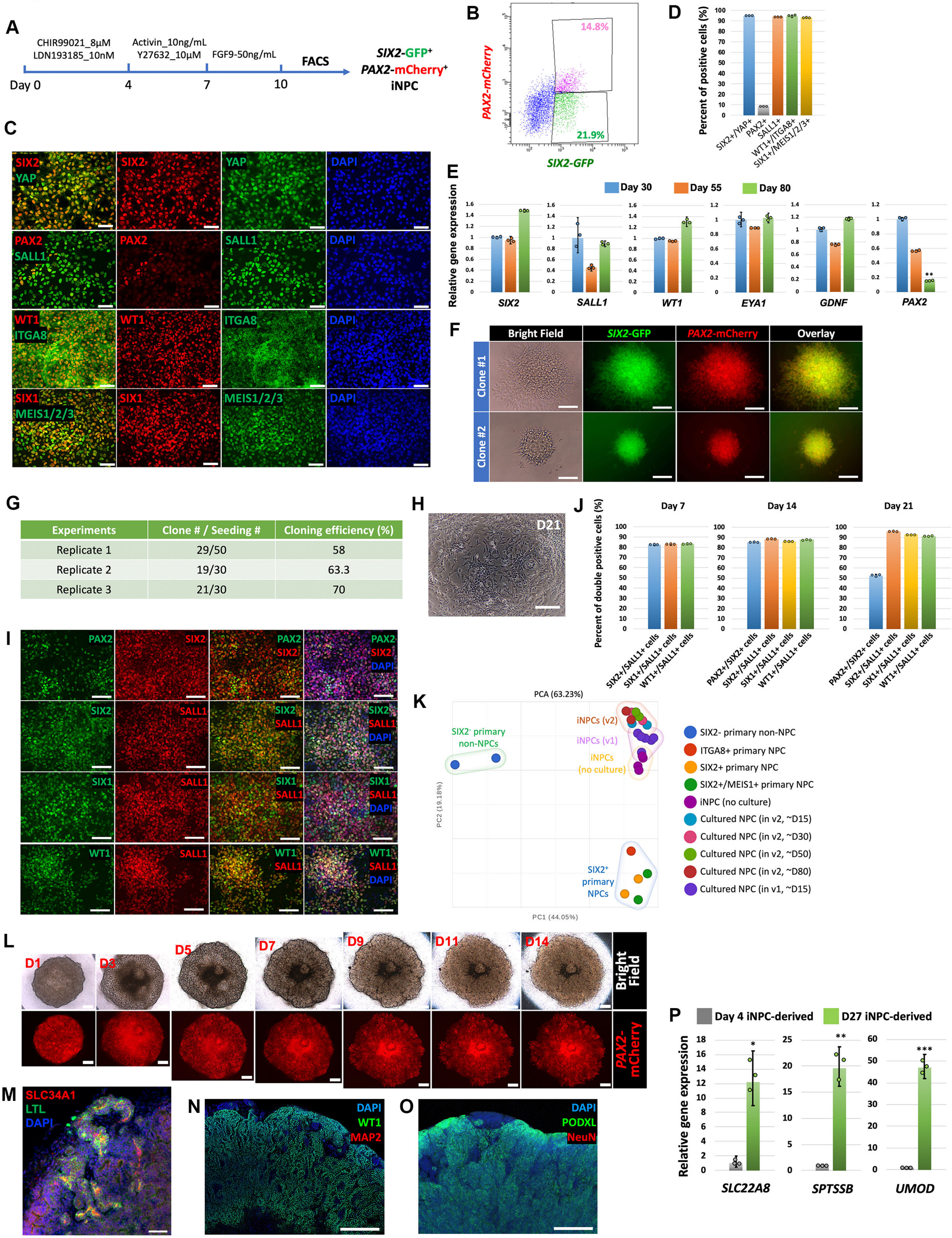


| Experiments | Clone # / Seeding # | Cloning efficiency (%) |
|---|---|---|
| Replicate 1 | 29/50 | 58 |
| Replicate 2 | 19/30 | 63.3 |
| Replicate 3 | 21/30 | 70 |

**Figure S9. Characterization of long-term expandable iNPCs derived from hPSCs. Related to Figure 6.**

(A) Schematic of the directed differentiation protocol to generate iNPCs from *SIX2*-GFP; *PAX2*-mCherry dual-reporter hPSC line. The protocol was adapted from Morizane *et al*., 2015, with minor modification to add ROCK inhibitor Y27632 on Step 2 to enhance cell survival. After 10 days of differentiation, *SIX2*-GFP$^+$/*PAX2*-mCherry$^+$ iNPCs were isolated by fluorescence-activated cell sorting (FACS) then cultured in hNPSR-v2 medium.

(B) Representative flow cytometry plot showing the percentages of *SIX2*-GFP$^+$/*PAX2*-mCherry$^+$ population and *SIX2*-GFP$^+$/*PAX2*-mCherry$^-$ population upon 10 days of differentiation from the dual-reporter hPSC line following the protocol described in (A).

(C and D) Immunofluorescence images (C) and quantification (D) of iNPCs cultured in hNPSR-v2 medium for 87 days, for the expression of YAP and various human NPC marker genes. Scale bars, 50 μm.

(E) qRT-PCR analyses of iNPCs cultured in hNPSR-v2 medium for 30 days, 55 days, and 80 days for expression of various NPC marker genes as indicated.

(F) Bright-field and live-cell fluorescence images showing the expression of *SIX2*-GFP and *PAX2*-mCherry in two representative iNPC clones derived from single cells. Scale bars, 50 μm.

(G) Summary of iNPC cloning efficiency from three independent experiments.

(H) Bright-field image of unpurified hPSC-derived iNPCs (all the cells after 10 days of hPSC differentiation following protocol described in (A), without FACS sorting for *SIX2*-GFP$^+$/*PAX2*-mCherry$^+$ population), upon culture in hNPSR-v2 medium for 21 days. Note that all the cells show highly uniform morphology that is indistinguishable from the cultured FACS-purified iNPCs shown in Figure 6G. Scale bar, 100 μm.

(I) Immunofluorescence images of unpurified iNPCs cultured in hNPSR-v2 medium for 21 days, for various NPC marker genes as indicated. Scale bars, 50 μm.

(J) Quantification of immunofluorescence staining results of unpurified iNPCs cultured in hNPSR-v2 medium for 7 days, 14 days, and 21 days, for various NPC marker genes as indicated.

(K) Two-dimensional PCA plot of bulk RNA-seq data. Different colors represent different primary NPCs, primary non-NPCs, iNPCs without culture, human NPCs cultured in hNPSR-v1 medium for 15 days, or human NPCs cultured in hNPSR-v2 medium for around 15 days, 30 days, 50 days, or 80 days.

(L) Time-course bright-field and live-cell fluorescence images showing the morphological changes and *PAX2*-mCherry reporter expression during the 14-day nephron organoid formation process starting from cultured iNPCs. Note the dramatic morphological changes from day 1 (D1) to day 3 (D3) when numerous tubule-like structures start to form, reflecting mesenchymal-to-epithelial transition, a key early step towards nephron formation. Scale bars, 100 μm.

(M–O) Whole-mount immunofluorescence analyses of human nephron organoids generated from iNPCs cultured in hNPSR-v2 medium for 42 days for proximal tubule marker genes LTL and SLC34A1 (M), and neuronal marker genes MAP2 (N), and NeuN (O). WT1 and PODXL are glomerulus marker genes serving as controls for immunostaining in (N) and (O), respectively. Scale bars, 200 μm.

(P) qRT-PCR analyses of human nephron organoids derived from iNPCs cultured in hNPSR-v2 medium for 4 days or 27 days, for proximal tubule marker *SLC22A8*, and loop of Henle markers *SPTSSB* and *UMOD*.

Data are presented as mean ± SD. Each column represents counts from three biological replicates (n=3). The significance was determined by two-tailed unpaired Student’s t tests; ns, not significant; *, $p<0.05$; **, $p<0.01$; ***, $p<0.001$.

### Figure S10

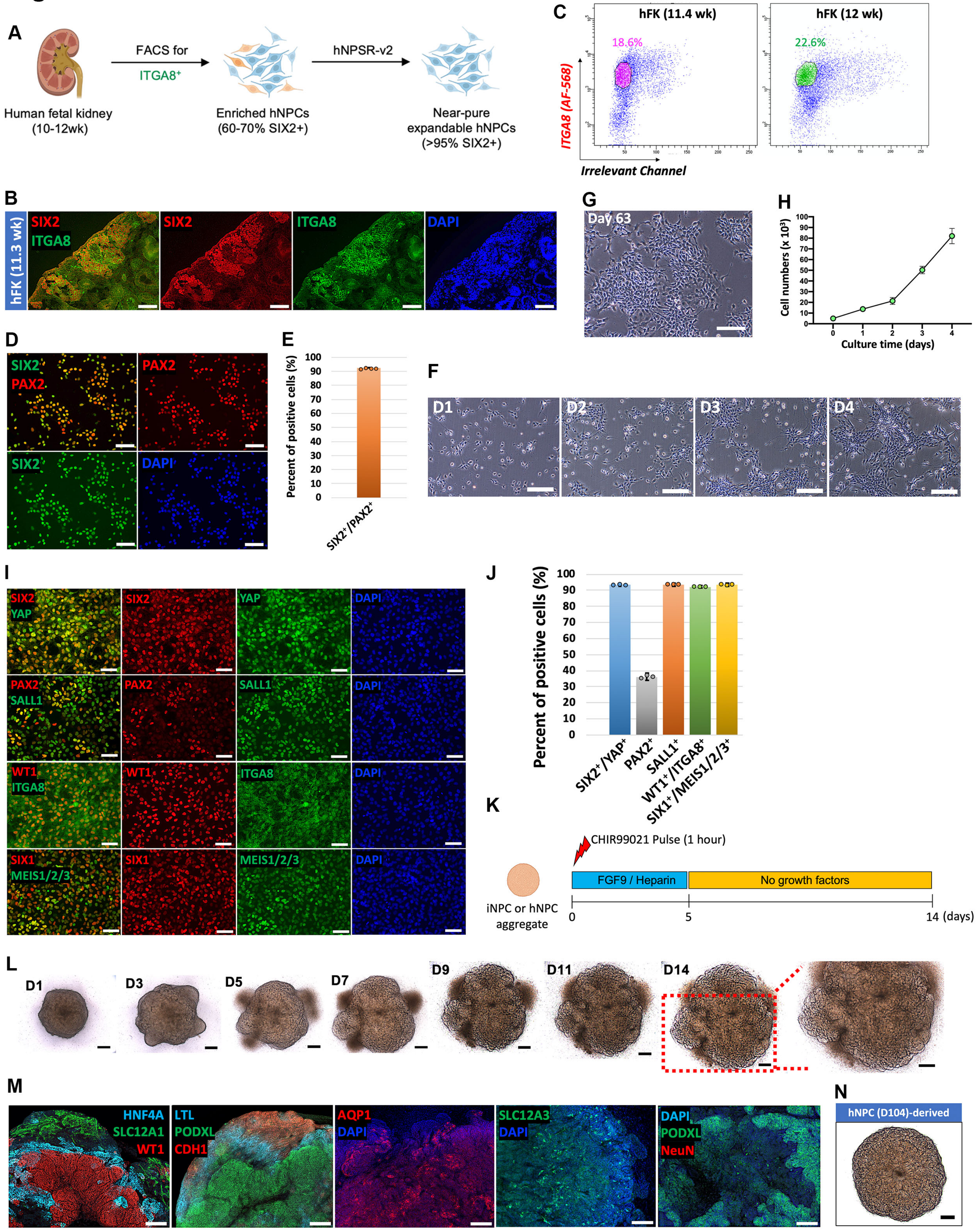

A
Human fetal kidney (10-12wk)
FACS for ITGA8+
Enriched hNPCs (60-70% SIX2+)
hNPSR-v2
Near-pure expandable hNPCs (>95% SIX2+)
B
hFK (11.3 wk)
SIX2
ITGA8
DAPI
C
hFK (11.4 wk)
18.6%
hFK (12 wk)
22.6%
ITGA8 (AF-568)
Irrelevant Channel
D
SIX2
PAX2
DAPI
E
Percent of positive cells (%)
SIX2+/PAX2+
F
D1
D2
D3
D4
G
Day 63
H
Cell numbers (x 10^3)
Culture time (days)
I
SIX2
YAP
PAX2
SALL1
WT1
ITGA8
SIX1
MEIS1/2/3
DAPI
J
Percent of positive cells (%)
SIX2+/YAP+
PAX2+
SALL1+
WT1+/ITGA8+
SIX1+/MEIS1/2/3+
K
CHIR99021 Pulse (1 hour)
FGF9 / Heparin
No growth factors
iNPC or hNPC aggregate
0
5
14 (days)
L
D1
D3
D5
D7
D9
D11
D14
M
HNF4A
SLC12A1
WT1
LTL
PODXL
CDH1
AQP1
DAPI
SLC12A3
NeuN
N
hNPC (D104)-derived

**Figure S10. Derivation of hNPC lines from human fetal kidney tissue. Related to Figure 6.**

(A) Schematic showing the experimental procedures for derivation of long-term expandable hNPC lines from human fetal kidney tissue with hNPSR-v2 medium.

(B) Immunofluorescence images of 11.3-week human fetal kidney sections co-stained for NPC marker genes SIX2 and ITGA8. Scale bars, 100 μm.

(C) Flow cytometry plots showing percentages of $ITGA8^+$ populations (circled) from 11.4-week or 12-week fetal kidneys as two representative examples.

(D and E) Immunofluorescence images (D) and quantification (E) of FACS-enriched primary $ITGA8^+$ hNPCs, after overnight culture in hNPSR-v2 medium, for NPC marker genes SIX2 and PAX2. Scale bars, 100 μm.

(F) Time-course bright-field images of the morphology and growth of representative human fetal kidney-derived hNPCs cultured in hNPSR-v2 medium in a typical passage cycle from day 1 (D1) to day 4 (D4). Scale bars, 100 μm.

(G) Bright-field image of human fetal kidney-derived hNPC line cultured in hNPSR-v2 medium for 63 days. Scale bar, 100 μm.

(H) Growth curve of human fetal kidney-derived hNPC line cultured in hNPSR-v2 medium in a typical 4-day passage cycle starting from 5,000 cells.

(I) Immunofluorescence images of human fetal kidney-derived hNPC line cultured in hNPSR-v2 medium for 87 days for the expression of YAP and various human NPC marker genes. Scale bars, 100 μm.

(J) Quantification of immunostaining results from (I).

(K) Schematic showing the chemically-defined stepwise human nephron organoid induction protocol starting from cultured hNPCs or iNPCs.

(L) Time-course bright-field images showing the induction of human nephron organoid from cultured hNPCs following the stepwise 14-day differentiation protocol as described in (K). Scale bars, 100 μm.

(M) Whole-mount immunofluorescence analyses of human nephron organoids generated from hNPCs cultured in hNPSR-v2 medium for 45 days. Different segments of the nephron were formed, including glomerulus (PODXL, WT1), proximal tubule (HNF4A, LTL), loop of Henle (AQP1, SLC12A1), and distal tubule (CDH1, SLC12A3). Note that no neuronal cells positive for NeuN were observed, suggesting limited off-target cell population in the cultured NPC-derived nephron organoid. Scale bars, 100 μm.

(N) Bright-field image of human nephron organoid generated from hNPCs cultured in hNPSR-v2 medium for 104 days. Scale bar, 200 μm.

Data are presented as mean ± SD. Each column represents counts from three biological replicates (n=3). The significance was determined by two-tailed unpaired Student's t tests; ns, not significant; *, $p<0.05$; **, $p<0.01$; ***, $p<0.001$.
