## Supplementary material for "Modeling kidney development, disease, and plasticity with clonal expandable nephron progenitor cells and nephron organoids": Table S1

**Table S1. Chemicals and growth factors used in the optimization of mNPSR-v2 medium.**

| Targeted signaling pathways | Chemicals and growth factors |
|---|---|
| Wnt signaling pathway | CHIR99021, R-Spondin 1, IWR-1 |
| Hippo/YAP signaling pathway | XMU-MP-1, Verteporfin, Lysophosphatidic acid (LPA), Sphingosine-1-phosphate (S1P), TRULI |
| Notch signaling pathway | DAPT |
| TGF-β and BMP signaling pathway | Activin A, SB431542, A83-01, LDN-193189, BMP4 |
| FGF signaling pathway | FGF1, FGF7, FGF8, FGF9, FGF10, FGF20, PD0325901, SB202190, SP600125, |
| Retinoic acid signaling pathway | All trans-Retinoic Acid, TTNPB |
| LIF signaling pathway | JAKI |
| VEGF signaling pathway | VEGF |
| NF-κB signaling pathway | TNF-alpha |
| EGF signaling pathway | EGF |
| Insulin signaling pathway | IGF-1, IGF-2 |
| HGF signaling pathway | HGF |
| SCF/c-KIT signaling pathway | SCF |
| PDGF signaling pathway | PDGF-BB |
| GDNF signaling pathway | GDNF |
| Others | Forskolin, AICAR |
