## Supplementary material for "Modeling kidney development, disease, and plasticity with clonal expandable nephron progenitor cells and nephron organoids": Table S2

### Table S2. mNPSR-v2 medium recipe

Basal medium: DMEM/F12 (1:1) (1X), Invitrogen, Cat. No. 11330-032.

Supplements:

| Reagent Name | Company | Cat. No. | Final Concentration |
|---|---|---|---|
| GlutaMAX-I (100X) | Invitrogen | 35050-079 | 1X |
| MEM NEAA (100X) | Invitrogen | 11140-050 | 1X |
| 2-Mercaptoethanol (55mM) | Invitrogen | 21985-023 | 0.1 μM |
| Pen Strep (100X) | Invitrogen | 15140-122 | 1X |
| B-27 Supplement (50X), minus vitamin A | Invitrogen | 12587-010 | 1X |
| ITS Liquid Media Supplement (100×) | Sigma | I3146-5ML | 1X |
| BMP-7 | R&D | 354-BP-010 | 50 ng/ml |
| FGF-2 | Peprotech | 100-18B | 200 ng/ml |
| Heparin | Sigma | H3149-100KU | 1 μg/ml |
| Y-27632 | Enzo | ALX-270-333-M025 | 10 μM |
| Mouse LIF | Millipore | ESG1107 | 1000 units/ml |
| LDN193189 | Reagents Direct | 36-F52 | 200 nM |
| A83-01 | STEMGENT | 04-0014 | 50 nM |
| SB202190 | Axon Medchem | 1364 | 5 μM |
| DAPT | Sigma | D5942-5MG | 5 μM |
| CHIR99021 | Reagents Direct | 27-H76 | 1.5 μM |
