## Supplementary material for "Modeling kidney development, disease, and plasticity with clonal expandable nephron progenitor cells and nephron organoids": Table S4

### Table S4. Summary of mNPC lines derived with mNPSR-v2 medium

| **mNPC cell lines** | E13.5, E14.5, E15.5, E16.5, E18.5 *Six2-GFP* mNPC lines | E11.5 MM wild type mNPC lines<br>E11.5 MM *Cas9-GFP* mNPC lines<br>E11.5 MM *Six2-tdT* mNPC lines | E12.5 wild-type whole kidney mNPC lines<br>E13.5, E15.5 *Six2-tdT* whole kidney mNPC lines<br>E12.5 *Wnt4-tdT* whole kidney mNPC lines | E12.5, E14.5, E16.5 *Six2-GFP* clonal mNPC lines<br>E11.5 MM *Cas9-GFP* clonal mNPC lines<br>E11.5 MM *Cas9-GFP* *Pkd1*$^{-/-}$ clonal mNPC lines<br>E11.5 MM *Cas9-GFP* *Pkd2*$^{-/-}$ clonal mNPC lines |
|---|---|---|---|---|
| **Sources** | *Six2-GFP* mouse strain | Metanephric Mesenchyme (MM) | Whole kidney | Single mNPC Cell |
| **Purification method** | FACS sorting | Manual dissection | Enzymatic dissociation | FACS sorting, manual dissection |
| **Derived cell lines number** | >12 | >40 | >15 | >45 |
| **Passage #** | P10-P15 | P10-P31 | P11-P13 | P8-P10 |
| ***In vitro* cultured time (Days)** | 30-45 | 30-93 | 33-39 | 24-30 |
| **Fold of NPC expansion** | $10^{11}$-$10^{20}$ | $10^{14}$-$10^{46}$ | $10^{14}$-$10^{17}$ | $10^{9}$-$10^{11}$ |
| **Average passage cycle (Days)** | 3 | 3 | 3 | 3 |
| **Doublings per day** | 0.6542-0.6666 | 0.6291-0.6417 | 0.6711-0.7031 | 0.5838-0.5937 |
| **# of successful differentiation (nephron organoid induction)** | >60 | >200 | >75 | >250 |
| **Differentiation competency** | 100% | 100% | 100% | 100% |
