## Supplementary material for "Modeling kidney development, disease, and plasticity with clonal expandable nephron progenitor cells and nephron organoids": Table S14

### Table S14. qRT-PCR primers sequences

| qRT-PCR Primers (Mouse) | | |
|---|---|---|
| **Gene Name** | **Forward Primer** | **Reverse Primer** |
| *Gapdh* | CATGGCCTTCCGTGTTCCTA | CCTGCTTCACCACCTTCTTGAT |
| mNPC marker genes | | |
| *Six2* | AGGAAAGGGAGAACAGCGAGAA | GGACTGGACGACGAGTGGT |
| *Wt1* | CCACACCCCTACTGACAGTT | TCACTCTCATACCCTGTGCC |
| *Osr1* | CTGCCCAACCTGTATGGTTT | TGGCACTTTAGAAAAAGAGG |
| *Eya1* | GGACAGGCACCGTACAGCTACC | GTGTGCTGGATACGGCGAGCTG |
| *Hoxd11* | TGGAACGCGAGTTTTTCTTT | TTGCAGACGGTCCCTGTTCA |
| *Pax2* | AAGCCCGGAGTGATTGGTG | CAGGCGAACATAGTCGGGTT |
| *Sall1* | CTCAACATTTCCAATCCGACCC | GGCATCCTTGCTCTTAGTGGG |
| *Gdnf* | TCCAACTGGGGGTCTACGG | GCCACGACATCCCATAACTTCAT |
| Mouse pretubular aggregates (PTA) and renal vesicles (RV) marker genes | | |
| *Pax8* | GGCTCTACCTACTCTATCAA | CTGCTGCTGCTCTGTGAGTC |
| *Lhx1* | CTTCTTCCGATGTTTCGGTA | TCATGCAGGTGAAGCAGTTG |
| Mouse interstitial progenitor cell (IPC) marker gene | | |
| *Foxd1* | TGAGCACGGAGATGTCCGATG | CACCACGTCGATGTCTGTCTC |
| Differentiated nephron marker genes | | |
| *Slc12a1* | TCATTGGCCTGAGGCGTAGTTG | TTTGTGCAAATAGCCGACATAGA |
| *Slc12a3* | ACACGGCAGCACCTTATACAT | GAGGAATGAATGCAGGTCAGC |
| *Aqp1* | AGGCTTCAATTACCCACTGGA | CTTTGGGCCAGAGTAGCGAT |
| Genes associated with cell cycle progression, mTOR signaling, and MYC activity | | |
| *Cdc25a* | ACAGCAGTCTACAGAGAATGGG | GATGAGGTGAAAGGTGTCTTGG |
| *Pcna* | TTTGAGGCACGCCTGATCC | GGAGACGTGAGACGAGTCCAT |
| *Rrm2* | TGGCTGACAAGGAGAACACG | AGGCGCTTTACTTTCCAGCTC |
| *Mcm4* | GAGGAAAGCAGGTCGTCACC | AGGGCTGGAAAACAAGGCATT |
| *Tra2b* | AATCCCGTTCTGCTTCCCG | TCGTGACCTTGTATAATGCCTTC |
| *Mcm5* | CAGAGGCGATTCAAGGAGTTC | CGATCCAGTATTCACCCAGGT |
| *Mcm6* | GCTGTTCCTAGACTTCCTGGA | CAACCAGCGTGTTTCTCTCAG |
| *Bub1* | AGAATGCTCTGTCAGCTCATCT | TGTCTTCACTAACCCACTGCT |
| *Actr3* | ACTGTGGCACGGGATATACAA | GTCACCCACTTTTGCAGACTC |
| *Brca1* | CTGCCGTCCAAATTCAAGAAGT | CTTGTGCTTCCCTGTAGGCT |
| *Chek1* | GTTAAGCCACGAGAATGTAGTGA | GATACTGGATATGGCCTTCCCT |
| *Rrm2* | TGGCTGACAAGGAGAACACG | AGGCGCTTTACTTTCCAGCTC |
| *Brca2* | ATGCCCGTTGAATACAAAAGGA | ACCGTGGGGCTTATACTCAGA |
| *Hells* | TGAGGATGAAAGCTCTTCCACT | ACATTTCCGAACTGGGTCAAAA |
| *Hmgb3* | AGGTGACCCCAAGAAACCAAA | GCAAAATTGACGGGAACCTCTG |
| *Srsf3* | GCGCAGATCCCCAAGAAGG | ATCGGCTACGAGACCTAGAGA |
| *Dhcr24* | CTCTGGGTGCGAGTGAAGG | TTCCCGGACCTGTTTCTGGAT |
| *Cyb5b* | GAGCCCTCCGTCACCTACTA | AGCTTTCAGTTGCATCAGCAC |
| *Dhfr* | CGCTCAGGAACGAGTTCAAGT | TGCCAATTCCGGTTGTTCAATA |
| *Psma7* | AGGAGGCGGTCAAAAAGGG | CATACAGACGTTATCGTCCAAGG |
| *Wee1* | GTCGCCCGTCAAATCACCTT | GAGCCGGAATCAATAACTCGC |

| | | |
|---|---|---|
| *Ezh2* | TGCCTCCTGAATGTACTCCAA | AGGGATGTAGGAAGCAGTCATAC |
| *Cdkn1a* | CCTGGTGATGTCCGACCTG | CCATGAGCGCATCGCAATC |
| **qRT-PCR Primers (Human)** | | |
| hNPC marker genes | | |
| *GAPDH* | GTGGACCTGACCTGCCGTCT | GGAGGAGTGGGTGTCGCTGT |
| *SIX2* | AGGAAAGGGAGAACAACGAGAA | GGGCTGGA TGA TGAGTGGT |
| *PAX2* | CCCAAAGTGGTGGACAAGAT | GAAAGGCTGCTGAACTTTGG |
| *EYA1* | GGACAGGCACCATACAGCTACC | ATGTGCTGGATACGGTGAGCTG |
| *HOXD11* | TGGAACGCGAGTTTTTCTTT | CTGCAGACGGTCTCTGTTCA |
| *OSR1* | CTGCCCAACCTGTATGGTTT | CGGCACTTTGGAGAAAGAAG |
| *SALL1* | CCCCGGTTGCTAACAAAAGC | GAGGTTGTGATCGCTGAGGTA |
| *WT1* | GTGACTTCAAGGACTGTGAACG | CGGGAGAACTTTCGCTGACAA |
| *GDNF* | GGCAGTGCTTCCTAGAAGAGA | AAGACACAACCCCGGTTTTTG |
| Differentiated human nephron marker genes | | |
| *SLC22A8* | CCAGAGTCCATACGCTGGTTG | TCACTTGCGGTGTACTTGGC |
| *SPTSSB* | GCTGTGCTGTTTTAGAGCCCT | CCAGGCGAATGTGGATTGG |
| *UMOD* | CGGCGGCTACTACGTCTAC | TGCCATCTGCCATTATTCGATTT |
